# The function and Evolution of Stegosaur Osteoderms and Hypothesized Sexual Dimorphism in *Hesperosaurus*

**DOI:** 10.1101/2025.04.10.648273

**Authors:** Evan T. Saitta, Vincent Bonhomme, Mitchell Lukens, Daniel Vidal, Nicholas R. Longrich, Dean R. Richmond, Maximilian T. Stockdale

## Abstract

*Stegosaurus* is one of the most iconic organisms in Earth’s history due to its impressive dermal osteoderms. These consisted of throat ossicles, two pairs of posterior tail spikes, and an estimated 18 large plates arranged in two staggered rows along the neck, back, and tail. The function and evolution of these structures in stegosaurs is a quintessential question in dinosaur paleontology. Although sociosexual display has become a popular explanation of plate function, sexual dimorphism in dinosaurs has been a contentious topic. Spikes are meanwhile often described as defensive structures, but we consider an additional intra-sex combat function worthy of further study.

To investigate the questions surrounding stegosaur osteoderms, we reevaluate variation in North American stegosaur osteoderms, with attention to hypothesized sexual dimorphism in *Hesperosaurus*, a close relative of *Stegosaurus* with similar osteoderms. We correct errors in previous analyses, address challenges of the dimorphism hypothesis, and use outline analysis and effect size statistics to provide further statistical support that one sex possibly had larger, wide/broad plates (hypothesized male), while the other sex had smaller, tall/narrow plates (hypothesized female). Stegosaur plates are a difficult case study for sexual variation because the multiple plates borne by a single individual can result in datasets that violate assumptions of independence. Despite appreciable variation from head to tail along an individual, the variation in size and shape seen in *Hesperosaurus* plates is still consistent with the presence of sexual variation as evidenced by both size-independent, two-dimensional outline analysis as well as size-versus-shape regression analysis of principal component data, with high confidence in the latter.

We note pathologies on stegosaur caudal vertebrae, particularly in old adults of wide-morph *Hesperosaurus*, as well as in *Stegosaurus* tail spikes. These pathologies are consistent with not just predator–prey interactions but also intraspecific combat. There currently is no unambiguous indication of dimorphism in stegosaur thagomizers because alternative hypotheses, such as ontogenetic or interspecific variation, are not yet fully tested.

Finally, we present a hypothetical model of stegosaur osteoderm evolution and function based on fossil and extant evidence. Osteoderms adapted for defense in earlier thyreophorans likely became exapted for sexual display (i.e., plates) and intra-sex combat (e.g., tail spikes in stegosaurs, clubs in ankylosaurs). Therefore, sexual selection may have initiated the differentiation of plates from the terminal thagomizer and induced the unique asymmetry of staggered plates in *Stegosaurus* and *Hesperosaurus*. Osteoderm development in hypothesized males could have been driven to physiological upper limits by sexual selection, while the intensity of predation pressure dictated osteoderm development in hypothesized females. Increasing predation pressure might correlate with large body size, open habitats, spine-like plate processes, long tail spikes, and gular ossicles. Within this hypothetical evolutionary model, *Hesperosaurus* would represent a stegosaur taxon with a high magnitude of sexual dimorphism, consistent with its smaller body size, rounder plates, presumed lack of gular ossicles, and niche occupation within a wet, forested, vegetation-rich habitat in the northern Morrison foreland basin.

The question of stegosaur plate function requires us to look more broadly at thyreophoran osteoderm variation and evolution and to consider complex, multifactor models that incorporate many previously proposed hypotheses. We must sort between primary versus secondary functions of plates, spikes, and ossicles and how these functions changed over time.

## INTRODUCTION

The function of stegosaur osteoderms (Galton & Upchurch 2004), primarily the plates and spikes of the North American Morrison Formation taxa *Stegosaurus* and *Hesperosaurus*, has been heavily debated. Various hypotheses for osteoderm function have been proposed over time including: 1) thermoregulation (Farlow *et al*. 1976; de Buffrenil *et al*. 1986; Farlow *et al*. 2010); 2) defense or deterrence from predation – with plates possibly used as a threat display to make the animal appear larger to predators (Carpenter 1998) and/or tail spikes used as offensive weapons against predators (Carpenter *et al*. 2005); and 3) sociosexual display (Carpenter 1998) – either as so-called ‘species recognition’ (Main *et al*. 2005), mutual sexual selection (Hone *et al*. 2012), or typical sexual selection (Saitta 2015). *Hesperosaurus* (Carpenter *et al*. 2001) has at times been synonymized with *Stegosaurus* at the generic level (Maidment *et al*. 2008), although recently the genus was again split (Raven & Maidment 2017).

### 1) Thermoregulation Hypothesis

Large surface areas of highly vascular tissue, as seen in stegosaur plates, undoubtedly impact an organism’s thermoregulation. Note that stegosaurs such as *Stegosaurus* and *Hesperosaurus* have been hypothesized to have had lower growth rates compared to other dinosaurs through bone histology (Redelstorff & Sander 2009). A more recent study using Raman spectroscopy of fossil bone has suggested evolution toward decreased metabolic rates in ornithischians, including *Stegosaurus* (Wiemann *et al*. 2022), but this metabolic proxy likely requires more evidence in light of the selective peak assignments and training dataset, extensive alteration of fossils with organic consolidants in historic collections such as the Yale Peabody Museum (YPM), and spectral artefacts that can occur in such spectra (Alleon *et al*. 2021).

Still, the hypothesis that stegosaur plates evolved primarily under selection for thermoregulation to function as ‘thermal windows’ (Andrade 2015; Tattersall 2016) for heat dissipation or absorption has contrasted with increasing evidence for elevated metabolisms and body temperatures broadly across Dinosauria, at least above those of ‘cold-blooded’ ectotherms (Grady *et al*. 2014) and more likely within the endothermic range (Amiot *et al*. 2006; Pontzer *et al*. 2009; Eagle *et al*. 2011; Benton 2021; Grigg *et al*. 2022). This growing evidence suggests that non-avian dinosaurs generally had more advanced physiological mechanisms by which to thermoregulate other than just sagittal fin– or sail-like structures reminiscent of more primitive groups, such as sprawling-limbed pelycosaurs. Instead, sagittal structures such as sails in *Spinosaurus*, are often proposed to be driven by sociosexual selection for display (Sereno *et al*. 2022; Myhrvold *et al*. 2024). It is unclear why stegosaurs would have required plates primarily for thermoregulation, while other non-avian dinosaurs (e.g., closely related ankylosaurs with similarly extensive osteoderms) would not, especially as large size reduces surface area to volume ratios of the body and thereby mitigates temperature fluctuations (Paladino *et al*. 1990). The evolution of osteoderms within Stegosauria further suggests that mediolaterally flattened osteoderms likely evolved before any significant thermoregulatory ability. For example, *Kentrosaurus*, *Miragaia*/*Dacentrurus*, *Wuerhosaurus*, *Tuojiangosaurus*, and *Yingshanosaurus* have plates of smaller size or, based on their shape, lower surface area/volume ratio than the plates of *Stegosaurus* and *Hesperosaurus*.

However, a rejection of thermoregulatory selection as a primary driver of plate evolution does not preclude their thermodynamic impact or even secondary functions. Even if plates did not evolve explicitly for purposes of thermoregulation, their development will still have an impact on the organism’s thermoregulation. This is especially true if the insulation (Kitchener 1991; Picard *et al*. 1994, 1996, 1999) around these vascular structures is not fully accounted through appropriate thicknesses of overlying corneous β-proteins (Holthaus *et al*. 2018), referred to as ‘keratin’ herein. Furthermore, once evolved for one purpose, plates could have later adopted a secondary function of ‘thermal windows’ so that even an endothermic animal could better thermoregulate. Thermoregulation through ‘thermal windows’ among large endotherms can be seen in the pinnae of elephants by using infrared thermography, especially among larger individuals (Weissenböck *et al*. 2010).

### 2) Defense Hypothesis

The hypothesis that the plates functioned as defensive armor against predation has lost support since it was first proposed more than a century ago (Gilmore 1914). However, a defensive function for the tail spikes of the thagomizer, as well as the gular ossicles in some *Stegosaurus* specimens/taxa, remains commonly proposed (e.g., Carpenter *et al*. 2005; Farke 2014). The plates’ thinness is often invoked as justification that they could not function as armor. However, we note that predators can be risk-averse while hunting and highly selective of prey to avoid injury and optimize success. This is known as the ‘life-dinner principle’ (Dawkins & Krebs 1979; Mukherjee & Heithaus 2013; Heurich *et al*. 2016; Elbroch *et al*. 2017; Humphreys & Ruxton 2020), which states that predators have the option to give up on a hunt, because the consequence of failure is only a continuation of hunger until the next successful hunt. In contrast, prey cannot give up because they face death and, therefore, may be inclined to injure the predator.

Stegosaur plates were thin osteoderms and were likely covered in hard keratin based on direct fossil evidence (Christiansen & Tschopp 2010) and bone correlates (Hieronymus *et al*. 2009), which means it is possible that they were sharply edged in life. They were also tightly packed along the dorsal margin and protruded high above the flesh such that plates may have sufficiently deterred theropod bites from above, especially the cervical plates above the vulnerable, long neck. Beyond functioning directly as armor, plates could also have functioned defensively via display by making the stegosaur appear larger to predators (Carpenter 1998).

Even advanced modern predators with relatively high hunting success rates, such as cougars, alter their killing strategy depending on the type of prey. For example, they avoid horns or antlers and target the throat of large prey, including moose or elk (Murphy *et al*. 2009). Therefore, while it is reasonable to assume that plates, spikes, and gular ossicles served different functions based on their differing morphology, we should be cautious about outright rejecting the possibility of plates functioning as predator deterrents.

### 3) Sociosexual Hypothesis

The hypothesis that stegosaur plates, and many other ‘exaggerated structures’ in extinct animals, functioned for sociosexual signaling has grown in popularity (Isles 2009; Tomkins *et al*. 2010; Hone *et al*. 2016), even in cases where sexual dimorphism is not necessarily assumed (Hone *et al*. 2012). Still, there are disagreements as to the nature of this signaling. Some, for example, have suggested that stegosaur plates and other ‘exaggerated structures’ in dinosaurs were displayed only to allow for so-called ‘species recognition’ (Main *et al*. 2005; Padian & Horner 2011, 2013, 2014). This hypothesis has been heavily criticized for numerous reasons, including the idea that traits must exhibit stark presence-versus-absence expression in order to be considered as a sexually dimorphic, secondary sexual characteristic (see Knell & Sampson 2011; Hone & Naish 2013; Knell *et al*. 2013; Mendelson & Shaw 2013; Borkovic & Russell 2014; Saitta *et al*. 2020).

Instead, sexual selection would more likely be the evolutionary driver for a display functional hypothesis given the overall morphology of and variation in ‘exaggerated structures’, their growth patterns (e.g., stegosaur plate growth is delayed compared to that of the rest of the skeleton [Hayashi *et al*. 2009]), and analogy to similar structures on extant animals (Saitta *et al*. 2020). Saitta (2015) proposed that *Hesperosaurus* plates exhibited sexual dimorphism in size and shape. At this point, a question of the relative roles of mate choice versus intra-sex competition arises. While the plates are often considered for inter-sexual display (e.g., females choosing male mates), the tail spikes are sometimes excluded from a sociosexual hypothesis and viewed as purely defensive structures against predators. As we discuss below, we caution against the outright rejection of sexual selection as a possible driver of tail spike functional evolution.

### A More Complex Answer to The Question of Stegosaur Plate Function?

*Stegosaurus* plates have been previously described in great detail. For example, the description of the highly articulated and complete *Stegosaurus* specimen NHMUK PV R36730 (i.e., “Sophie”, previously “Sarah”) by Maidment *et al*. (2015) is shown here (Fig. 1), with an alternative plate arrangement as proposed by Saitta (2015). Still, it is worth examining the evolution and function of plates, tail spikes, and gular ossicles while considering all possible evolutionary drivers, as others have attempted (e.g., Farlow *et al*. 2010).

**Figure 1.**
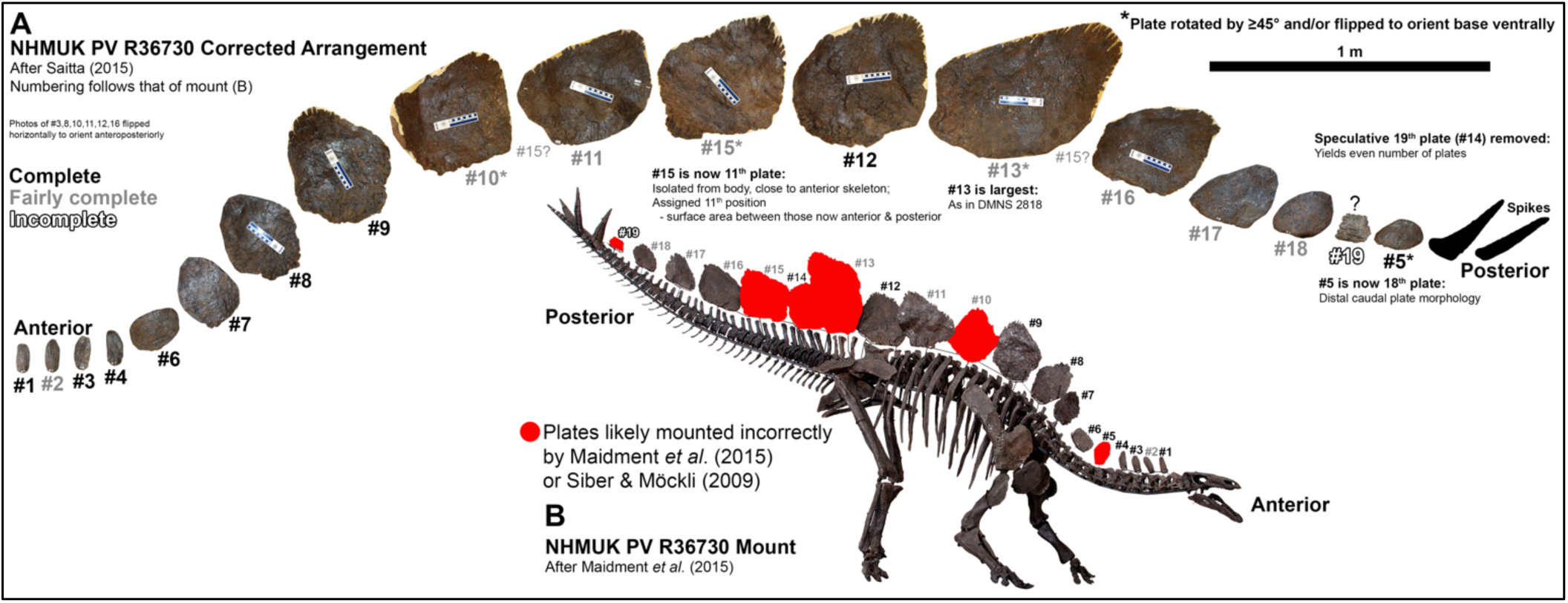
Proposed corrections to NHMUK PV R36730 plate arrangement (Siber & Möckli 2009; Maidment *et al*. 2015) after Saitta (2015). A, plates are arranged from anterior to posterior with the rearrangement mapped onto the original numbering (original number indicated by #) of the mount (spike photographs modified from Siber & Möckli [2009]). All plates are shown with anterior ends to the left, some with rotation and flipping to correctly orient the plates compared to either Siber & Möckli (2009) or Maidment *et al*. (2015), as indicated by an asterisk. Plate numbering font color indicates the level of completeness. Plate #5 was found disarticulated and has the distinctive posterior caudal plate morphology. Plates #11 and #12 were found in close association/possible articulation, so the rearrangement of the disarticulated plate #15 in between them is based solely on its intermediate size and to make the largest plate the 13^th^ position as in DNMS 2818 (largest plate on USNM 4934 may be in 14^th^ position, however). It is also possible that #15 could have come before #11 or after #13 as originally proposed. Likewise, the highly incomplete preservation of #19 makes it exceedingly difficult to reconstruct its position. The speculative plate #14 is removed, for a total of 18 plates. Scale bar correlates to panel A. B, incorrectly mounted plates highlighted onto the fully mounted skeleton of NHMUK PV R36730 figured in Maidment *et al*. (2015), with original plate numbering shown. Anterior is to the right. The original photograph is copyright of The Natural History Museum, while the panel here was downloaded and modified from Figure 1 of Maidment *et al*. (2015) in *PLOS One* whose copyright is as follows: “Copyright: © 2015 Maidment et al. This is an open access article distributed under the terms of the Creative Commons Attribution License, which permits unrestricted use, distribution, and reproduction in any medium, provided the original author and source are credited”.

For example, multiple lizard species in the family Cordylidae show variation in their osteoderms that seemingly cannot be explained by predation alone (e.g., as measured by predator diversity or predator bite force relative to skin toughness) (Broeckhoven *et al*. 2015). Indeed, studies of modern osteoderms highlight the importance of considering more than just defense, while also considering evolutionary tradeoffs. In cordylid lizards, for example, there appears to be a functional trade-off of reduced osteoderm strength with increased thermal capacity as vascularity of the bone increases (Broeckhoven *et al*. 2017a).

## MATERIALS & METHODS

### Institutional Abbreviations

Note that some institutional abbreviations listed below apply only to fossils in the dataset of tail spikes (supplemental material). Furthermore, many Sauriermuseum Aathal (SMA) specimens, including stegosaurs, are scheduled to be donated to the Natural History Museum of the University of Zurich (NMZ) in Zurich, Switzerland. Specifically, *H. mjosi* specimen SMA 0018 (i.e., “Victoria”) will become NMZ 1000018, and *H. mjosi* specimen SMA 0092 (i.e., “Lilly”) will become NMZ 1000092 (Dennis Hansen pers. comm. 2025). Since NMZ is effectively retaining the original SMA numbers, we describe these specimens using their SMA identifiers, in keeping with previous literature. Finally, note that the fossils from the Hayashibara Museum of Natural Sciences (HMNS) were transferred to the Fukui Prefectural Dinosaur Museum (FPDM) in Katsuyama, Japan after the museum’s closure in 2015 (Sonoda & Noda 2016). Specimen HMNS 14, the holotype of *Hesperosaurus*, is now labelled FPDM-V9674. For the time being, we still refer to this holotype as HMNS 14 for consistency with previous literature.

AMNH: American Museum of Natural History, New York, New York, USA

BYU: Brigham Young University Museum, Salt Lake City, Utah, USA

CM: Carnegie Museum of Natural History, Pittsburgh, Pennsylvania, USA

DINO: Dinosaur National Monument, Jensen, Utah, USA

DMNS: Denver Museum of Nature and Science, Denver, Colorado, USA

FPDM: Fukui Prefectural Dinosaur Museum, Katsuyama, Japan

HMNS: Hayashibara Museum of Natural Sciences, Okayama, Japan

JRDI: Judith River Dinosaur Institute, Billings, Montana, USA

MWC: Museum of Western Colorado, Grand Junction, Colorado, USA

NHMUK: Natural History Museum, London, UK

NMZ: Natural History Museum of the University of Zurich, Zurich, Switzerland

SMA: Sauriermuseum Aathal, Aathal, Switzerland

UCRC: University of Chicago Research Collection, Chicago, Illinois, USA

UMNH: Natural History Museum of Utah, Salt Lake City, Utah, USA

USNM: United States National Museum, Washington D.C., USA

VFSMA: Verein für das Sauriermuseum, Aathal, Switzerland

WDC: Wyoming Dinosaur Center, Thermopolis, Wyoming, USA

YPM: Yale Peabody Museum, New Haven, Connecticut, USA

Woodruff *et al*. (2019) claimed that JRDI specimens are “unavailable for study” based on ownership status. This statement is self-evidently untrue: 1) the fossils have been studied (Saitta 2015) and this freely accessible data has been reanalyzed by others since (Mallon 2017; Motani 2021), 2) the fossils have resided within this collection for about two decades, and 3) two authors of Woodruff *et al*. (2019) have accessed and worked on the fossils. Maidment assisted in their excavation, and Woodruff accessed the fossil collection and quarry (JRDI pers. comm. 2024). In fact, several researchers have visited the collection and quarry over the years (e.g., Richmond 2023). Here, we even present novel data (additional measurements/counts and photogrammetry) obtained during a 2023 collection visit, a decade after initial study by Saitta. Prior to its purchase by NHMUK, “Sophie” or NHMUK PV R36730 (previously “Sarah”, SMA RCR0603, SMA DS-RCR-2003-02, or SMA S01) was likewise published on by Maidment *et al*. (2008), Redelstorff & Sander (2009), and Maidment *et al*. (2012) while it was privately owned, moving between two private institutions (SMA and Black Hills Institute of Geological Research), and for sale.

### Approach For This Investigation

Here, we first correct errors in and reanalyze freely available data from the open-access publication of Saitta (2015), which proposed sexual dimorphism in *Hesperosaurus mjosi* plates. We address criticisms and provide novel support for hypothesized sexual dimorphism in *H. mjosi* plates, comparing different statistical approaches based on size and/or shape variables. This dataset contains 40 fairly to highly complete *H. mjosi* plates from four isolated individuals found separately in northern Wyoming with plates in association to their skeletons (seven plates from HMNS 14, two plates from SMA 0018, 11 plates from SMA 0092, six plates from VFSMA 001) as well as plates from two multi-individual, disarticulated bonebeds: the JRDI 5ES Quarry in central Montana containing a minimum of five individuals based on femora and pelvises (nine sufficiently complete plates from an unknown number of individuals) and the WDC DMQ bonebed in southeast Wyoming containing a minimum of two individuals based on pelvises (five plates from an unknown number of individuals).

Analyses include an examination of plate angle distributions compared to those from three highly articulated and complete *Stegosaurus* specimens (DMNS 2818, NHMUK PV R36730, USNM 4934). Next, the 40 fairly to highly complete *H. mjosi* plates underwent two-dimensional outline analysis in the R package MOMOCS where Procrustes-aligned, elliptical-Fourier-transformed plate outlines were studied using linear discriminant analysis (LDA), principal component analysis (PCA), and multivariate analysis of variance (MANOVA) to examine variation across body position and between hypothesized sexual dimorphs. Finally, the divergence of plate shape at larger plate size between hypothesized sexual dimorphs was constrained using PCA of six variables and calculating 95% confidence and prediction intervals to each hypothesized morph. This divergence analysis 1) compared the morphology of the only known juvenile North American stegosaur plate (DMNS 33359) to those of the 40 *H. mjosi* plates, 87.5% of which are evidenced to be sexually mature through histology of the plates and/or associated skeletons (Redelstorff & Sander 2009; Hayashi *et al*. 2012; Saitta 2015), and 2) compared the results between the non-transformed variables of Saitta (2015) and the linearly transformed variables of Motani (2021).

We also examined plate and spike variation and pathology across North American stegosaur taxa. This includes a consideration of plates from uncertain North American stegosaur taxa of uncertain taxonomic status, but with plates that resemble those of *H. mjosi*. Next, we describe pathologies of North American stegosaur vertebrae and tail spikes. Then, we examined the distribution of tail spike lengths from 36 *Stegosaurus* tail spikes using a Gaussian kernel density function.

Finally, we propose a hypothetical model for osteoderm evolution and function. This model requires consideration of multiple functions and physiological effects of osteoderms and the different types of selective pressures acting upon them – selection pressures whose intensities can furthermore vary across time and space. The model is informed by fossil and extant evidence.

### *Hesperosaurus* Data: Hypothesized Sexual Dimorphism

To start this investigation, it is critical to first examine the plates of one taxon of North American stegosaur, *H. mjosi* (Carpenter *et al*. 2001) (Fig. 2). Saitta (2015) published a case for sexual dimorphism addressing a full range of alternate hypotheses on 40 fairly to highly complete plates of *H. mjosi* (at the time, synonymized as *Stegosaurus mjosi*). This hypothesized sexual dimorphism has been both challenged (Mallon 2017) and supported (Motani 2021) in subsequent studies. Given 1) that there is intra-individual variation in osteoderms along the back of a single individual from head to tail, 2) that osteoderms can easily disarticulate from their *in vivo* positions, and 3) that not all specimens are found as highly complete isolated individuals with plates associated/articulated with the rest of the skeleton, it is unsurprising that this question is a challenging one.

**Figure 2.**
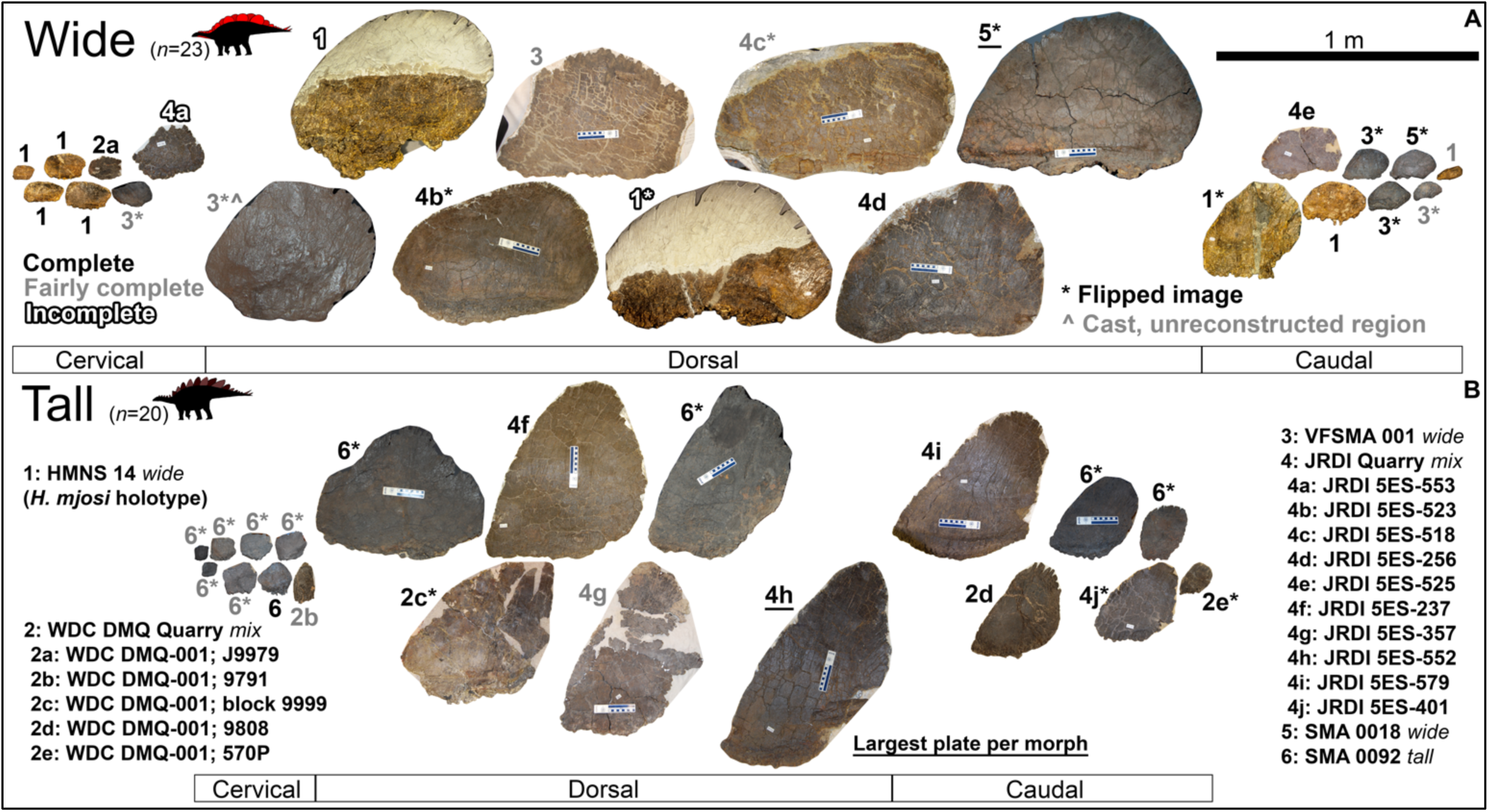
Wide– and tall-morph plates of *H. mjosi* (Saitta 2015). Plates are scaled to each other in size. The anterior edge is to the left, with some images flipped horizontally for consistency (indicated by an asterisk). Plates are organized roughly from estimated anterior to posterior body position. Font color is coded for level completeness. Plates from multi-individual quarries are indicated with lettering after the number. The anterior-most dorsal plate of VFSMA 001 (here, 3*^) is a photo of the cast and only shows the unreconstructed region of the plate without the superfluous apical tip that was added to it for exhibition. The largest plate by surface area of each morph is underlined.

*H. mjosi* was hypothesized (Saitta 2015) to consist of a wide, broad, oval plate morph that reached sizes ∼45% larger in surface area than the tall, narrower, subtriangular plate morph. Saitta (2015) hypothesized that the larger, wider plates could be consistent with males selected for visual display, while the smaller, taller, more pointed plates could be consistent with females who experience selection to develop plates as ‘prickly’ deterrence against predation. At the time, Saitta (2015) possibly presented the most thorough argument for sexual dimorphism in a single non-avian dinosaur taxon (e.g., compared to Larson 2008, Rinehart *et al*. 2009, Raath 1990, Barden & Maidment 2011, Weishampel & Chapman 1990, Chapman *et al*. 1981, Dodson 1976, Smith 1998, Persons *et al*. 2015). Using different lines of evidence from morphometrics, histology, and taphonomy, Saitta (2015) tested multiple alternative hypotheses for the quantitatively observed divergence in plate shape at larger size – with stegosaur plates being an *a priori* plausible sexual display structure.

The 40 fairly to highly complete *H. mjosi* plates studied by Saitta (2015) derived from four isolated individuals and two multi-individual bonebeds (JRDI 5ES, WDC DMQ). The isolated individual SMA 0092 (i.e., “Lilly”) has a total of 11 fairly to highly complete plates. One plate from SMA 0092 was erroneously excluded by Saitta (2015) as an outlier in PCA under the incorrect assumption that it obscured the variation in the remainder of the sample (Motani 2021; see below). The total sample of 40 sufficiently complete *H. mjosi* plates, as well as the unnecessarily reduced PCA dataset of 39 plates in Saitta (2015), represents up to 18 individuals (Fig. 2). A minimum of 8 individuals can also be estimated, if plates from the two multi-individual bonebeds belonged to at least two individuals each, based on the presence of both putative plate morphs in both of these bonebeds. Otherwise, the absolute minimum would be six individuals.

Nine fairly to highly complete plates and one incomplete plate were recovered from the JRDI 5ES Quarry (i.e., JRDI “Quarry 2” in Richmond [2023]) bonebed in central Montana (near Grass Range). These 10 total plates were found alongside a minimum of five individual skeletons ascertained from the number of pelvises (Saitta 2015) and right femora (Table 1). The pelvises and femora in the JRDI 5ES Quarry exhibit a range of sizes and degrees of skeletal fusions. However, histology indicates that all nine fairly to highly complete plates themselves in the JRDI 5ES Quarry come from at least subadult/young adult individuals or older (Saitta 2015). In fact, a total of 35 out of the 40 *H. mjosi* plates in the dataset of Saitta (2015) are known to come from at least subadult/young adult individuals based on histology (Redelstorff & Sander 2009; Hayashi *et al*. 2012).

**Table 1.** *H. mjosi* pelvic and femoral size data for the JRDI 5ES Quarry in central Montana (near Grass Range). Specimens are ordered roughly from largest to smallest. A minimum of five individuals are represented over the range of sizes. Measurements are rounded to the nearest cm.

| Pelvis # | JRDI<br>5ES-424 | JRDI<br>5ES-263 | JRDI<br>5ES-396 | JRDI<br>5ES-494<br>(Right<br>ilium<br>only) | JRDI<br>5ES-311 |  |  |  |  |
| --- | --- | --- | --- | --- | --- | --- | --- | --- | --- |
| # sacral vertebrae | 4 | 4 | 4 | ? | 4 |  |  |  |  |
| # fused presacral<br>vertebrae | 2 | 1 | 1 | ? | 1 |  |  |  |  |
| # total fused sacral<br>vertebrae | 6 | 5 | 5 | ? | 5 |  |  |  |  |
| # free presacral<br>transverse<br>processes pairs | 2 | 1 | 1 | ? | 0 |  |  |  |  |
| Anteroposterior<br>length of all fused<br>centrums (cm) | 46 | 34 | 34 | ? | 32 |  |  |  |  |
| # fused neural<br>spines | 5 | 5 | 4 | ? | 4 |  |  |  |  |
| Width at narrowest<br>point of ilia (cm) | 61 | ? | ~66 | ? | ~27 |  |  |  |  |
| # openings in iliac<br>shelf right side | 4 | 4 | 5 | ? | 7 (+1 above<br>posterior-most<br>prezygapophyses) |  |  |  |  |
| # openings in iliac<br>shelf left side | 4 | 5 | 5 | ? | 7 (+1 above<br>posterior-most<br>prezygapophyses) |  |  |  |  |
| Anteroposterior<br>length of ilium<br>(cm) | ~97 | 94 | ~88 | 84 | ~81 |  |  |  |  |
| Femur # | JRDI<br>5ES-230 | JRDI<br>5ES-501 | JRDI<br>5ES-423 | JRDI<br>5ES-425 | JRDI<br>5ES-474 | JRDI<br>5ES-101 | JRDI<br>5ES-354 | JRDI<br>5ES-371 | JRDI<br>5ES-105<br>(Incomplete<br>femur) |
| Right or Left? | Right | Left | Right | Left | Right | Left | Right | Left | Right |
| Total length (cm) | 94 | 91 | 84 | 84 | 82 | ~81 | 71 | 70 | ? |
| Proximal width<br>(cm) | 25 | 24 | 24 | 20 | 24 | ? | 21 | 20 | ? |
| Mid-shaft width<br>(cm) | 13 | 14 | 12 | 11 | 11 | 10 | 10 | 10 | ? |
| Distal width (cm) | ? | 25 | 22 | 21 | 21 | 21 | 18 | 19 | ? |

Five fairly to highly complete plates are known from the WDC DMQ bonebed (i.e., Meilyn Quarry) near Como Bluff in southeast Wyoming (Schmude & Weege 1996; Foster 2003). These five plates were found alongside a minimum of two individuals based on the number of pelvises. No histological analysis has been performed for the WDC DMQ specimens, so these five plates have uncertain ontogenetic status in the dataset of Saitta (2015).

### Saitta (2015) Dataset Error

Motani (2021) noticed an error in Saitta (2015). Three measurements of a single plate were inadvertently entered into the wrong cells of the data matrix during PCA: 1) ‘width’; 2) distance from the center of the base to the apex; and 3) surface area on both sides of the plate (i.e., the measured surface area from a photograph of one side of the plate doubled). The error was introduced in the Supporting S2 Dataset of Saitta (2015) when transcribing the correct, raw data (S1 Dataset of Saitta [2015]) into a form amenable for analysis in R (S1 Script of Saitta [2015]). The error occurred in the first-mounted dorsal plate of SMA 0092, which caused it to become an extreme outlier in the initial PCA (figure 2A of Saitta [2015]).

Saitta (2015) overlooked the incorrectly entered data for that plate. Saitta (2015) instead attributed the outlier result to the plate’s somewhat unusual morphology – it has a wide base but a well-defined, high-angle, triangular apex consistent with the other 10 tall-morph plates on this isolated individual. However, the outlier result was not due to the plate’s morphology and high angle value, as Saitta (2015) assumed, but rather due to the error in the dataset. Fortuitously, this plate was dropped from the dataset, and the PCA was repeated without using it – mitigating the effect of this typographical error, although it reduced the sample size by one plate (Saitta 2015).

The correct values are as follows (rounded to nearest whole number): ‘width’ (i.e., the major axis of the oval basal portion of the plate) = 63 cm; distance from the center of the base to the apex = 51 cm; and surface area of both sides of the plate (i.e., double the surface area of one photographed side) = 4,647 cm^2^. To avoid any further errors, all reanalysis here used the correct, raw, and unedited data from the S1 Dataset of Saitta (2015), meaning that input values were rounded to the nearest whole number prior to analysis.

Note that there was also a typographical error in the Materials and Methods under the section “Plate Measurements” in Saitta (2015) that initially read, “The shape of the large caudal plate on wide morph specimen HMNS 14 (#1 in S4A Fig) lent itself to have ‘width’ measured in the same manner as those of tall morph plates such that ‘width’ equals the minor axis”. This sentence should instead read: The shape of the large caudal plate on wide-morph specimen HMNS 14 (#1 in S4A Fig) lent itself to have ‘width’ measured *such that ‘width’ equals the anteroposterior width*. This typographical error did not affect the data or analysis.

### Recently Described North American Specimens & Stegosaur Biogeography

Recently described stegosaur specimens from North America may have affinity to *H. mjosi* or implications for stegosaur biogeography, but they are not necessarily amendable to further analysis here. Maidment *et al*. (2018) hypothesized habitat partitioning between *H. mjosi* in the northern Morrison Formation and the lumped *Stegosaurus* taxon in the southern Morrison Formation (as also suggested by Whitlock *et al*. [2018] and Foster [2020]) and a possibly unique northern Morrison biota. However, Maidment *et al*. (2018) did not include a more northerly, isolated individual previously attributed to *Stegosaurus*. This specimen was initially labeled JRDI 155 (i.e., ‘Giffen’). It was found in the early 2000’s near Stockett, in central Montana, and is housed at the Great Plains Dinosaur Museum in Malta, Montana. They did not include this specimen despite 1) that doing so potentially complicates their suggestion of habitat partitioning, 2) that this northernmost specimen has been published before by Maidment *et al*. (2008) and described as *Stegosaurus* therein, and 3) that this specimen is curated at a listed institution of the authorship of Maidment *et al*. (2018).

‘Giffen’ (JRDI 155), now listed as GDPM 178, was instead subsequently described in further detail in Woodruff *et al*. (2019) by two of the same authors as, and shortly after, the prior paper by Maidment *et al*. (2018). They concluded that no diagnostic autapomorphies were present in this specimen, further highlighting the potential issue with the biogeographic analysis of Maidment *et al*. (2018). The taxonomic identity of ‘Giffen’ (JRDI 155/GDPM 178) is currently uncertain, its only prepared plate is incomplete, and three unprepared plates (potentially cervical plates) were not figured in Woodruff *et al*. (2019). Therefore, while it is possible that this specimen is *H. mjosi*, we exclude it from our dataset here.

Woodruff *et al*. (2019) also reported additional forelimb fossils from a likely *H. mjosi* individual from the same central Montana grassy hillside as the JRDI 5ES Quarry (i.e., “Quarry 2”). This specimen, GPDM 205 (i.e., “Gates”), was horizontally separated by less than ∼135 m and of uncertain stratigraphic position (possibly within ∼2 m vertically) from the JRDI 5ES Quarry material included in Saitta (2015) and herein. Furthermore, this material lacks plates, femora, and a pelvis of potential relevance to the study by Saitta (2015). “Gates” may represent an unrelated individual or it may signify an eroded partial individual from the JRDI 5ES Quarry assemblage, and extensive vegetation on the hillside makes stratigraphic correlation difficult. If it is from this same mass death assemblage, then it would presumably increase the minimum individual count of *H. mjosi* to six individuals.

Another specimen was assigned to *H. mjosi* (MOR 9728) by Maidment *et al*. (2018) from the O’Hair Quarry near Livingston, in southern Montana. MOR 9728 was likewise not added to the dataset here because of the incomplete preservation of its plates.

Biogeographic and stratigraphic details (e.g., Maidment 2023) of North American stegosaur diversity deserve to be further investigated as new specimens are uncovered. For now, we find it reasonable to consider *H. mjosi* as a relatively more northern Morrison Formation stegosaur taxon, with confirmed specimens being so far found in only Wyoming and Montana, compared to *Stegosaurus*, which is known from much further south in the formation, especially Utah and Colorado. Note that the specimen of *Stegosaurus* NHMUK PV R36730 found at the Red Canyon Ranch (Maidment *et al*. 2015) is also relevant to these biogeographic questions because it was found in the same area as multiple *H. mjosi* specimens near Shell, in northern Wyoming (SMA 0018 [i.e., “Victoria”] from the Howe-Stephens Quarry; SMA 0092 and VFSMA 001 [i.e., “Moritz”] both from the Howe-Scott Quarry, although at different stratigraphic horizons from each other) (Siber & Möckli 2009). As such, if stratigraphic overlap is also present, it is possible there was some geographic overlap between the two genera in at least Wyoming.

## CHALLENGES TO *HESPEROSAURUS* DIMORPHISM: BOTH PLATE MORPHS LIKELY DO NOT APPEAR ON ONE INDIVIDUAL

Saitta (2015) presented evidence that the variation in plates of *H. mjosi* was not solely explainable by ontogeny. Histological analyses show that both tall and wide morphs could come from fully grown adults and 35 plates in the sample (87.5%) were from at least young adults or older. For example, the tall plates in the JRDI 5ES Quarry tended to be histologically more mature than the wide plates from the same quarry, indicating that a single individual did not likely have both plate morphs. Histological analysis shows that we currently have examples of fully grown, old adults with external fundamental systems (EFS) in their bones for both wide-morph, isolated individuals (SMA 0018 with EFS in the long and girdle bones [Redelstorff & Sander 2009], HMNS 14 with an EFS in the plate [Hayashi *et al*. 2012]) and within tall-morph plates from a multi-individual bonebed (plates JRDI 5ES-357 and JRDI 5ES-552 [Saitta 2015]).

Furthermore, the variation in plates was not solely explainable by interspecific variation, as evidenced by taphonomy and diagnostic characters in the skeleton other than the plates (Saitta 2015). At least two multi-individual bonebeds (JRDI 5ES, WDC DMQ) contained plates of both morphs among a group that lived at the same time and place and could even have been a social group in life (Saitta 2015). However, an alternative hypothesis that the plate variation can be solely explained by intra-individual variation anteroposteriorly along the back is deserving of further investigation.

### Plate Angle & Plate Position

Mallon (2017) concluded that statistically significant bimodality in the second principal component (PC2) of the *H. mjosi* plate PCA data of Saitta (2015) meant that proposed dimorphism instead represented intra-individual variation. The reasoning was that the angle between the center of the plate’s base and its apex heavily loads PC2, and this angle decreases posteriorly along the back of an individual stegosaur (figure S6 in Saitta [2015]). However, the bimodality of plate angles in the dataset is moot since dimorphism of plate angles was not proposed by Saitta (2015). Instead, plate shape (as described by PC3) diverged at larger sizes (as described by PC1). This point was raised by Motani (2021) in response to Mallon (2017). The plate angle has relatively little loading to PC1 or PC3 in the PCA of Saitta (2015).

Regardless, even if intra-individual variation in plates (from small to large plates [i.e., PC1] or from anterior to posterior plates [i.e., PC2] in Saitta [2015]) is large or even larger than potential sexual variation (from wide to tall [i.e., PC3]), this does not preclude the possibility of dramatic sexual dimorphism. As an analogy, even if feather morphology and color in birds can vary greatly within a single individual across its body, this does not preclude sexual dimorphism in plumage. There are also weaknesses in significance tests (Amrhein *et al*. 2019) and in test of bimodality as it relates to questions of dimorphism (Saitta *et al*. 2020).

Furthermore, there are several lines of evidence against both wide and tall plate morphs appearing on a single individual (Saitta 2015). 1) Studied isolated individuals do not show obvious mixtures of plates of both morphs (i.e., HMNS 14, SMA 0018, and VFSMA 001 are wide-morph individuals, while SMA 0092 is a tall-morph individual [Fig. 2]). 2) Both plate morphs possess cervical, dorsal, and caudal plate morphologies (Fig. 2). Note that if the tall-morph plates simply represented low-angle wide-morph plates, then such tall-morph plates would be restricted to the tail, which they do not appear to be (Fig. 2). 3) Both *H. mjosi* plate morphs show similar angle distributions to the bimodal distribution of highly articulated and complete *Stegosaurus* specimens (DMNS 2818, NHMUK PV R36730, and USNM 4934), showing that both morphs represent a reasonably complete plate series from head to tail (Fig. 3, figure S20 of Saitta [2015]). Note how the *H. mjosi* plates show similar patterns (including bimodality) and spread of angle distributions (Fig. 3) as seen in highly articulated and complete *Stegosaurus*, suggesting that the samples from both morphs represent a relatively complete plate series.

**Figure 3.**
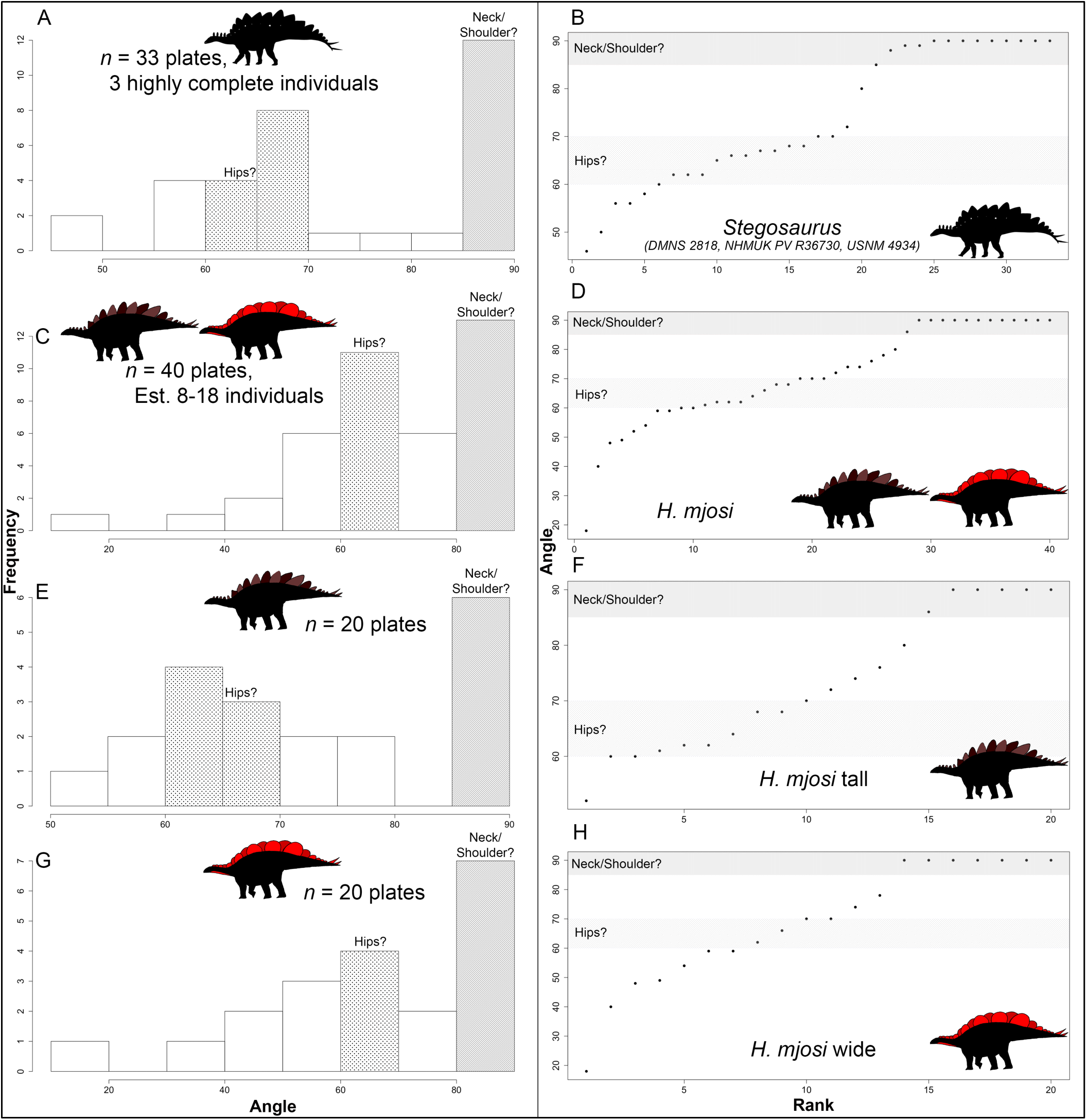
Distributions of angles between the center of the base and apical tip of plates from North American stegosaurs as histograms (A, C, E, G) and rank ordered (B, D, F, H). Plates are fairly to highly complete. Angle values are shaded for 60– 70° and 80–90°, which in highly articulated and complete *Stegosaurus* roughly correspond to the hip and neck/shoulder regions, respectively. A, B, highly articulated and complete *Stegosaurus* specimens. C, D, both morphs of *H. mjosi* combined. E, F, tall-morph *H. mjosi*. G, H, wide-morph *H. mjosi*. Data and *H. mjosi* silhouettes from Saitta (2015).

### Plates on Isolated Individuals

Isolated individuals appear to show only one morph of plate along their backs – HMNS 14, SMA 0018, and VFSMA 001 are wide-morph specimens, while SMA 0092 is a tall-morph specimen (Fig. 2).

The anterior dorsal plate from SMA 0092, an isolated individual with 10 other fairly to highly complete tall-morph cervical, dorsal, and caudal plates (Fig. 2), has a wide basal portion and is more similar to the wide-morph plates than any other assigned tall-morph plate in the dataset. However, it has a pronounced triangular apex. This triangular apex is unlike that of a plate from a similar body position on the isolated wide-morph specimen VFSMA 001. Because it was prepared prior to our current understanding of *H. mjosi* morphology (Siber & Möckli 2009), the VFSMA 001 anterior dorsal plate was casted with a superfluously reconstructed apex added. VFSMA 001 has five other fairly to highly complete wide-morph cervical, dorsal, and caudal plates (Fig. 2).

*Stegosaurus* plates show a dramatic expansion on the lower portion of the plate as they progress from cervical plates above the neck to anterior dorsal plates above the shoulder (e.g., as in DMNS 2818, NHMUK PV R36730 [Fig. 1], and USNM 4934, as well as plates from other specimens [Galton & Upchurch 2004; Saitta 2014]). It is perhaps unsurprising that a *H. mjosi* specimen (SMA 0092) with tall-morph plates, including many cervical plates that differ from wide-morph cervical plates (Fig. 2), shows basal anteroposterior expansion in the anterior dorsal plates. Therefore, neither SMA 0092 nor VFSMA 001 can be said to unambiguously exhibit both plate morphotypes.

## NOVEL ANALYSES OF *HESPEROSAURUS* PLATE DIMORPHISM

### Outline Analyses

One way to approach the hypothesized sexual dimorphism in *H. mjosi* plates is to attempt to quantitatively describe plate shape as precisely as possible. Here, we use the MOMOCS package (Bonhomme *et al*. 2014) outline analysis in R (version 4.1.1) to describe plate shape more precisely than did Saitta (2015), who used six one– and two-dimensional morphometric variables.

Note that a weakness of the analysis is that we only examine shape without considering size. Analyses of shape variation alone without considering ontogeny or sexual dimorphism in both shape and size can fail to recognize sexual dimorphism even in structures of extant taxa commonly thought to be under sexual selection, such as wildebeest horns (Gerstenhaber & Knapp 2022) and cassowary casques (Green *et al*. 2022). This is due to one sex, often adult females, with secondary sexual structures distinct from those of the larger adult males becoming hard to distinguish from younger/smaller males when both sexes undergo similar allometric developmental trajectories.

The MOMOCS methodology follows this analytical pipeline: Procrustes alignment of the x and y coordinates of the plate outlines using the anterior and posterior edges of the base as landmarks, followed by elliptical Fourier transformation, LDA on the elliptical Fourier coefficients trained for either body position of the plate or the morph of the plate, PCA on the elliptical Fourier coefficients, and finally, MANOVA on the PC values (PC1–9) testing between the different categories among position (cervical, dorsal, caudal) and morph (wide, tall) as coded by Saitta (2015).

### Trained outline analyses

The LDA of MOMOCS outlines (i.e., elliptical Fourier coefficients) for all 40 fairly to highly complete *H. mjosi* plates trained against plate position along the body (i.e., cervical, dorsal, caudal as coded by Saitta [2015]) suggests that plate position can be readily identified, even by shape alone, independent of size (Fig. 4A). Caudal plates can be most easily recognized since they can be separated from the other plates by the first linear discriminant (LD1; 67.9% of the inter-category variance; correlates with increasing plate angle). LD2 (32.1% of the inter-category variance; correlates with increasing roundness/better-defined apex in dorsal plates and decreasing ‘blockishness’/flat top in cervical plates) then distinguishes cervical from dorsal plates. All three categories are cleanly separated in LDA without overlap. This further supports rejection of the hypothesis that variation ascribed to sexual dimorphism by Saitta (2015) is instead antero-posterior variation along the back (Mallon 2017), since a given plate’s shape is likely consistent with its position along the body and both putative tall and wide morphs can be identified in cervical, dorsal, and caudal plates. We acknowledge that as a trained analysis, uncertain assignment of body position for disarticulated plates can bias the results. Still, the ease with which the position categories can be discriminated is consistent with the idea that isolated plates can be assigned a body position with reasonable accuracy.

**Figure 4.**
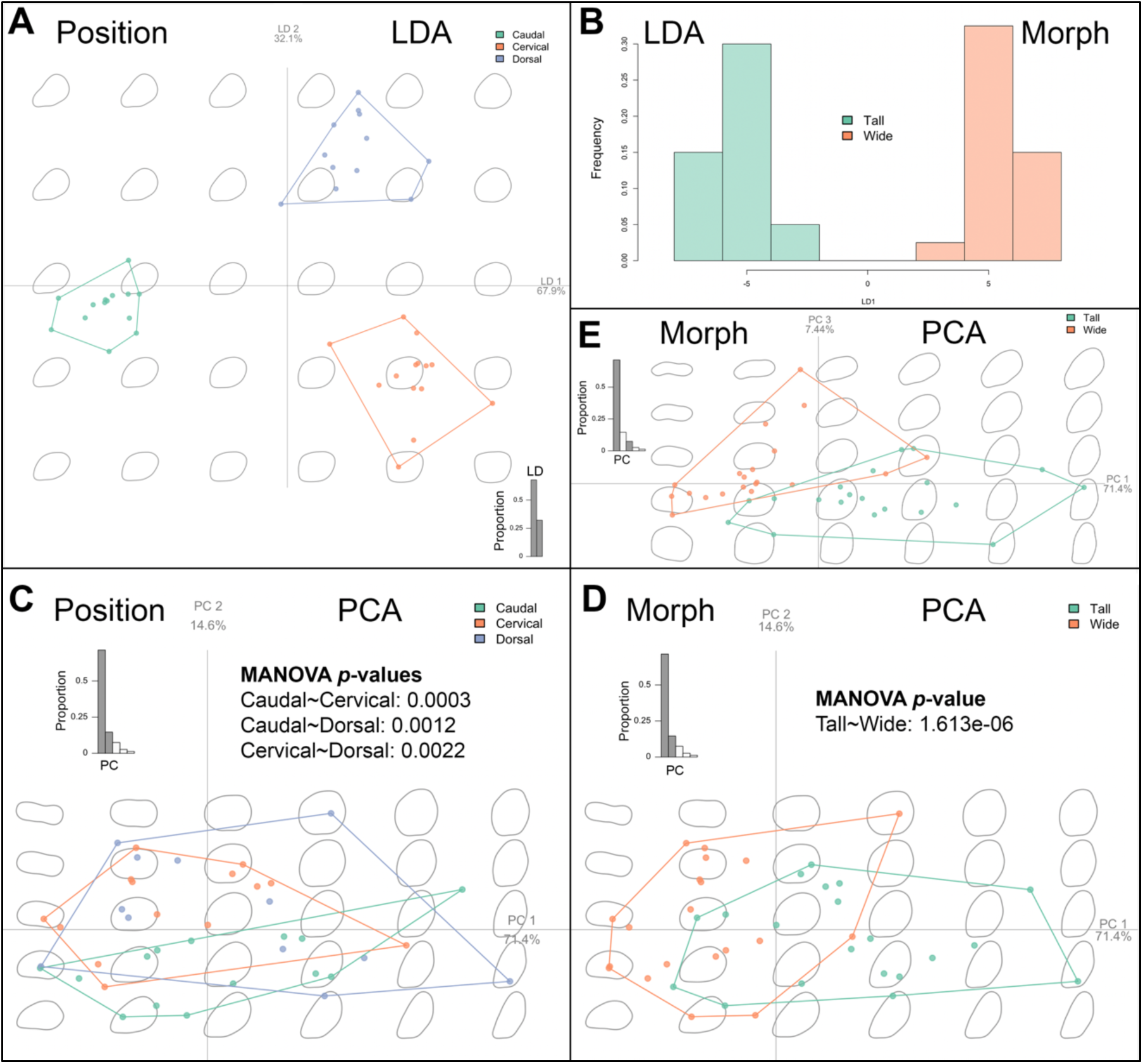
MOMOCS outline analysis of 40 fairly to highly complete *H. mjosi* plates factored by either position along the body (cervical, dorsal, caudal) or morph (tall, wide). Tall and wide morphs are coded according to the hypothesized sex assignment of Saitta (2015). A, LDA on the elliptical Fourier coefficients trained for position. LD1 (67.9%) and LD2 (32.1%) are plotted. B, LDA on the elliptical Fourier coefficients trained for morph. Only LD1 is plotted as a histogram because there are only two categories of morph. C, PCA on the elliptical Fourier coefficients grouped according to position, with minimum convex polygons shown. PC1 (71.4%) and PC2 (14.6%) are plotted. Results from MANOVA of PC values (PC1–9) are shown. D, Same PCA as in panel C, but grouped according to morph. E, Same PCA as in panel D, but with PC1 plotted against PC3 (7.44%).

The LDA of MOMOCS outlines (i.e., elliptical Fourier coefficients) for all 40 fairly to highly complete *H. mjosi* plates trained against plate morph (i.e., tall, wide as coded by Saitta [2015]) along the body suggests that shape alone, independent of size, can be used to distinguish plates of the two morphs (Fig. 4B). There is non-overlapping separation between the two morphs along LD1. Although the assignment of plate morph carries uncertainty, especially for the plates from the two multi-individual bonebeds, the results are consistent with the hypothesis that *H. mjosi* plates occur in two recognizable morphs according to shape.

### Untrained outline analyses

The PCA of MOMOCS outlines (i.e., elliptical Fourier coefficients) for all 40 fairly to highly complete *H. mjosi* plates shows that dorsal plates show the greatest variation in shape along both PC1 and PC2, compared to either cervical or caudal plates, consistent with the hypothesis that the largest plates would be the most likely to show sexual variation, especially as they might relate to a visual display function (Fig. 4C). Increasing PC1 (71.4% of plate shape variation) correlates with taller/narrower plates, consistent with hypothesized sexual dimorphism being a prominent driver of plate shape variation. Decreasing PC2 (14.6% of plate shape variation) correlates with a more pointed/better-developed apex. Overall, there is much overlap in plate shape, independent of size, across different body positions in the untrained analysis – consistent with the two plate morphologies (i.e., tall, wide) being expressed across the length of the body. MANOVA of PC values between cervical, caudal, and dorsal plates further shows that all three categories exhibit different size-independent mean shapes from each other at commonly accepted levels of statistical significance thresholds according to the *p*-value of the *F* statistic (Fig. 4C).

The PCA also shows that the two morphs (i.e. tall, wide), while showing some overlap, tend to exhibit different shapes, independent of size, with tall plates tending toward higher PC1 and lower PC2 values compared to wide plates (Fig. 4D). This is consistent with the hypothesis of sexual dimorphism. Plate morph shows less category overlap in shape than does plate position in this untrained analysis. This result is inconsistent the with hypothesis of Mallon (2017), which states that it is instead anteroposterior variation along the body that best explains proposed plate dimorphism in *H. mjosi*. Although PC3 only explains 7.44% of the variation in plate shape and is somewhat difficult to interpret (increasing PC3 perhaps correlates with an anteriorly shifted base and a posterior flange, as well as with decreasing plate angle), examining PC3 instead of PC2 further reduces overlap between the tall and wide morphs (Fig. 4E). MANOVA of PC values between tall and wide plates further shows that the two categories exhibit different size-independent mean shapes from each other at commonly accepted levels of statistical significance thresholds according to the *p*-value of the *F* statistic (Fig. 4D).

### Divergence Analyses

To account for weaknesses of the above size-independent outline analysis and examine the variation in both shape and size in *H. mjosi* plates, we can return to the PCA approach of Saitta (2015) and test for evidence of divergence in shape at larger sizes (Saitta *et al*. 2020). We have corrected the error spotted by Motani (2021) here and reperformed the PCA without excluding the anterior dorsal plate of SMA 0092 (i.e., our sample size is larger than that of the final PCA in Saitta [2015] by one plate). See Table 2 for a summary of the PCA performed here on the corrected Saitta (2015) data of 40 fairly to highly complete plates.

**Table 2.** Summary of PCA on corrected data of 40 fairly to highly complete *H. mjosi* plates (Saitta 2015), with plate angle and surface area (italics) either not transformed (as in Saitta [2015]) or transformed into units of cm to be consistent with the other four variables (as suggested by Motani [2021]). Variable loadings on each principal component are shown, with prominent loadings indicated in bold to inform upon interpretations of the PC axes. The proportion of variance for each principal component is presented as a percentage in parentheses underneath the principal component number. Transforming all of the data into cm units leads to PCA which is better described by PC1 and PC2. Also shown is the summary of PCA in which a juvenile *Stegosaurus* plate (DMNS 33359) was added to the untransformed *H. mjosi* data.

| Data transformed? | PC axis (proportion of variance) | *Angle between base and apex* | Length of base | Perimeter | Width' | Distance from center of the base to apex | *Surface area* | PC axis interpretation |
| --- | --- | --- | --- | --- | --- | --- | --- | --- |
| No<br>(Saitta 2015)<br>*H. mjosii*<br>*n* = 40 plates | **PC1 (78.32%)** | 0.02 | **-0.45** | **-0.46** | **-0.44** | **-0.43** | **-0.46** | Size |
|  | PC2 (16.95%) | **0.99** | 0.02 | -0.05 | 0.08 | -0.09 | 0.07 | Body position: posterior to anterior |
|  | **PC3 (3.17%)** | 0.13 | -0.29 | 0.13 | **-0.52** | **0.78** | -0.08 | Shape: wide to tall |
| No<br>(Saitta 2015)<br>*H. mjosii* + Juvenile *Stegosaurus*<br>*n* = 41 plates | **PC1 (78.44%)** | 0.04 | **-0.45** | **-0.46** | **-0.44** | **-0.43** | **-0.45** | Size |
|  | PC2 (16.89%) | **0.99** | 0.03 | -0.04 | 0.08 | -0.08 | 0.08 | Body position: posterior to anterior |
|  | **PC3 (3.13%)** | 0.13 | -0.29 | 0.13 | **-0.51** | **0.78** | -0.07 | Shape: wide to tall |
| Data transformed? | PC axis (proportion of variance) | *Height of plate above back* | Length of base | Perimeter | Width' | Distance from center of the base to apex | *Square root of surface area* | PC axis interpretation |
| Yes<br>(Motani 2021)<br>*H. mjosii*<br>*n* = 40 plates | **PC1 (93.95%)** | **-0.41** | **-0.4** | **-0.42** | **-0.4** | **-0.4** | **-0.42** | Size |
|  | **PC2 (4.64%)** | **0.48** | **-0.38** | -0.02 | **-0.58** | **0.54** | -0.05 | Shape: wide to tall |
|  | PC3 (1.11%) | 0.04 | **0.83** | -0.25 | **-0.42** | 0.08 | -0.26 | Uncertain: widening to narrowing apically from base? |

We also ran another PCA using the transformations of area and angle values, as suggested by Motani (2021), to get all measures into the same linear units (i.e., cm). We took the square root of the area. We also took the sin values of the angles, multiplied by the distance from the center of the base to the apex to get roughly the extent to which the plate rose above the back of the animal. This allows us to describe plate size versus shape using PC1 and PC2, rather than PC1 and PC3. This variable transformation further results in the principal components of size and shape describing a combined 98.59% of the variation in plates, compared to 81.49% without variable transformation.

### Considering ontogenetic & intra-individual variation

Initially, Saitta (2015) simply observed the separation of the data along two PC axes that represented size and shape. Here, we investigate this separation further.

Smaller plates of known stegosaur specimens are generally the anterior cervical and posterior caudal plates rather than plates from very young and small individuals. Almost all specimens examined histologically by Saitta (2015) were likely at least young adults, such as the sexually mature but still growing SMA 0092 and VFSMA 001 (Redelstorff & Sander 2009; Hayashi *et al*. 2012), with only the WDC plates of uncertain ontogenetic status. A method to investigate sexual dimorphism in fossils as described by Saitta *et al*. (2020) might instead assume that size data is an appropriate proxy for age (i.e., that the separation observed is controlled against ontogeny). However, a sample of plates known to come almost entirely from young adults and old adults will contain the most informative region of plate growth dynamics for potential sexual dimorphism, because dimorphism is expected to correlate with sexual maturity and be most prevalent in adults. Furthermore, we predict that dimorphism would be most prevalent in the largest plates, if plates were under sexual selection for a visual display function.

To further judge whether ontogeny is sufficiently controlled for in our PCA, we can examine the only known plate from a very small juvenile of *Stegosaurus*, a complete dorsal plate (DMNS 33359). No juvenile plates are known from *H. mjosi*. When DMNS 33359 measurements are included with the *H. mjosi* data of Saitta (2015) and the PCA is rerun, the same overall pattern persists. The juvenile dorsal plate falls within the region of the morphospace of the smallest tall-morph plates and has a positive PC3 value (Fig. 5). The tall morph was hypothesized by Saitta (2015) to be female, and a juvenile stegosaur dorsal plate having similar shape to the small plates on larger females would not be implausible (i.e., juvenile males often resemble females). This limited result suggests the PCA of *H. mjosi* plates can also account for ontogenetic variation in addition to intra-individual variation anteroposteriorly along the back.

**Figure 5.**
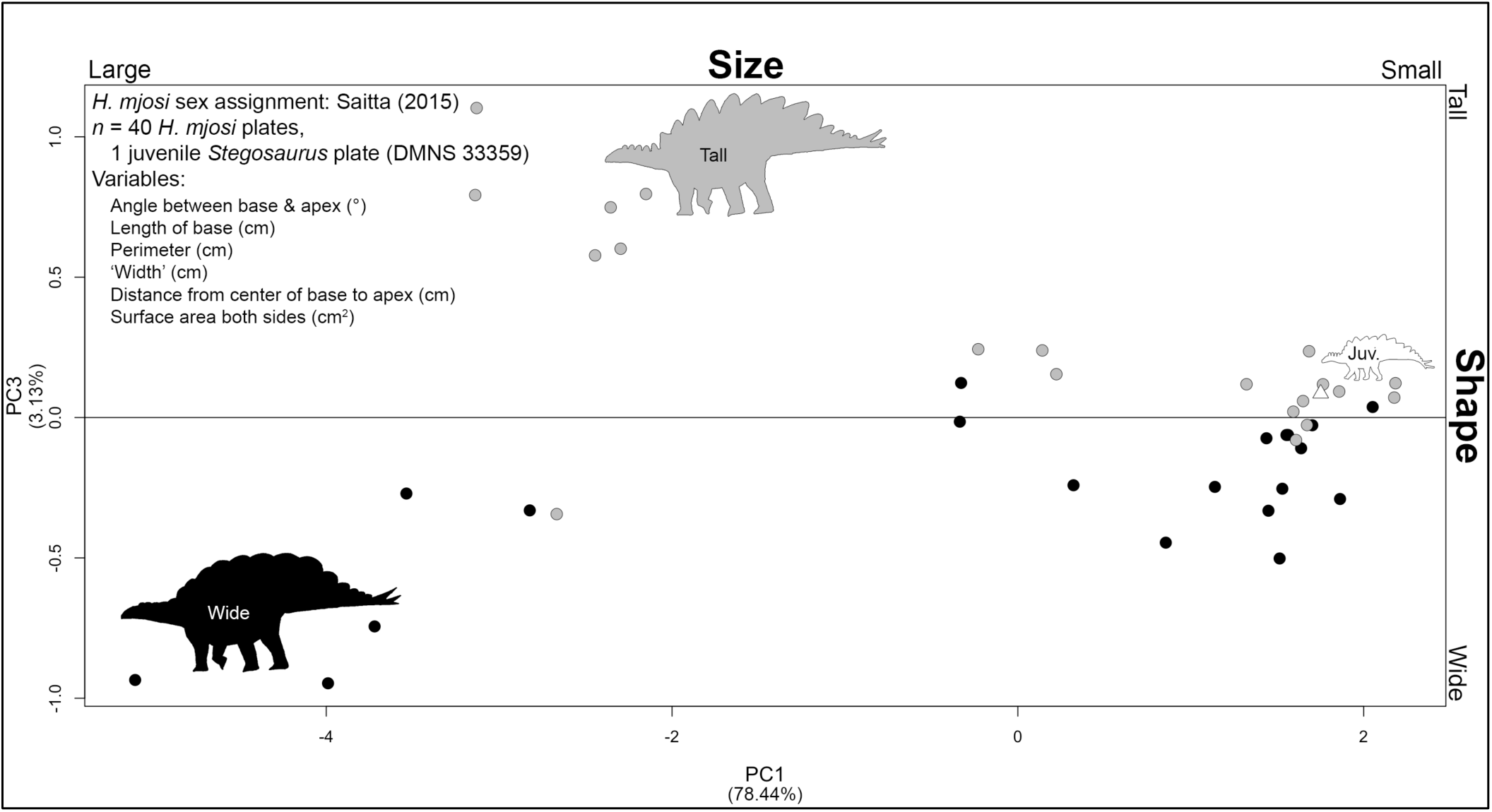
PCA of corrected data from 40 fairly to highly complete *H. mjosi* plates from Saitta (2015) without variable transformation (six morphometric variables) and with a juvenile *Stegosaurus* plate (DMNS 33359, white triangle) added to the dataset (*n* = 41 plates) to compare to the otherwise sexually mature sample. PC1 describes size and PC3 describes shape. Grey and black points are those identified by Saitta (2015) as tall and wide morphs, respectively.

### The challenge of non-independence

Another way to view this challenge when studying structures such as stegosaur plates is that data points are only sometimes independent, because multiple plates can be derived from a single individual. Examining plates only from specific plate positions along the back of the animal might be counterproductive if their function depended upon a combined effect. For example, stegosaur visual stimuli might have utilized a ‘chorus effect’ of all plates being displayed as singular unit – the plate series as a whole profile.

The problem of non-independence is most apparent during the sex estimation step. Plates from the same individual cannot be assigned to different sexes. Since 25 out of 40 *H. mjosi* plates examined by Saitta (2015) come from isolated individuals, where we can ensure that mixed-sex assignments are avoided, the effect of this complication is reduced. For this reason, we use the categorical sex assignments from Saitta (2015) for our divergence analysis. Furthermore, residual plots after linear regression applied to the entire PCA morphospace (Fig. 6A–B) show that divergence/bifurcation of the data can be readily observed before any attempt to categorize the two sexes/morphs. Indeed, very few plates categorically assigned to a particular morph lie on the opposite region of the morphospace (i.e., positive versus negative PC values for shape). Most exceptions are small plates expected to fall relatively close to a PC value of zero for shape.

**Figure 6.**
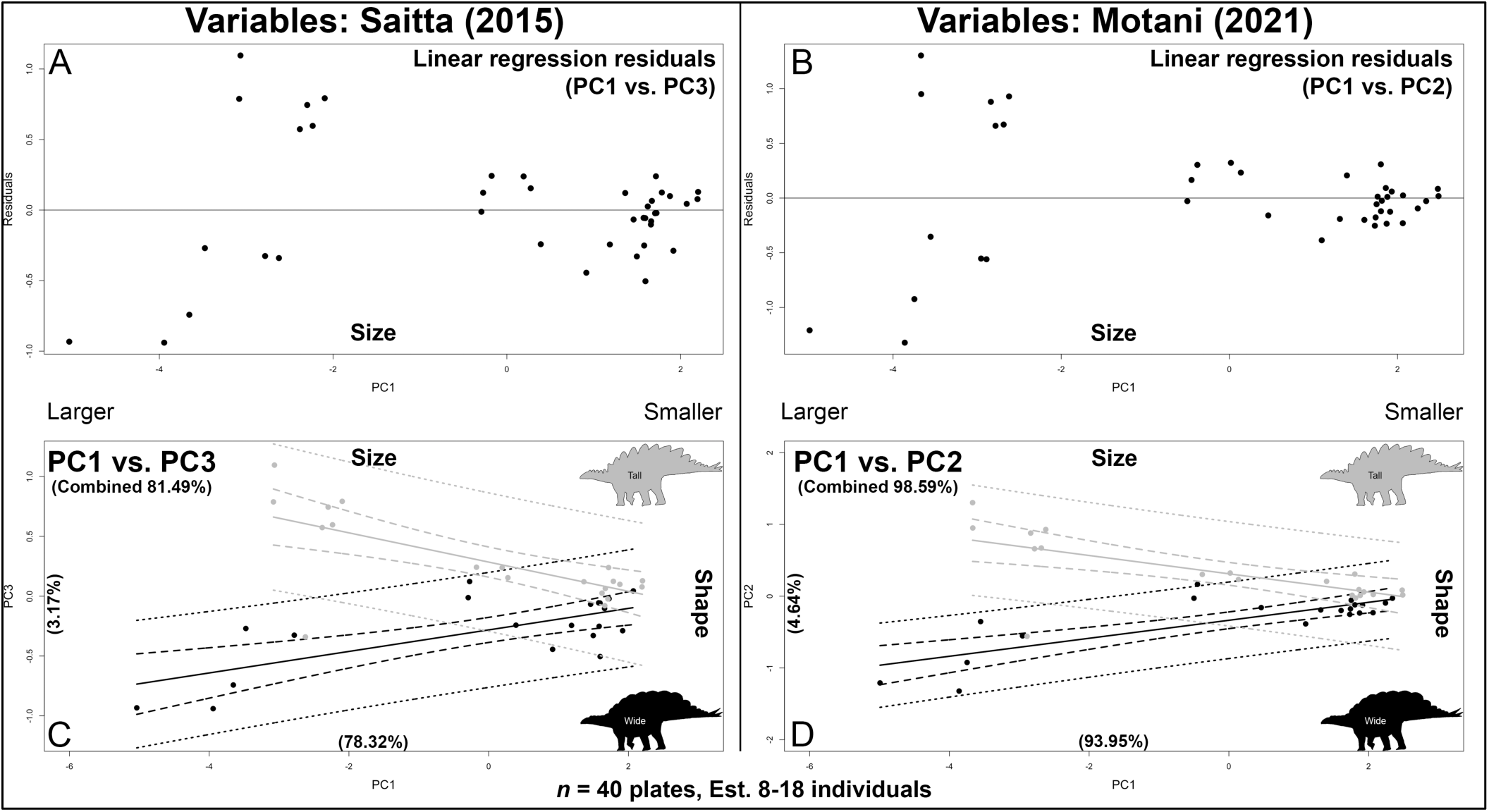
Divergence analysis of 40 fairly to highly complete *H. mjosi* plates based on PCA of six morphometric variables. A–B, Residual plots after linear regression was performed on the total dataset, plotted against the size descriptor (PC1). C–D, PCA morphospace shows size (PC1) vs. shape (either PC3 or PC2). Assigned sex in both C–D follows the categorization of tall (grey) and wide (black) morphs in Saitta (2015). Dashed lines are 95% confidence intervals, while dotted lines are 95% prediction intervals. Results from both (A,C) non-transformed (Saitta 2015) and (B,D) transformed variables (Motani 2021) are shown. Transformation of the variables into the same linear units improves the total percentage of variation in plates explained by the two axes of the morphospace. Both approaches result in the separation of both confidence and prediction intervals at larger plate sizes.

### Divergence & confidence in PCA

When examining the divergence of *H. mjosi* plate shape (i.e., tall versus wide morphs) using PCA (Fig. 6C–D), both 95% confidence and prediction intervals separate at larger plate sizes, even with the inclusion of the unusually wide-based anterior dorsal plate of SMA 0092 that was erroneously dropped in Saitta (2015). Therefore, it can be estimated with high confidence that larger plates in *H. mjosi* do indeed diverge in shape across two hypothesized morphologies suggestive of sexual dimorphism.

As more *H. mjosi* fossils are found, additional data might reinforce this trend (e.g., increased difference in regression slopes between the two morphs, increased range in plate sizes, increased range in the shape of the largest plates, or narrowed confidence and prediction intervals). Alternatively, new data might weaken this trend – although note that sexually variable traits often show considerable overlap, such as human height dimorphism (Schilling *et al*. 2002).

The currently observed differences in *H. mjosi* plate size and shape could be further amplified when accounting for unpreserved keratin sheathing (e.g., as preserved on a plate in SMA 0118 [Christiansen & Tschopp 2010]). In this case, variation in the size and shape plates could be greater in the outermost keratin than in the underlying bone that more readily fossilizes. Note that the *H. mjosi* tail spike from the JRDI 5ES Quarry identified in the undergraduate thesis of Saitta (Saitta 2014) as preserving a phosphatically fossilized keratin sheath was most likely misidentified, instead representing unusually crushed and fragmented bone of the spike’s tip.

### Specimens with Plates Possibly Similar to Those of *Hesperosaurus*

While *Stegosaurus* was first named in 1877 (Marsh 1877), *H. mjosi* was described in 2001 (Carpenter *et al*. 2011). The *H. mjosi* holotype, HMNS 14, was found near the town of Buffalo in northern Wyoming (Carpenter *et al*. 2001). Could additional, unidentified *H. mjosi* specimens already exist in museum collections, mislabeled as *Stegosaurus*? With variation in *H. mjosi* plates now better characterized, plates from additional specimens with similar morphologies and uncertain or undescribed taxonomy can be reexamined. We do not make any concrete taxonomic diagnoses of these specimens here through thorough investigation of skeletal autapomorphies. Rather, we simply wish to highlight specimens and their plate morphology and how they might compare to tall and wide *H. mjosi* morphs.

Even after 2001, additional *H. mjosi* specimens likely exist whose current curatorial status is uncertain. Heritage Auctions claimed in 2011 (Richter 2011) that it sold what was presumably a wide-plated specimen of *H. mjosi* (i.e., “Fantasia”) from the Dana Quarry near Ten Sleep in northern Wyoming (Albersdörfer & Galiano 2009; Saleiro & Mateus 2017) to an undisclosed museum outside of the USA. However, it is unclear what or how much of that specimen is original fossil material. The Knuthenborg Museum of Evolution recently announced a new, highly complete stegosaur specimen (i.e., “Stephanie”) from Wyoming which resembles *H. mjosi* and at least one of its plates may have a tall morphology from available photos online, but a more complete description of the specimen is not yet available. Similarly, SMA is currently preparing a stegosaur specimen (i.e., “Cheyenne”) from the Meilyn Quarry near Como Bluff, Wyoming (H.J. Kirby Siber, pers. comm. 2025). Although its precise taxonomic identity is still uncertain, one possibility is that it is a wide-plated *H. mjosi* based on photos shared by the SMA.

UCRC PV6 (Fig. 7B) is a highly complete plate found with associated stegosaur skeletal material still undergoing preparation. It resembles the anterior dorsal plate of the tall morph *H. mjosi*, as seen in SMA 0092 – with a wide base, but a well-defined triangular apex. UCRC PV6 was found in the Big Al Quarry near Shell in northern Wyoming by a University of Chicago team under P. C. Sereno. Again, the area near Shell, Wyoming is notable for producing several *H. mjosi* specimens (SMA 0018, SMA 0092, VFSMA 001).

**Figure 7.**
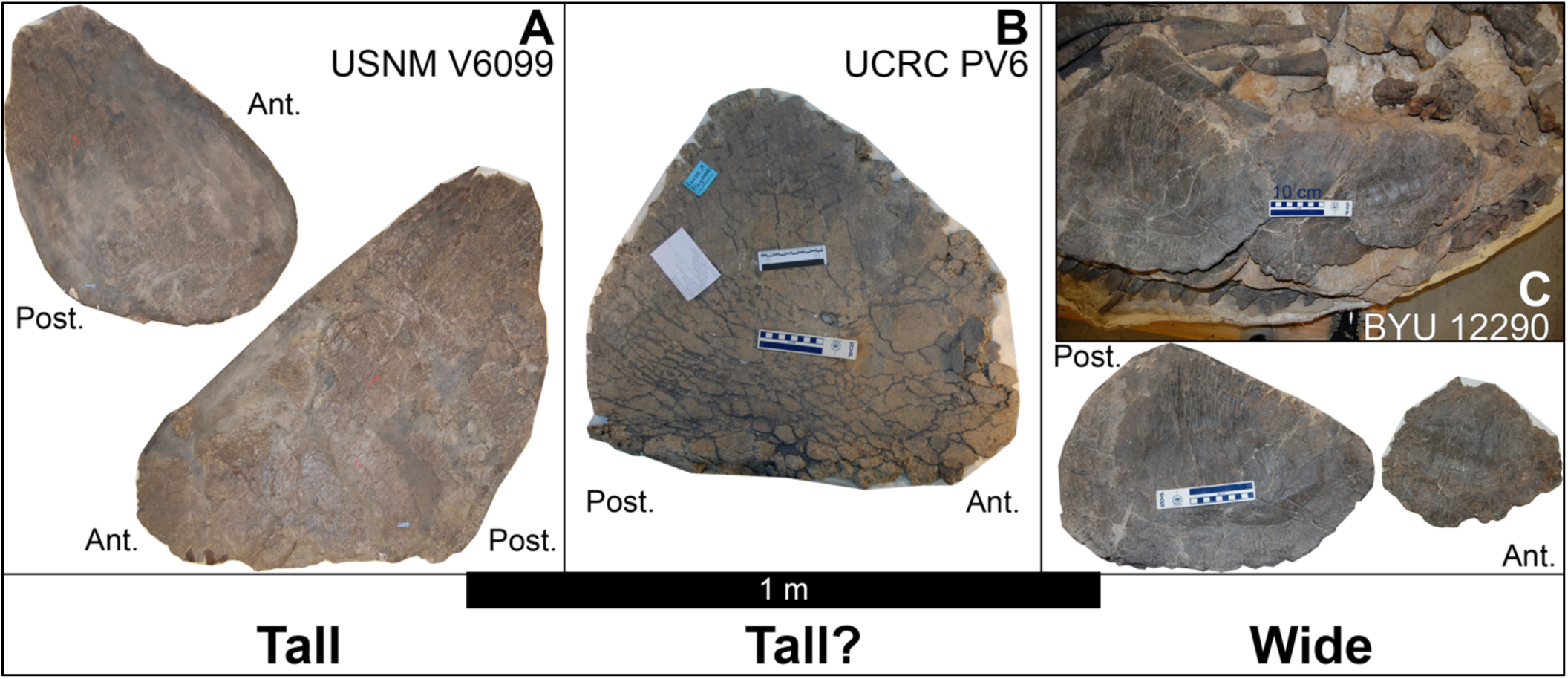
Other North American stegosaur specimens with plates similar to those of *H. mjosi*. A, USNM V6099, consistent with a tall morphology. B, UCRC PV6, possibly consistent with a tall morphology. C, BYU 12290, consistent with a wide morphology. The black scale bar applies to all of the isolated images of plates, while the photograph of the articulated skeleton of BYU 12290 (top of panel C) includes a scale bar of 10 cm length within it (left side of scale bar).

Further back in history, the specimen number USNM V6099 (Gilmore 1914) applies to two plates of fair, but not high, completeness (Fig. 7A). They are similar to the tall morphology in *H. mjosi*. This material comes from the very productive Como Bluff Quarry 13 in southeast Wyoming. The WDC DMQ *H. mjosi* bonebed (i.e., Meilyn Quarry) also comes from the Como Bluff area. USNM V6099 was initially described as *Stegosaurus ungulatus*.

Finally, BYU 12290 (Fig. 7C) preserves an articulated torso with three fairly to highly complete oval plates in articulation with each other in two staggered rows but flipped down over the ribcage (i.e., the ventral bases are now directed dorsally). These plates are similar to those of wide-morph *H. mjosi*. This relatively small-bodied specimen, further consistent with *H. mjosi* in size, was found in the Jensen/Jensen Quarry near Jensen in northeastern Utah, further south than what might be expected based on the localities of other *H. mjosi* specimens.

## OSTEODERM VARIATION, FUNCTION, & EVOLUTION ACROSS NORTH AMERICAN STEGOSAURS

### Combat Before Display?

While much of this discussion has focused on plate variation, if the hypothesis for sexual dimorphism in *H. mjosi* plates is indeed supported, it raises the possibility of sexual selection acting upon osteoderms. Therefore, it is worth questioning if tail spikes in North American stegosaurs functioned solely for defense against predation. Farke (2014) noticed that stegosaur spikes are consistent with either inter-or intra-specific combat based on biomechanical modelling (Carpenter *et al*. 2005; Mallison 2011; Arbour 2009) and pathology. Dynamic modeling *in silico* has suggested that some stegosaurs could swing their tails at high speeds to produce powerful strikes (Mallison 2011; Lategano *et al*. 2024).

The main argument against tail spikes being used in intra-specific or intra-sex combat might be the avoidance of highly injurious ‘total war’. Under the hypothesis of ‘total war’ avoidance (Maynard Smith & Price 1973), long-lived species have multiple opportunities to mate across their lifetime, so they opt for less injurious ‘limited war’ strategies in sexual conflict. Such a strategy would especially apply to younger individuals compared to older ones approaching senescence. Under ‘limited war’ strategies, the presumed injury that could have been produced through a forceful impact by a thagomizer (Carpenter *et al*. 2005; Mallison 2011; Arbour 2009) might have precluded males, for example, from using them in sexual competition. It is unclear if stegosaur tail spikes could have pierced bone, as studies attempting to estimate such forces have disagreed (Carpenter *et al*. 2005; Arbour 2009; Mallison 2011).

In many species that engage in intraspecific combat, the weapons used are blunt. While effective in combat, they are less likely to result in lethal injuries. As such, one possibility is that the keratinous sheath around stegosaur spikes was blunter than the underlying bone, although there is currently no direct evidence for this preserved one way or the other. The blunt horns of bighorn sheep (Ackermans *et al*. 2022), horn bases of muskoxen (Ackermans *et al*. 2022) and cape buffalo (Prins 1989), and ossicones of giraffes (Simmons & Scheepers 1996) are all structures that are effective in combat but non-lethal or at least less lethal. The large head of male sperm whales has been suggested to function as a battering ram to attack other males (Panagiotopoulou *et al*. 2016). A few species of birds have blunt knobs on the wing that are used to strike other birds, including steamer ducks (*Tachyeres*) (Nuechterlein & Storer 1985), sheathbills (*Chionis*) (Chu 1995), and crowned pigeon (*Goura*) (Rico-Guevara & Hurme 2019). Similarly blunt hornlets or thickened skulls are also seen in some non-avian dinosaurs, including Pachycephalosauridae (Maryańska *et al*. 2004) and ceratopsids such as *Pachyrhinosaurus* (Sternberg 1950). These dinosaurian structures have been hypothesized to have functioned in intraspecific combat (e.g., Snively & Theodor 2011).

### Violent intraspecific combat

However, other species evolve weapons that appear to be more lethal. Among mammals, these include the large canines of elephant seals (Briggs & Morejohn 1975) and certain deer such as chevrotains (Terai *et al*. 1998), the bladelike teeth of certain beaked whales (Besharse 1971), and the tusks of wild boars (Barrette 1986), which are used in intraspecific combat. Various birds have dagger-like spurs on the wings, including plovers (*Vanellus*) (Meissner *et al*. 2021), screamers (Anhimidae), and spur-winged geese (*Plectropus gambensis*) (Rand 1954). Sharp spurs are also developed on the feet of male phasianids, including jungle fowl (*Gallus*) (Desta 2019), peacocks (*Pavo*), and turkeys (*Meleagris*) (Davidson 1985).

Indeed, intra-sex combat over mates in extant animals can often be extremely arduous, dangerous, and sometimes lethal for the individuals involved. This can be true even in relatively long-lived species and for individuals of high rank compared to their intra-sexual competition. In greater kudus, their horns sometimes get locked together, leading to the death of both sparring combatants (Owen-Smith 1993). Elephant seals use their chests and teeth to wound rivals in bloody fights (Leboeuf 1972; Haley *et al*. 1994). Throughout the breeding season, high-ranking male elephant seals with high reproductive success expend more energy and experience greater loss of mass than do lower-ranking males, dropping over 40% of their initial body weight (Deutsch *et al*. 1990). Beyond the risk of cranial bone fracture (Peterson *et al*. 2013), headbutting animals, such as bighorn sheep and muskoxen, are known to experience traumatic brain injuries (Ackermans *et al*. 2022). Even violence in humans, especially men, for social status can be dangerous. Severe health risks and injuries can befall professional athletes, such as in American football or combat sports.

Pathologies in thyreophoran ankylosaur osteoderms consistent with the use of tail clubs in intraspecific combat (Arbour *et al*. 2022) and analogy to extant lizards using tail whips in intraspecific combat (Schall *et al*. 1989) require us to take seriously the possibility that stegosaur tail spikes might have been under sexual selection as weapons in intra-sex combat, in addition to their presumed utility as defensive weapons. Furthermore, a long and flexible tail (Mallison 2010, 2011), small forelimb to hindlimb ratio, and center of mass located near the hips (Henderson 1999; Maidment *et al*. 2012; Mallison 2014), might have stabilized the hips during tail whips and allowed for easier rotation when pushing off with the forelimbs to direct the tail towards the target. These biomechanics would also be consistent with the use of tail spikes as intrasexual combat weapons.

Osteoderms appear to be involved in some cases of modern intrasexual aggression. In two populations of extant Cape cliff lizards, osteoderms likely undergo selection for intrasexual aggression based on several lines of evidence (Broeckhoven *et al*. 2017b). 1) The osteoderms develop at or after the onset of sexual maturity from the lateral to medial sides of the body, consistent with secondary sexual characteristics and with osteoderms protecting against bites to the lateral flank. 2) Males have greater osteoderm volume than females. Finally, 3) bite force (a proxy for intrasexual aggression) correlated with osteoderm expression in the two lizard populations (Broeckhoven *et al*. 2017b).

### Prehistoric lekking?

Males of many extant taxa may first compete against each other before displaying to females, particularly over territorial positions in leks. For example, male great snipes and black grouse will fight each other in leks, sometimes ripping out the feathers of their rival (Alatalo *et al*. 1991; Höglundi *et al*. 1992). Some ungulates will form leks, such as the topi, where males will engage in intense agonistic behavior and repeatedly spar with each other using their horns, which will often break, before displaying to females using a high-step gait (Gosling *et al*. 1987; Bro-Jørgensen & Durant 2003). Fighting between males in fallow deer and lechwe antelope using antlers and horns, respectively, is more frequent when they are lekking (Apollonio *et al*. 1992; Nefdt & Thirgood 1997). In birds of paradise (Paradisaeidae), lekking species show greater sexual dichromatism, and lekking species where males directly compete against each other in communal display courts have more carotenoid-pigmented ornaments and fewer melanin-pigmented ornaments (Miles & Fuxjager 2018).

Stegosaur spikes and plates could have allowed for this behavior, and lekking would be consistent with these differentiated osteoderms in *Stegosaurus* and *H. mjosi*. Displays using the plates could have reduced the risk of ‘total war’, allowing males to size each other up before deciding to engage in violent combat. Sizing up can occur in extant animals, such as chest-beating behavioral displays of gorillas used to avoid dangerous fights between unevenly sized males (Wright *et al*. 2021). Male elephant seals recognize the acoustic displays of their rivals and respond according to how they performed against those rivals in previous contests, thereby avoiding highly costly conflict (Casey *et al*. 2015).

In stegosaurs, male combat could have progressed as follows: display of osteoderms to rival males to judge size before combat, then combat between evenly sized males using their thagomizers, then display of osteoderms to females by high-status males after successful competition against other males, and finally, choosing of male mates by the females. With the ability to reconstruct paleo-color from some exceptionally preserved fossils and knowing that pigments such as carotenoids can fossilize (Roy *et al*. 2020), it will be interesting to see if future fossil finds and methodological advancements might allow for color reconstruction of stegosaur plates. Would those results be consistent with brightly colored, lekking species in which males compete against each other? Possessing both sexual ornaments and armaments in stegosaurs was likely not unique within Ornithischia. Likewise, ceratopsian frills and horns could fill combat and display roles, respectively (Farlow & Dodson 1975). Note that substrate scratching track marks used as evidence of lekking courtship displays in large theropods (Lockley *et al*. 2016) could also represent other forms of behavior (Saitta *et al*. 2020) and might be less likely to represent lekking behavior if large carnivores are predisposed to large territories and solitary behavior/small social group sizes due to elevated food requirements.

### Stegosaur tail spike & caudal vertebrae pathologies

Structures used for intraspecific combat frequently sustain injuries. For example, injuries are common on the skull of *Triceratops*, likely as a result of intraspecific combat (Farke *et al*. 2009). Similarly, a high frequency of pathology in the domes of pachycephalosaurids is thought to result from intraspecific combat (Peterson *et al*. 2013).

Although possibly consistent with defense against predation, pathologies in North American stegosaurs might have resulted from forceful tail swings during intraspecific combat. Caudal vertebrae pathologies could represent bones broken during intraspecific combat that subsequently healed, became infected, and/or fused together (Fig. 8A–H). Similar fusions in sauropod vertebrae (Cruzado-Caballero *et al*. 2023) have been described as diffuse idiopathic skeletal hyperostosis and linked to both sexual dimorphism (Rothschild & Berman 1991) and tail whipping behavior (Myrhvold & Currie 1997).

**Figure 8.**
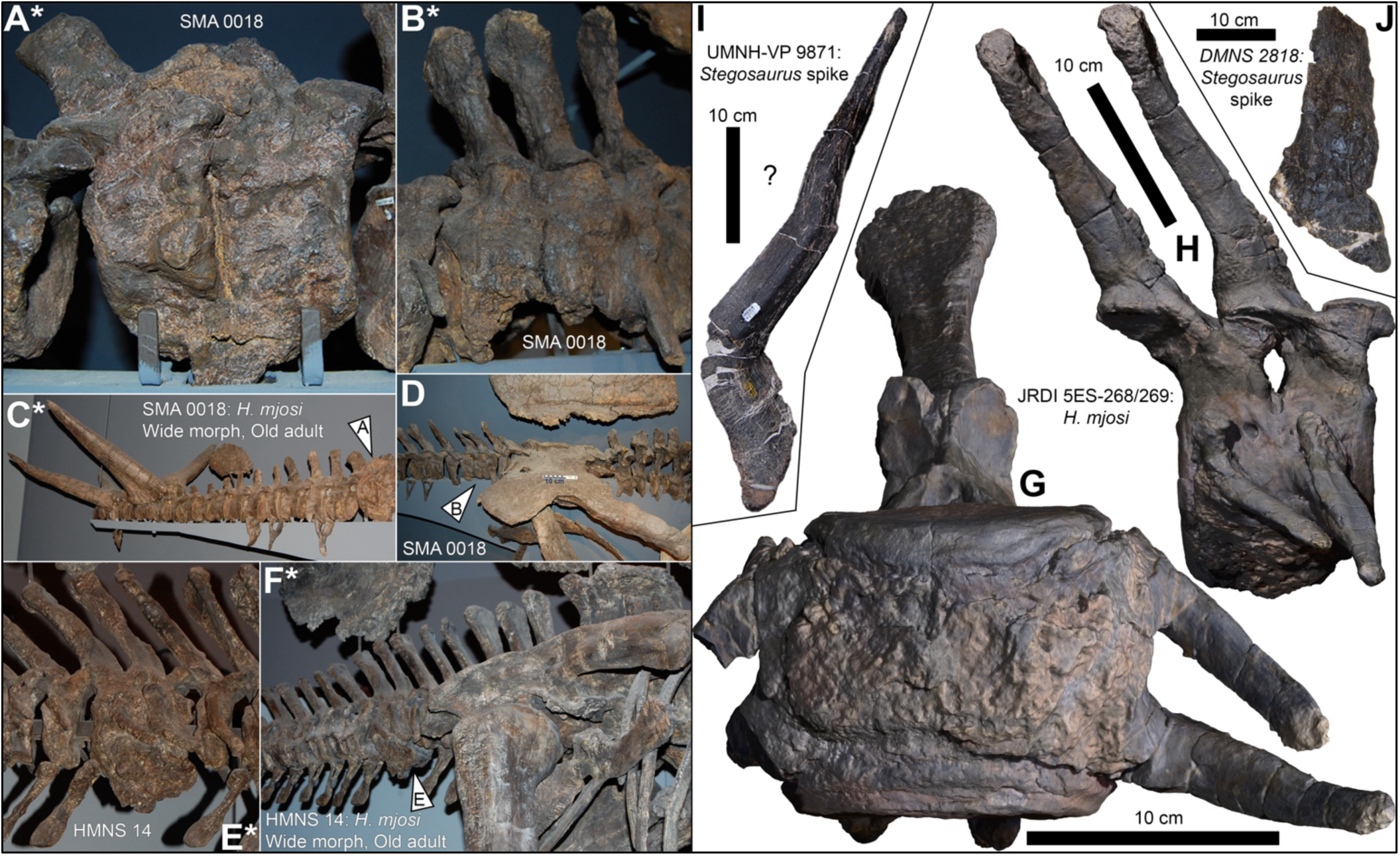
Possible pathologies on North American stegosaur caudal vertebrae and tail spikes, some of which may be consistent with either interspecific defense or intra-sex combat (e.g., male combat). Pathologically fused posterior, A, and anterior, B, caudal vertebrae from wide-morph, old adult *H. mjosi* SMA 0018, C–D. Pathologically fused anterior caudal vertebrae, E, from wide-morph, old adult *H. mjosi* HMNS 14, F. The asterisks indicate that no scale bar is included due to the photographs being taken from a distance and at an oblique angle to the mount. The physical scale bar on the ilium in D shows a 10 cm scale on its left side. Arrows in C–D, F indicate location of pathologies on the mounted skeletons. G–H, Photogrammetry model of pathologically fused caudal vertebrae JRDI 5ES-268/269 from the central Montana (near Grass Range) multi-individual *H. mjosi* bonebed. I, an extremely bent *Stegosaurus* tail spike UMNH-VP 9871 showing diagenetic plastic deformation, trampling bioturbation, and/or *in vivo* deformity. Proximal base to the bottom. J, *Stegosaurus* tail spike from DMNS 2818 with obvious pathology consistent with injury. Proximal base to the bottom.

Interestingly, these caudal vertebrae pathologies are present in wide-morph, hypothetical male *H. mjosi*. These specimens, SMA 0018 and HMNS 14, are known through histology to be fully grown, old adults that would have been sexually mature for multiple years prior to death. They were, therefore, capable of accumulating combat-related injuries over time. Additionally, they might have opted for riskier confrontations under ‘total war’ strategies as they approached senescence (Maynard Smith & Price 1973), although this would assume that stegosaurs did experience some degree of reproductive senescence (de Magalhães 2023). One of these specimens, SMA 0018, is also reported to have pathologies on the neural spines of the sacral vertebrae (Siber & Möckli 2009). Similar pathologies in tails and pelvises have been noted in ankylosaurs (Arbour & Currie 2013).

Pathologies are well known in *Stegosaurus* spikes. Posttraumatic osteomyelitis has been proposed in the spikes of DMNS 2818 (Fig. 8J) and USNM 6646, for example. In a survey of 51 spikes, an estimated ∼10% might show similar pathologies (McWhinney *et al*. 2001). Although this type of tail spike pathology is often implicated with predator-prey interaction (Carpenter *et al*. 2005), it could also derive from intra-sex combat.

Another *Stegosaurus* spike, UMNH-VP 9871, is extremely bent (Fig. 8I). Most of this deformation is along a single plane, suggestive of diagenetic plastic deformation during burial or of trampling bioturbation hypothesized to have occurred at the Cleveland-Lloyd Dinosaur Quarry (Gates 2005). However, its back-and-forth ‘zigzag’ is extreme and potentially unusual. Further work, especially histological examination, could determine if any of this deformation is pathological. Some of the deformation in this spike could be similar to *in vivo* or congenital horn deformities of extant bovids, including highly bent horns that abruptly change direction (Lyon 1908; Ackermann *et al*. 2010). While not necessarily related to injury, if so, it would be interesting if stegosaur spikes experienced similar developmental abnormalities as bovid horns. This is because horns are commonly used structures in intraspecific combat. In extant taxa, horns are under sexual selection across diverse clades and disparate horn morphologies (Saitta *et al*. 2020).

### Spike dimorphism is currently uncertain

One powerful line of evidence that stegosaurs used their tail spikes for intra-sex combat, and not only for defense against predators, would be evidence of sexual dimorphism in the spikes. Unfortunately, our available *H. mjosi* sample currently consists of only 13 spikes. However, our sample of 36 *Stegosaurus* tail spikes can more readily be examined for variation within (Fig. 9). Among these, most *Stegosaurus* spikes correspond to two size modes: adult posterior (shorter) and adult anterior (longer) spikes. Note that the giant spikes once described as *Stegosaurus longispinus* are now considered to be from an entirely different genus, *Miragaia longispinus* (Costa & Mateus 2019). Therefore, these spikes are possibly incomparable to the *Stegosaurus* terminal thagomizer if *M. longispinus* spikes were positioned more anteriorly (e.g., near the shoulders).

**Figure 9.**
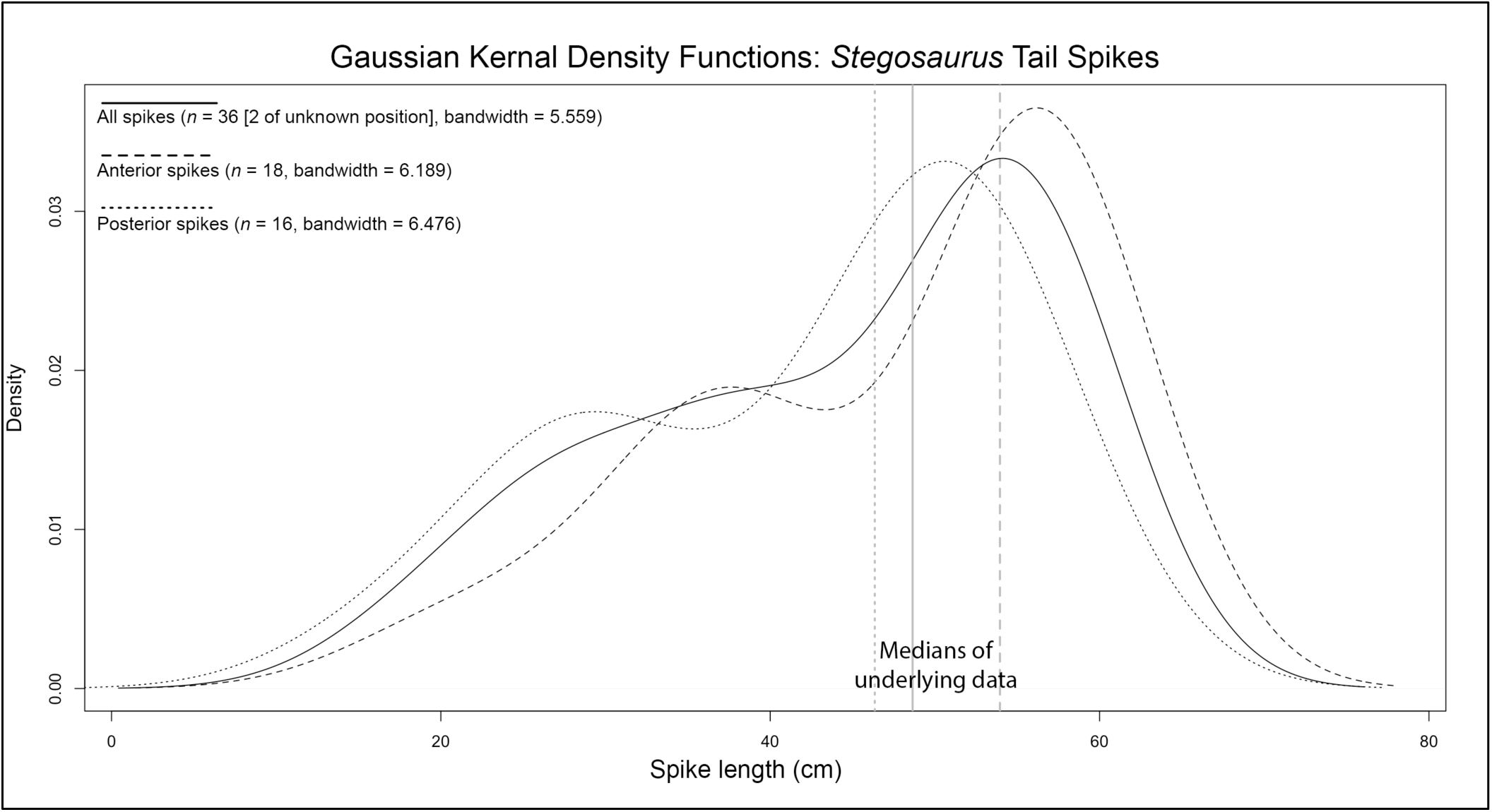
Tail spike proximal-distal length distribution in *Stegosaurus* (i.e., excluding *H. mjosi*). Distributions approximated as Gaussian kernel density functions using the default density() function in R, with sample sizes and bandwidths reported. Three functions are shown: all spikes, anterior spikes only, and posterior spikes only. Vertical lines indicate the median length of the underlying spike datasets (i.e., prior to applying the density function).

Although there is potential bimodality present in both posterior and anterior spike density functions, this could simply signify an oversampling of younger individuals at that corresponding thagomizer size rather than sexual variation. Since thagomizers consist of two pairs of spikes, this bimodality could also be influenced to oversampling the left and right sides from the same individuals at that corresponding thagomizer size. Future work could build upon this sample while controlling for 1) ontogeny through histological analyses, 2) overweighting caused by multiple spikes from the same individual, and 3) any probable species diversity within the genus *Stegosaurus*. For now, the argument for sexual dimorphism in stegosaur spikes is far more lacking than in the plates of *H. mjosi*.

### Evolution & Function of Stegosaur Plates, Spikes, & Ossicles

Osteoderms have evolved independently many times across vertebrates, apart from birds (Maden *et al*. 2023) and have been proposed to perform diverse functions involving protection, sensory biology, physiology, and ecology/locomotion (Ebel *et al*. 2024). With the above considerations, it is possible to propose a hypothetical model for stegosaur osteoderm evolution and function that best explains their morphological variation (Fig. 10), following the example of others before (e.g., Farlow *et al*. 2010). This model takes multiple evolutionary drivers into consideration. Although the model is clearly hypothetical, it represents a synthesis of what we can reasonably predict about stegosaur (and more broadly, thyreophoran) evolution based on both their fossil record and an understanding of extant organisms. Future data will either support or alter this proposed evolutionary dynamic framework. It should also be noted that, although we use male and female descriptors based on widespread trends among amniotes, we are not excluding the possibility of sex-role reversal (Eens & Pinxten 2000).

**Figure 10.**
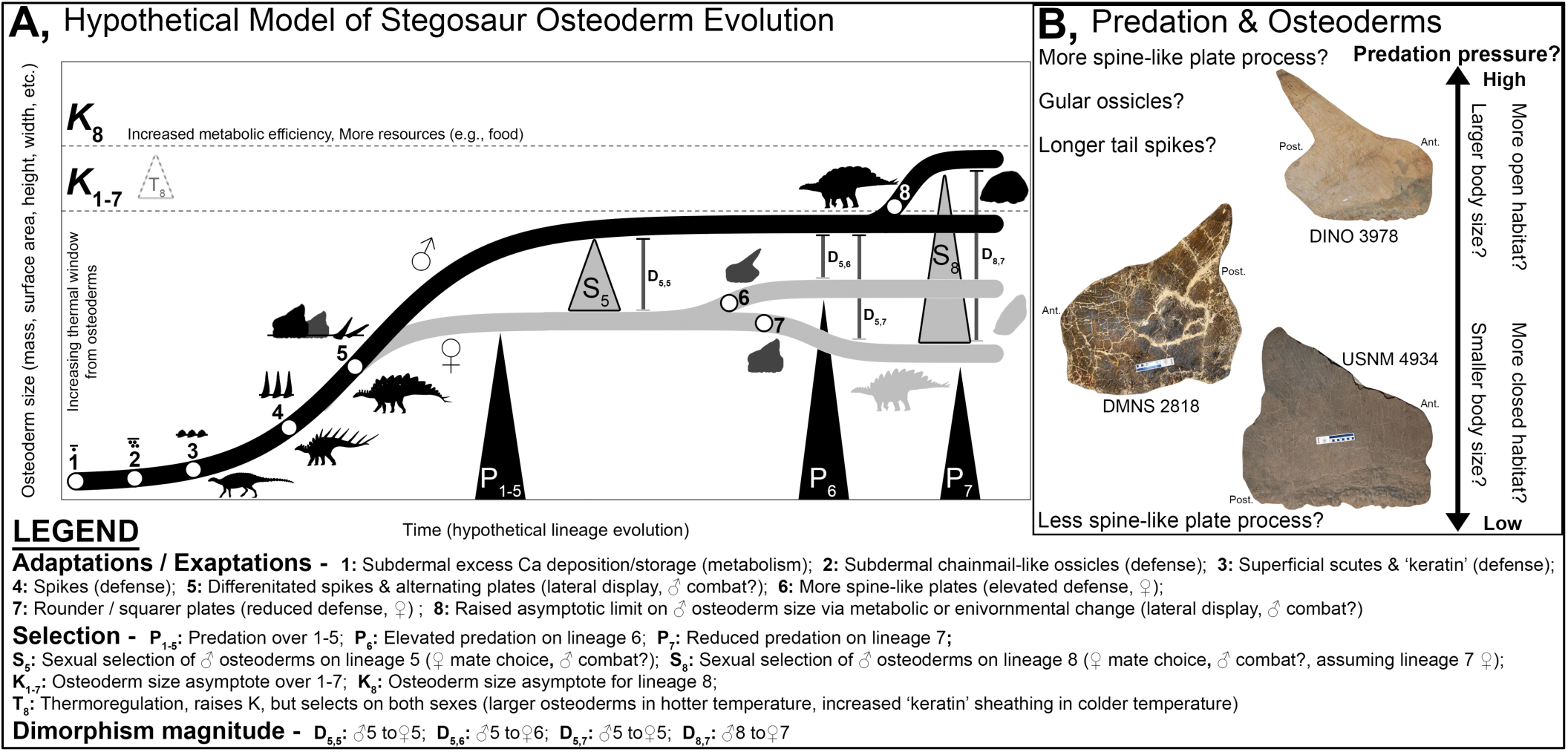
Summary of a hypothetical model for the evolution of stegosaur osteoderms and their functions. A, hypothesized interaction between osteoderm size, key adaptations/exaptations, selective pressures, and dimorphism magnitude between the sexes. B, the hypothesized impact of predation pressure on stegosaur osteoderms, including variation in spine-like projections on the plates, with three *Stegosaurus* plates shown as examples and scaled to each other in size. DMNS 2818 and USNM 4934 plates shown are the largest preserved plates on those respective individuals.

### Mineral storage

Although defense can be offered as an explanation for the origin of osteoderms, for sufficiently sized and dense structures to act as armor, an underlying mechanism of skin calcification must already be in place. One can hypothesize (Fig. 10A) that the ultimate origin of osteoderms could have derived from metabolic pathways that functioned to deposit and store excess calcium subdermally (point 1 in Fig. 10A). This hypothesis (Farlow *et al*. 2010) is consistent with mineral storage functions of osteoderms in modern organisms, especially female lizards and crocodilians which use mineral resorption to assist in eggshell formation (Dacke *et al*. 2015; Broeckhoven & du Plessis 2022). Even a study of osteoderm density in the viviparous armadillo lizard suggests that osteoderms might function as calcium reservoirs and that a ‘mineral storage hypothesis’ should be considered as a candidate to explain the evolution of osteoderms (Broeckhoven & du Plessis 2022). Among non-avian dinosaurs, titanosaur sauropod osteoderms have also been suggested to function in calcium storage for oogenesis (Vidal *et al*. 2017).

### Exaptation for defense

Over time, with increased size and coverage, these calcifications could have been exapted into defensive chainmail-like arrays of simple, subdermal osteoderms (point 2 in Fig. 10A). Osteoderm arrays are seen in organisms such as extinct ground sloths (Hill 2006) and some extant fish, frogs, lizards, snakes, and rodents (Broeckhoven 2022; Frýdlová *et al*. 2023; Maden *et al*. 2023). Analogous structures might also be seen in the gular ossicles of some *Stegosaurus* specimens/taxa. Since even highly successful modern predators target weak points (e.g., the throat) when hunting large prey and avoid horns and antlers (Murphy *et al*. 2009), it is reasonable to assume that gular ossicles of some *Stegosaurus* specimens/taxa protected against predatory bites to the throat.

Increasing predation pressure could lead to larger, more prominent osteoderms further adapted for defense. This selection would produce more superficially protruding, scute-like osteoderms covered by keratin that rose out and above the surrounding skin (point 3 in Fig. 10A). This state would be the condition seen in ankylosaurs and basal thyreophorans (Norman *et al*. 2004; Vickaryous *et al*. 2004), such as *Scutellosaurus* and *Scelidosaurus* (e.g., Thompson *et al*. 2011). Similar structures appear in many other organisms, such as titanosaurs (Vidal *et al*. 2017), *Ceratosaurus* (Hendrickx *et al*. 2022), crocodilians (Vickaryous & Hall 2008), lizards such as Cordylidae (Broeckhoven *et al*. 2018), and armadillos (Vickaryous & Hall 2006). Further selection could eventually lead to long, thin spikes (point 4 in Fig. 10A), ultimately taken to the extreme in stegosaurs such as *Kentrosaurus* (Galton 1982) and *Miragaia* (Mateus *et al*. 2009) that also possessed long shoulder spikes, as well as some nodosaurid ankylosaurs with prominent shoulder spikes, such as *Borealopelta* (Brown 2017).

Passively defensive ossicles in the skin can undergo selection towards use in more actively defensive behavior, especially those involving tail movements in reptiles. Sand boas are the only clade of snakes with species possessing dermal armor, like those of other squamates such as gerrhosaurids and geckos. Their hidden subdermal osteoderms in the tail are associated with modified, enlarged caudal vertebrae and anti-predator tail displays (Frýdlová *et al*. 2023). Agamid lizards use defensive tail whips (Schall *et al*. 1989), including the spiny-tailed lizard (*Uromastyx*) (Pianka & Vitt 2003). Extensively osteoderm-bearing cordylid lizards also engage in defensive tail whips, such as *Smaug*, which possesses large bony spikes on its tail (Fogel 2000). Ankylosaur tail clubs might also have undergone similar evolution.

Over time, stegosaur spikes became restricted to and specialized at the terminal tail as spike arrays (i.e., thagomizers) for use in specific behaviors. Alternatively, in ankylosaurs, flatter scutes were possibly modified directly into clubs (Arbour & Currie 2015; Soto-Acuña *et al*. 2021). Overall, osteoderms switch from passive defense against the teeth and claws of predators into offensive weapons of the tail that can be used against predators and even against intra-specific rivals.

### Exaptation for sexual combat/display

With the evolution of large, external osteoderms, their defensive function could be partly exapted into a sexual combat or display function under the influence of sexual selection (point 5 in Fig. 10A). Indeed, osteoderms in many extant and extinct vertebrates (e.g., fish, frogs, lizards, crocodilians, xenarthans, arthrodires, temnospondyls, glyptodonts, pareiasaurs, thyreophorans, and phytosaurs) are hypothesized to have evolved at least partly under sexual selection for intraspecific combat (i.e., ‘fighting-advantage hypothesis’), rather than defense alone (Broeckhoven 2022).

Defensive tail spikes in stegosaurs might have been used in intra-sex combat, like the tail clubs of ankylosaurs (Arbour *et al*. 2022). Meanwhile, the more anterior spikes of basal stegosaurs and other thyreophorans could be expanded into plate-like ‘splates’, as in polocanthine ankylosaurs (Blows 2001; Blows & Honeysett 2014). Eventually, ‘splates’ could widen into true plates to function in visual display. Of course, stegosaur plates used for lateral display in sexual contexts (e.g., males sizing each other up before combat) could also be used alongside the spikes in threat displays to deter predators by making the prey appear larger (Carpenter 1998). The evolution of true plates would have several implications. It would differentiate the most posterior osteoderms at the end of the tail from the more anterior osteoderms – becoming the thagomizer and plate row, respectively. Therefore, the term thagomizer is a useful scientific descriptor since it underscores the morphological, and thereby functional, differentiation of osteoderms in stegosaurs. A thagomizer is a functional module. Assuming that thagomizers were under sexual selection and used for intra-sexual combat, stegosaur behavior could include male combat before visual display to attract females, possibly in leks.

A number of modern species have prominent sagittal crests along the back which serve a sexual display function. Among lizards, males of various iguanas (e.g., *Iguana*) have dorsal crests elaborated by long scales (Dugan 1982). Other examples include the crested anole (*Anolis cristatellis*) (Prado-Irwin *et al*. 2019), basilisks (*Basiliscus*) (Taylor *et al*. 2017), sailfin lizards (*Hydrosaurus*) (Ord & Stuart-Fox 2006), and some chameleons (Eckhardt *et al*. 2012), which are typically better developed in males than females. Of course, this does not prove that stegosaur plates must have functioned in the same fashion, but the repeated evolution of dorsal crests for display, and the existence of sexual dimorphism in these structures, makes the hypothesized display function and hypothesized sexual dimorphism in stegosaurs more plausible.

*The color of stegosaur plates* – Unlike metabolically active skin, the ‘dead’ keratin covering of stegosaur plates and spikes might preclude the hypothesis that they visibly flushed with blood as part of a color-based, inter-or intra-specific display as suggested by Carpenter (1998). This might be especially true if a keratinous sheath around the osteoderms was very thick and extensive – more akin to horn sheaths in bovids than to the thinner integument cover on crocodilian osteoderms or on testudine carapaces/plasterons. We know of no extant organism that can blush thick, hard keratinous structures (e.g., rhamphotheca, horn sheaths, cranial casques) in such a manner. Presumably, the vascularity of plates is sufficiently consistent with their rapid growth upon sexual maturity as a secondary sexual characteristic (Hayashi *et al*. 2009) or for any thermoregulatory purposes (Farlow *et al*. 1976; de Buffrenil *et al*. 1986; Farlow *et al*. 2010).

Still, the conspicuousness of stegosaur plates raises questions as to any possible coloration or sexual dichromatism. Keratinous structures can be colored using pigments such as melanin or carotenoids (Roy *et al*. 2020) or using non-pigment-based structural color (Prum & Torres 2003). Toucans have brightly colored and pigmented rhamphotheca (Mora & López Umaña 2020), as do other birds (Davis & Clarke 2022). Reptiles also pigment their scales (Rutland *et al*. 2019), including the integument covering osteoderms, such as in the outermost corneous layer above crocodilian osteoderms (Alibardi 2011). Rather than changing blood flow, crocodilians can redistribute melanosomes in integumentary melanophores of the living dermis to rapidly change their color in response to environmental conditions (Merchant *et al*. 2018). This is not to say that the blood flow to the plates could not be physiologically altered, such as in vasoconstriction or vasodilation during thermoregulation (Andrade 2015; Tattersall 2016). Toucans can alter blood flow to their bill depending on ambient temperatures (Tattersall *et al*. 2009), and the same phenomenon is seen in cassowary casques (Eastick *et al*. 2019).

Although fossilized plate sheathing has been reported in some stegosaur specimens (e.g., the old adult, wide-morph *H. mjosi* SMA 0018 [Christiansen & Tschopp 2010]), no organic compounds have been reported. Nevertheless, the diagenetic stability of pigments such as melanins, carotenoids, and porphyrins (Roy *et al*. 2020) means that it may be possible to test hypotheses of stegosaur osteoderm coloration as more fossils are uncovered in the future.

*Asymmetry and chirality* – With display becoming a primary function of the plates, the model would predict that the appearance of offset staggering of osteoderms correlates with the appearance of large plates acting as a large, visual ‘billboard’. This prediction is so far matched by the fossil record of stegosaurs. Only the laterally compressed true plates of *Stegosaurus* and *H. mjosi* clearly lack exact matching pairs in size and shape, and highly articulated specimens directly preserve this staggering (e.g., BYU 12290, DMNS 2818, NHMUK PV R36730, USNM 4934). In contrast, the spikes within the thagomizer of these specimens/taxa remain paired. If the primitive condition of plates was originally mediolaterally paired, and the plates became asymmetrically offset during development, this might predict that a complete plate series consisted of an even number of plates. Indeed, the highest known plate series from a single individual consists of 18 preserved plates in NHMUK PV R36730. Although secondary sexual characteristics can show fluctuating asymmetry (which may not be under direct sexual selection [Balmford *et al*. 1993; Ditchkoff *et al*. 2001; Swaddle 2003]) and internal organs of animals are typically left–right asymmetrical (Blum & Ott 2018), such extreme and consistent external asymmetry/chirality in *Stegosaurus* and *H. mjosi* is only seen in some narrow cases among extant animals (Levin 1999; Palmer 2009). This is particularly true as it relates to structures under presumed sexual selection or that are sexually dimorphic: for example, the enlarged claw of male fiddler crabs used for intrasexual combat and intersexual display (Reaney *et al*. 2008); the asymmetric antlers of caribou, especially males whose tines might protect the eye during threshing displays (Pruitt 1966; Gross 1990); and the single enlarged erupting tusk of male narwhals on their left side that are likely used in male combat (Gerson & Hickie 1985; Nweeia *et al*. 2009). Variation in chiral handedness (i.e., left-versus right-handed) and degree of asymmetry in stegosaur plates across populations, sexes, and taxa is uncertain. The largest plate above the hips/base of the tail has been previously suggested to be on the right side based on USNM 4934 (and possibly DMNS 2818 and NHMUK PV R36730) (Cameron *et al*. 2016).

Based on early wind tunnel experiments, the staggered asymmetry of broad plates in *Stegosaurus* and *H. mjosi* would also have subsequently impacted the animal’s thermoregulation by allowing for forced convection by air passing around the plates under windy conditions (Farlow *et al*. 1976).

### Male and female evolutionary dynamics

With the exaptation of defensive structures into secondary sexual structures, male and female osteoderm size, shape, and relative functions will begin to diverge. The sex with the larger osteoderms is typically expected to be male (Summers & Ord 2022). In the males, sexual selection (*S* in Fig. 10A) from the opposite sex (mate choice) or within the sex (intrasexual competition) is modelled to drive up the size of osteoderms towards a physiological upper limit (carrying capacity, *K*, in Fig. 10A). Males would have large, broad plates for display and possibly long spikes in the thagomizer for combat. Sexual ornaments and armaments of extant animals are known to be driven by sexual selection. For example, in bovids, male horn length correlates negatively with male territoriality and positively with social group size, two variables that are suggested to impact the average number of mates per male (Bro-Jørgensen 2007).

Our model follows the concept of ‘runaway’ sexual selection in males (Chandler *et al*. 2013; Henshaw & Jones 2020). The physiological upper limit to this ‘runaway’ male osteoderm size is dictated by the metabolic efficiency in growing bone and keratin tissue, as well as food availability in the environment. The upper limit could also be influenced by other factors, such as thermoregulation. Larger, well-vascularized osteoderms mean larger ‘thermal windows’ (Andrade 2015; Tattersall 2016), which can facilitate heat exchange with the environment unless changes in relative keratin insulation are accounted for (as in bovid horns [Picard *et al*. 1994, 1996, 1999]). Therefore, changes in climate might have impacted the physiological upper limit for osteoderm size, especially in smaller-bodied stegosaur taxa. We might also predict that the male could have a larger body mass than the female when under selection for intrasexual combat.

The sex with the smaller osteoderms is typically expected to be female (Summers & Ord 2022). In females, the size of the plates and spikes is modeled here as being driven by predation pressure. The minimal size of spikes and plates required to deter predation adequately becomes expressed by the female. The plates in females are expected to be more spine-like in shape for ‘prickly’ predator deterrence compared to those of the male, akin to the more basal spiny condition of stegosaur osteoderms. In males, osteoderms would exceed the size of female osteoderms due to sexual selection. If predation pressure increases (point 6 in Fig. 10A) or decreases (point 7 in Fig. 10A) (e.g., due to more open or closed habitat, respectively), then osteoderm size may change in females accordingly. Females might also be expected to have a smaller body mass if they do not engage in intrasexual combat as extensively as do males.

These hypothesized evolutionary dynamics are inspired by observations of extant bovid horns. Although they are not osteoderms, horns are structures like thyreophoran spikes insofar as they are vascularized, conical bone covered by a keratin sheath. In bovids (Packer 1983; Caro *et al*. 2003; Stankowich & Caro 2009), male horns are adapted for intra-sex combat – they have broad bases with downward-pointing tips to resist lateral stress during non-lethal sparring. Female horns, when present, are adapted for defense against predators – they have thin bases with outward-pointing tips for goring. Analogies can be drawn to defensive body armor as well as offensive weaponry (e.g., horns). Sexual dimorphism occurs in body armor of cordyline lizard species that possess isolated osteoderms (as in many thyreophorans) rather than continuously imbricated osteoderms, and these dimorphic osteoderms lack obvious ecological primary functions (Broeckhoven *et al*. 2018). Thus, Broeckhoven *et al*. (2018) suggest that ecological selective pressures (e.g., defense against predators) can reduce dimorphism magnitude in osteoderms, even if sexual selection is pervasive across a clade.

*Sexual dimorphism magnitude* – The difference in osteoderm size and shape between the two sexes in our hypothetical model is the magnitude of sexual dimorphism. This magnitude is therefore a consequence of 1) sexual selection driving males towards the physiological upper limit of osteoderm size and 2) predation pressure dictating the minimum adaptive osteoderm size in females. While the early ontogeny of and extent of sexual variation in thyreophoran osteoderms remain poorly known in most of the clade, no thyreophoran specimen has been reported to confidently demonstrate an absence of osteoderms *in vivo*. So far, specimens without associated osteoderms are consistent with incomplete preservation of the skeleton rather than an *in vivo* absence of osteoderms. The presence of osteoderms is even used as a synapomorphy of the clade (Norman *et al*. 2004). It is therefore reasonable to predict that thyreophorans experienced consistent predation pressure to some degree. A similar pattern is seen in extant bovid horns and possibly in the ossicones and antlers of other extant mammals. Among these extant species, those that live in open habitats and that are more conspicuous due to larger body size (i.e., they cannot rely on crypsis) are more likely to have females with horns for defense against predators; furthermore, compared to species with hornless females, horned females are more likely to be territorial and to engage in intrasexual competition (via sparring) (Roberts 1996; Stankowich & Caro 2009). Thus, some amount of territoriality and intrasexual competition between female stegosaurs, in addition to competition between males, may be plausible.

*Predation and stegosaur osteoderms* – The impact of predation pressure on osteoderm evolution can be hypothesized (Fig. 10B) based on our understanding of horn dimorphism patterns in extant bovids. In extant bovids, females of large taxa in open habitats (e.g., poorly vegetated) are more likely to express horns for defense than females of small taxa in closed habitats (Roberts 1996; Stankowich & Caro 2009). Stegosaur taxa with lower dimorphism magnitude might occur in open environments where large-bodied animals cannot effectively hide from predators. These stegosaur taxa might also be hypothesized to show more spine-like processes in their plates, gular ossicles to protect the throat, and longer tail spikes. In contrast, small stegosaur taxa in closed environments (e.g., highly vegetated) might have higher dimorphism magnitude, bear rounder or more boxy plates with less-pronounced spine-like processes, may be less likely to possess gular ossicles, and may have shorter tail spikes.

‘*Thermal windows’* – Increasing osteoderm size would increase the size of the potential ‘thermal window’ (Andrade 2015; Tattersall 2016), facilitating heat exchange with the environment. Extant lizards show a tradeoff between the strength of osteoderms as it relates to defense and their thermal capacity to exchange heat as osteoderm vascularity increases (Broeckhoven *et al*. 2017a). The high vascularity of stegosaur osteoderms, particularly their plates, may therefore be weighted toward heat exchange over biomechanical strength. In small stegosaur taxa especially, their body’s surface area to volume ratios may be high enough that they must account for this osteoderm ‘thermal window’ to maintain a proper body temperature. This aspect of our model is inspired by observations of extant bovid horns, whereby taxa from low latitudes and warm climates transported to higher latitude zoos can develop frostbite on their horns – this is due to a lower ratio of insulative external keratin sheathing to vascularized internal bone cores compared taxa from colder climates (Kitchener 1991; Picard *et al*. 1994, 1996, 1999). Therefore, in warmer environments and lower latitudes, we predict stegosaurs (of both sexes) were possibly under selection (*T* in Fig. 10A) for larger osteoderms, lower keratin-to-bone ratios in their osteoderms, and/or increased osteoderm vascularity (which may reduce their biomechanical strength as defensive structures). Stegosaur taxa in colder environments and higher latitudes would be expected to show the inverse trend. Therefore, in addition to factors such as food availability and metabolic efficiency of bone/keratin growth, temperature might also help to raise (point 8 in Fig.10A) or lower the upper limit on osteoderm size.

### *H. mjosi* within the model

Within our hypothetical model, *H. mjosi* would be expected to fall at the more extreme ends of dimorphism magnitude (e.g., D_8,7_ in Fig. 10A). It is so far the only stegosaur taxon with high-confidence evidence for large-magnitude sexual dimorphism in plates – with a 45% difference in maximum single-plate surface area between the two hypothesized sexes. This conclusion is consistent with the small body size of *H. mjosi* (estimated 6.5 m body length [Siber & Möckli 2009]) compared to some species of *Stegosaurus* (estimated 9 m body length in largest *Stegosaurus* species [Galton & Upchurch 2004]).

As part of a possibly unique northern Morrison biota (Whitlock *et al*. 2018; Foster 2020), *H. mjosi* likely inhabited higher latitudes compared to most *Stegosaurus*. The northern Morrison was a wetter and more heavily vegetated environment, consisting of boreal forest with an herbaceous and fern undergrowth (Richmond 2023). This could suggest that *H. mjosi* experienced lower predation pressure, since smaller size and a more closed habitat predicts lower predation risk in some extant large herbivores (Roberts 1996; Stankowich & Caro 2009). Low predation risk would allow females to invest less heavily in osteoderms. Females would be predicted to bear tall plates that, although still more consistent with defense than those of the hypothesized male, were relatively rounded compared to the more rectangular, triangular, or spine-like plates in *Stegosaurus* (e.g., Fig. 10B). Furthermore, hypothesized male *H. mjosi* might have had a high upper limit on osteoderm growth if a wetter environment led to plentiful vegetation for food. Males would be predicted to bear wide, broad plates that were highly optimized for visual ‘billboard’ display more so than ‘prickly’ defense. Low predation pressure is further consistent with the apparent lack of gular ossicles in *H. mjosi*. Considering all of these factors, we would predict maximal sexual dimorphism magnitude in this taxon.

The habitat considerations for *H. mjosi* would also be consistent with the hypothesis proposed by Maidment *et al*. (2018) that larger-bodied *Stegosaurus* 1) would have a lower metabolism per mass compared to *H. mjosi*, requiring less food on a per mass basis, and 2) would have longer limbs to cover ground more efficiently across more arid environments where food is scarcer or more widely dispersed. It is also interesting that in extant African antelopes, only about half of the genera have horn-bearing females and these genera are usually heavier (Packer 1983). The correlation between body mass and armamented females in bovids is consistent with the hypothesis that heavier *Stegosaurus* species would show lesser magnitudes of sexual dimorphism than the lighter *H. mjosi*. Across stegosaur taxa, greater magnitudes of dimorphism in species like *H. mjosi* could be hypothesized to correlate with 1) lekking, 2) larger and less territorial social groups (Bro-Jørgensen 2007), and 3) perhaps polygyny, given the interspecific variation in thyreophoran osteoderms. Indeed, *H. mjosi* have been found in multi-individual mass death assemblages consistent with sociality (JRDI 5ES Quarry). Predicted polygyny would be consistent with patterns seen in extant bovids, where morphological disparity of male horns is greater between polygynous species than between monogamous species (Caro *et al*. 2003) – North American stegosaurs show high variability in plate shape, even in the underlying bone core without accounting for keratin sheathing.

## CONCLUSIONS

When evaluating hypotheses for stegosaur osteoderm evolution and function, considering multiple functions is helpful. These functions include sexual display/combat, defense, and thermoregulation. Their relative importance could vary across time and space, between sexes, and across different regions of the body (e.g., gular ossicles, plates, and thagomizers). As such, reexamining the evidence for sexual dimorphism in the plates of *H. mjosi* is worthwhile.

Various statistical and taphonomic evidence contradicts the alternative hypothesis that plate variation in *H. mjosi* is due primarily or entirely to different positions anteroposteriorly along the body. Both hypothesized morphs of *H. mjosi* plates appear to come from cervical, dorsal, and caudal regions, and taphonomically isolated individuals appear to show the presence of only one morph among all their associated plates. Furthermore, highly precise two-dimensional representations of plate shape variation using outline analysis are consistent with two distinguishable morphs of wide versus tall plates. In our sample, likely full of plates from sexually mature adults, an appreciable effect between the shape and maximum size of the two plate morphs is supported with high confidence as they diverge at larger plate sizes. However, *H. mjosi* provides a challenging case study, given the violations of assumptions of independence among the datapoints (i.e., multiple plates can come from a single individual in the dataset). When examining differences between small and large plates, ontogenetic variation can confound with intra-individual anteroposterior variation. Still, even in the presence of appreciable intra-individual anteroposterior variation in plate size and shape, the data is currently consistent with sexual dimorphism. Additional *H. mjosi* specimens may further contribute support to the sexual dimorphism hypothesis as more fossils are collected.

However, questions of sexual dimorphism in stegosaur body mass and tail spikes remain. If wide-plated *H. mjosi* are hypothetically male, do they also reach larger body masses than the opposite sex, in addition to larger plate sizes? Increased body mass dimorphism correlates positively with polygyny (Pérez-Barbería *et al*. 2002; Cassini 2020). Might wide-plated *H. mjosi* also have longer tail spikes than the hypothesized females? Pathologies in tail spikes of *Stegosaurus* and caudal vertebrae of ontogenetically old, wide-morph *H. mjosi* could be consistent with intra-sexual combat, not just defense against predators. It is not unusual for males to first compete against each other before displaying to attract females.

We propose a model of stegosaur osteoderm evolution that originates from calcium storage and is exapted into defensive ossicles, scutes, and spikes. Further exaptation through sexual selection is predicted to correlate with 1) the differentiation of plates from the thagomizer and 2) the unusual chiral asymmetry in the two rows of staggered plates in *Stegosaurus* and *H. mjosi*. Plates are likely adapted for visual display, while spikes are possibly for male combat and defense. Male osteoderms are then driven up to a physiological/environmental upper size limit, while female osteoderm size is determined by predation pressure. More intense predation pressure may correlate with larger-bodied taxa, more open habitat, more spine-like processes in plates, the presence of gular ossicles, and/or longer tail spikes. Together, sexual selection, predation pressure, and even thermoregulatory demands could work to increase or decrease the magnitude of sexual dimorphism in osteoderms. In our hypothetical model, *H. mjosi* would be predicted to show high levels of sexual dimorphism in osteoderms. The prediction is based on its 1) smaller size, 2) wetter, more vegetated habitat (a boreal forest with an herbaceous and fern undergrowth [Richmond 2003]), 3) apparent lack of gular ossicles, and 4) rounder plates than in *Stegosaurus*. *Stegosaurus*, on the other hand, would have reached larger sizes, lived in lower latitudes and drier climates, and had more triangular, rectangular, or spine-like plates (as well as gular ossicles in some instances).

The question of stegosaur plate function is one of the most intriguing and longest standing in paleontology. However, the solution requires us to consider complex, multifactor evolutionary dynamics as well as primary versus secondary functions. Overall, it seems that thyreophoran osteoderms adapted for defense became coopted for sexual display and combat in later stegosaurs and ankylosaurs.

## ACKNOWLEDGEMENTS

We thank James O. Farlow (Purdue University Fort Wayne) for helpful comments. University of Chicago crews led by Paul C. Sereno collected the specimen UCRC PV6. Kenneth Carpenter provided images of HMNS 14. We thank Nate L. Murphy (JRDI), Jim Shaffer, and Dave & Rosalie Hein for their assistance in obtaining data from the JRDI specimens and for their helpful feedback. We also thank M. Murphy and JRDI volunteers for their excavation and preparation efforts. Bill Wahl assisted in measuring some of the bones at WDC. Thomas Bolliger (SMA) and Hans-Jakob “Kirby” Siber (SMA) assisted E. T. Saitta during a research visit to SMA, and they also provided helpful information about the transfer of SMA specimens to NMZ along with Dennis Hansen (Natural History Museum, University of Zurich). We also thank the many individuals who allowed access to the specimens at all the repositories (see also the acknowledgements in Saitta [2014]). *Stegosaurus* (Scott Hartman, Creative Commons Attribution 3.0 Unported [CC BY 3.0]), *Kentrosaurus* (Emily Willoughby, Attribution-ShareAlike 3.0 Unported [CC BY-SA 3.0]), and *Scelidosaurus* (Matthew Dempsey, Attribution 4.0 International [CC BY 4.0]) silhouettes were obtained through phylopic.org.

## LITERATURE CITED

1. Ackermann, R. R., Brink, J. S., Vrahimis, S. and De Klerk, B., 2010. Hybrid wildebeest (Artiodactyla: Bovidae) provide further evidence for shared signatures of admixture in mammalian crania. South African Journal of Science, 106(11), pp.1–5.

2. Ackermans, N. L., Varghese, M., Williams, T. M., Grimaldi, N., Selmanovic, E., Alipour, A., Balchandani, P., Reidenberg, J. S. and Hof, P. R., 2022. Evidence of traumatic brain injury in headbutting bovids. Acta neuropathologica, 144(1), pp.5–26.

3. Alatalo, R. V., Höglund, J. and Lundberg, A., 1991. Lekking in the black grouse—a test of male viability. Nature, 352(6331), pp.155–156.

4. Albersdörfer, R., and Galiano, H. (2009). Dinosaurs from Dana Quarry. Ten Sleep, WY, USA: Dinosauria International, LLC.

5. Alibardi, L., 2011. Histology, ultrastructure, and pigmentation in the horny scales of growing crocodilians. Acta Zoologica, 92(2), pp.187–200.

6. Alleon, J., Montagnac, G., Reynard, B., Brulé, T., Thoury, M. and Gueriau, P., 2021. Pushing Raman spectroscopy over the edge: purported signatures of organic molecules in fossil animals are instrumental artefacts. BioEssays, 43(4), p.2000295.

7. Amrhein, V., Greenland, S. and McShane, B., 2019. Scientists rise up against statistical significance. Nature, 567(7748), pp.305–307.

8. Andrade, D. V., 2015. Thermal windows and heat exchange. Temperature, 2(4), pp.451–451.

9. Apollonio, M., Festa-Bianchet, M., Mari, F., Mattioli, S. and Sarno, B., 1992. To lek or not to lek: mating strategies of male fallow deer. Behavioral Ecology, 3(1), pp.25–31.

10. Arbour, V. M., 2009. Estimating impact forces of tail club strikes by ankylosaurid dinosaurs. PLoS One, 4(8), p.e6738.

11. Arbour, V. M. and Currie, P. J., 2011. Tail and pelvis pathologies of ankylosaurian dinosaurs. Historical Biology, 23(4), pp.375–390.

12. Arbour, V.M. and Currie, P.J., 2015. Ankylosaurid dinosaur tail clubs evolved through stepwise acquisition of key features. Journal of Anatomy, 227(4), pp.514–523.

13. Arbour, V. M., Zanno, L. E. and Evans, D. C., 2022. Palaeopathological evidence for intraspecific combat in ankylosaurid dinosaurs. Biology Letters, 18(12), p.20220404.

14. Balmford, A., Jones, I. L. and Thomas, A. L., 1993. On avian asymmetry: evidence of natural selection for symmetrical tails and wings in birds. Proceedings of the Royal Society of London. Series B: Biological Sciences, 252(1335), pp.245–251.

15. Barden, H. E., and Maidment, S. C., 2011. Evidence for sexual dimorphism in the stegosaurian dinosaur *Kentrosaurus aethiopicus* from the Upper Jurassic of Tanzania. Journal of Vertebrate Paleontology, 31(3), pp.641–651.

16. Barrette, C., 1986. Fighting behavior of wild *Sus scrofa*. Journal of Mammalogy, 67(1), pp.177–179.

17. Benton, M. J., 2021. The origin of endothermy in synapsids and archosaurs and arms races in the Triassic. Gondwana Research, 100, pp.261–289.

18. Besharse, J.C., 1971. Maturity and sexual dimorphism in the skull, mandible, and teeth of the beaked whale, *Mesoplodon densirostris*. Journal of Mammalogy, 52(2), pp.297–315.

19. Blows, W. T. 2001. Dermal armor of the polacanthine ankylosaurs. In: K. Carpenter (ed.), The Armored Dinosaurs, 363–385. Indiana University Press, Bloomington

20. Blows, W. T. and Honeysett, K., 2014. First Valanginian Polacanthus foxii (Dinosauria, Ankylosauria) from England, from the Lower Cretaceous of Bexhill, Sussex. *Proceedings of the Geologists’* Association, 125(2), pp.233–251.

21. Blum, M. and Ott, T., 2018. Animal left–right asymmetry. Current Biology, 28(7), pp.R301–R304.

22. Borkovic B, Russell A. Sexual selection according to Darwin: A response to Padian and Horner’s interpretation. Comptes Rendus Palevol. 2014 Nov 1;13(8):701–7.

23. Bonhomme, V., Picq, S., Gaucherel, C., and J. Claude. 2014. Momocs: Outline analysis using R. Journal of Statistical Software 56:1–24.

24. Briggs, K. and Morejohn, G.V., 1975. Sexual dimorphism in the mandibles and canine teeth of the northern elephant seal. Journal of Mammalogy, 56(1), pp.224–231.

25. Broeckhoven, C., 2022. Intraspecific competition: A missing link in dermal armour evolution?. Journal of Animal Ecology, 91(8), pp.1562–1566.

26. Broeckhoven, C. and du Plessis, A., 2022. Osteoderms as calcium reservoirs: Insights from the lizard *Ouroborus cataphractus*. Journal of Anatomy, 241(3), pp.635–640.

27. Broeckhoven, C., Diedericks, G. and Mouton, P. L. F. N., 2015. What doesn’t kill you might make you stronger: functional basis for variation in body armour. Journal of Animal Ecology, 84(5), pp.1213–1221.

28. Broeckhoven, C., de Kock, C. and Mouton, P. L. F. N., 2017b. Sexual dimorphism in osteoderm expression and the role of male intrasexual aggression. Biological Journal of the Linnean Society, 122(2), pp.329–339.

29. Broeckhoven, C., du Plessis, A. and Hui, C., 2017a. Functional trade-off between strength and thermal capacity of dermal armor: insights from girdled lizards. Journal of the Mechanical Behavior of Biomedical Materials, 74, pp.189–194.

30. Broeckhoven, C., De Kock, C. and Hui, C., 2018. Sexual dimorphism in the dermal armour of cordyline lizards (Squamata: Cordylinae). Biological Journal of the Linnean Society, 125(1), pp.30–36.

31. Bro-Jørgensen, J., 2007. The intensity of sexual selection predicts weapon size in male bovids. Evolution, 61(6), pp.1316–1326.

32. Bro-Jørgensen, J. and Durant, S. M., 2003. Mating strategies of topi bulls: getting in the centre of attention. Animal Behaviour, 65(3), pp.585–594.

33. Brown, C.M., 2017. An exceptionally preserved armored dinosaur reveals the morphology and allometry of osteoderms and their horny epidermal coverings. PeerJ, 5, p.e4066.

34. Cameron, R. P., Cameron, J. A. and Barnett, S. M., 2016. Stegosaurus chirality. arXiv preprint arXiv:1611.08760.

35. Caro, T. M., Graham, C. M., Stoner, C. J. and Flores, M. M., 2003. Correlates of horn and antler shape in bovids and cervids. Behavioral Ecology and Sociobiology, 55, pp.32–41.

36. Carpenter, K., 1998. Armor of *Stegosaurus stenops*, and the taphonomic history of a new specimen from Garden Park, Colorado. Modern Geology, 23, pp.127–144.

37. Carpenter, K., Miles, C. A. and Cloward, K., 2001. New primitive stegosaur from the Morrison Formation, Wyoming. The Armored Dinosaurs, pp.55-75.

38. Carpenter, K., Sanders, F., McWhinney, L. A. and Wood, L., 2005. Evidence for predator-prey relationships: examples for Allosaurus and Stegosaurus. The Carnivorous Dinosaurs, pp. 325–350.

39. Casey, C., Charrier, I., Mathevon, N. and Reichmuth, C., 2015. Rival assessment among northern elephant seals: evidence of associative learning during male–male contests. Royal Society Open Science, 2(8), p.150228.

40. Cassini, M. H., 2020. A mixed model of the evolution of polygyny and sexual size dimorphism in mammals. Mammal Review, 50(1), pp.112–120.

41. Chandler, C. H., Ofria, C. and Dworkin, I., 2013. Runaway sexual selection leads to good genes. Evolution, 67(1), pp.110–119.

42. Chapman, R. E., Galton, P. M., Sepkoski Jr, J. J. and Wall, W. P., 1981. A morphometric study of the cranium of the pachycephalosaurid dinosaur *Stegoceras*. Journal of Paleontology, pp.608–618.

43. Christiansen, N. A. and Tschopp, E., 2010. Exceptional stegosaur integument impressions from the Upper Jurassic Morrison Formation of Wyoming. Swiss Journal of Geosciences, 103, pp.163–171.

44. Chu, P.C., 1995. Phylogenetic reanalysis of Strauch’s osteological data set for the Charadriiformes. The Condor, 97(1), pp.174–196.

45. Costa, F. and Mateus, O., 2019. Dacentrurine stegosaurs (Dinosauria): A new specimen of *Miragaia longicollum* from the Late Jurassic of Portugal resolves taxonomical validity and shows the occurrence of the clade in North America. PloS one, 14(11), p.e0224263.

46. Cruzado-Caballero, P., Filippi, L.S., González-Dionis, J. and Canudo, J.I., 2023. How common are lesions on the tails of sauropods? Two new pathologies in titanosaurs from the late cretaceous of argentine patagonia. Diversity, 15(3), p.464.

47. Dacke, C. G., Elsey, R. M., Trosclair III, P. L., Sugiyama, T., Nevarez, J. G. and Schweitzer, M. H., 2015. Alligator osteoderms as a source of labile calcium for eggshell formation. Journal of Zoology, 297(4), pp.255–264.

48. Davison, G.W.H., 1985. Avian spurs. Journal of Zoology, 206(3), pp.353–366.

49. Davis, S. N. and Clarke, J. A., 2022. Estimating the distribution of carotenoid coloration in skin and integumentary structures of birds and extinct dinosaurs. Evolution, 76(1), pp.42–57.

50. Dawkins, R. and Krebs, J. R., 1979. Arms races between and within species. Proceedings of the Royal Society of London. Series B. Biological Sciences, 205(1161), pp.489–511.

51. de Buffrenil, V., Farlow, J. O. and de Ricqlès, A., 1986. Growth and function of *Stegosaurus* plates: evidence from bone histology. Paleobiology, 12(4), pp.459–473.

52. de Magalhães, J. P., 2023. The longevity bottleneck hypothesis: Could dinosaurs have shaped ageing in present-day mammals?. *Bioessays: News and Reviews in Molecular*, Cellular and Developmental Biology, pp.e2300098–e2300098.

53. Desta, T.T., 2019. Phenotypic characteristic of junglefowl and chicken. World’s Poultry Science Journal, 75(1), pp.69–82.

54. Deutsch, C. J., Haley, M. P. and Le Boeuf, B. J., 1990. Reproductive effort of male northern elephant seals: estimates from mass loss. Canadian Journal of Zoology, 68(12), pp.2580–2593.

55. Ditchkoff, S. S., Lochmiller, R. L., Masters, R. E., Starry, W. R. and Leslie Jr, D. M., 2001. Does fluctuating asymmetry of antlers in white–tailed deer (Odocoileus virginianus) follow patterns predicted for sexually selected traits?. Proceedings of the Royal Society of London. Series B: Biological Sciences, 268(1470), pp.891–898.

56. Dodson, P., 1976. Quantitative aspects of relative growth and sexual dimorphism in *Protoceratops*. Journal of Paleontology, pp.929–940.

57. Dugan, B., 1982. A field study of the headbob displays of male green iguanas (*Iguana iguana*): variation in form and context. Animal behaviour, 30(2), pp.327–338.

58. Eagle, R. A., Tütken, T., Martin, T. S., Tripati, A. K., Fricke, H. C., Connely, M., Cifelli, R. L. and Eiler, J. M., 2011. Dinosaur body temperatures determined from isotopic (^13^C-^18^O) ordering in fossil biominerals. Science, 333(6041), pp.443–445.

59. Eastick, D. L., Tattersall, G. J., Watson, S. J., Lesku, J. A. and Robert, K. A., 2019. Cassowary casques act as thermal windows. Scientific Reports, 9(1), p.1966.

60. Ebel, R., Herrel, A., Scheyer, T.M. and Keogh, J.S., 2024. Review of osteoderm function and future research directions. Journal of Zoology.

61. Eckhardt, F.S., Gehring, P.S., Bartel, L., Bellmann, J., Beuker, J., Hahne, D., Korte, J., Knittel, V., Mensch, M., Nagel, D. and Pohl, M., 2012. Assessing sexual dimorphism in a species of Malagasy chameleon (Calumma boettgeri) with a newly defined set of morphometric and meristic measurements. Herpetology Notes, 5, pp.335–344.

62. Eens, M. and Pinxten, R., 2000. Sex-role reversal in vertebrates: behavioural and endocrinological accounts. Behavioural Processes, 51(1-3), pp.135–147.

63. Elbroch, L. M., Feltner, J. and Quigley, H. B., 2017. Stage-dependent puma predation on dangerous prey. Journal of Zoology, 302(3), pp.164–170.

64. Farke, A. A., 2014. Evaluating combat in ornithischian dinosaurs. Journal of Zoology, 292(4), pp.242–249.

65. Farke, Andrew A., Ewan DS Wolff, and Darren H. Tanke. Evidence of combat in *Triceratops*. PLoS One 4, no. 1 (2009): e4252.

66. Farlow, J. O. and Dodson, P., 1975. The behavioral significance of frill and horn morphology in ceratopsian dinosaurs. Evolution, pp.353–361.

67. Farlow, J. O., Thompson, C. V. and Rosner, D. E., 1976. Plates of the dinosaur *Stegosaurus*: forced convection heat loss fins?. Science, 192(4244), pp.1123–1125.

68. Farlow, J. O., Hayashi, S. and Tattersall, G. J., 2010. Internal vascularity of the dermal plates of *Stegosaurus* (Ornithischia, Thyreophora). Swiss Journal of Geosciences, 103, pp.173–185.

69. Fogel, G., 2000. Observations on the Giant Sungazer Lizard, Cordylus giganteus, in captivity. Bull. Chic. Herpetol. Soc, 35, pp.277-280.

70. Foster, J. R., 2003. Paleoecological Analysis of the Vertebrate Fauna of the Morrison Formation (Upper Jurassic), Rocky Mountain Region, USA: Bulletin 23 (Vol. 23). New Mexico Museum of Natural History and Science.

71. Foster, J., 2020. Jurassic West: the dinosaurs of the Morrison Formation and their world. Indiana University Press, Bloomington.

72. Frýdlová, P., Janovská, V., Mrzílková, J., Halašková, M., Riegerová, M., Dudák, J., Tymlová, V., Žemlička, J., Zach, P. and Frynta, D., 2023. The first description of dermal armour in snakes. Scientific Reports, 13(1), p.6405.

73. Galton, P. M. 1982. The postcranial anatomy of stegosaurian dinosaur *Kentrosaurus* from the Upper Jurassic of Tanzania, East Africa. Geologica et Palaeontologica 15: 139–160.

74. Galton, P. M., & Upchurch, P. (2004). “Stegosauria”. In Weishampel, D. B., Dodson, P., & Osmolska, H. (eds.) The Dinosauria. Berkeley: Berkeley University Press, 343–362.

75. Gates, T. A., 2005. The Late Jurassic Cleveland-Lloyd dinosaur quarry as a drought-induced assemblage. Palaios, 20(4), pp.363–375.

76. Gerson, H. B. and Hickie, J. P., 1985. Head scarring on male narwhals (Monodon monoceros): evidence for aggressive tusk use. Canadian Journal of Zoology, 63(9), pp.2083–2087.

77. Gerstenhaber, C. and Knapp, A., 2022. Sexual selection leads to positive allometry but not sexual dimorphism in the expression of horn shape in the blue wildebeest, Connochaetes taurinus. BMC Ecology and Evolution, 22(1), p.107.

78. Gilmore, C. W., 1914. Osteology of the armored Dinosauria in the United States National Museum: with special reference to the genus Stegosaurus (No. 89). US Government Printing Office.

79. Grady, J. M., Enquist, B. J., Dettweiler-Robinson, E., Wright, N. A. and Smith, F. A., 2014. Evidence for mesothermy in dinosaurs. Science, 344(6189), pp.1268–1272.

80. Green, T. L., Kay, D. I. and Gignac, P. M., 2022. Intraspecific variation and directional casque asymmetry in adult southern cassowaries (*Casuarius casuarius*). Journal of Anatomy, 241(4), pp.951–965.

81. Grigg, G., Nowack, J., Bicudo, J. E. P. W., Bal, N. C., Woodward, H. N. and Seymour, R. S., 2022. Whole-body endothermy: ancient, homologous and widespread among the ancestors of mammals, birds and crocodylians. Biological Reviews, 97(2), pp.766–801.

82. Gosling, L. M., Petrie, M. and Rainy, M. E., 1987. Lekking in topi: A high cost, specialist strategy. Animal Behaviour, 35(2), 616–618.

83. Goss, R. J., 1990. Interactions between asymmetric brow tines in caribou and reindeer antlers. Canadian journal of zoology, 68(6), pp.1115–1119.

84. Haley, M. P., Deutsch, C. J. and Le Boeuf, B. J., 1994. Size, dominance and copulatory success in male northern elephant seals, Mirounga angustirostris. Animal Behaviour, 48(6), pp.1249–1260.

85. Hayashi, S., Carpenter, K. and Suzuki, D., 2009. Different growth patterns between the skeleton and osteoderms of *Stegosaurus* (Ornithischia: Thyreophora). Journal of Vertebrate Paleontology, 29(1), pp.123–131.

86. Hayashi, S., Carpenter, K., Watabe, M. and McWhinney, L. A., 2012. Ontogenetic histology of *Stegosaurus* plates and spikes. Palaeontology, 55(1), pp.145–161.

87. Hieronymus, T.L., Witmer, L.M., Tanke, D.H. and Currie, P.J., 2009. The facial integument of centrosaurine ceratopsids: morphological and histological correlates of novel skin structures. The Anatomical Record: Advances in Integrative Anatomy and Evolutionary Biology: Advances in Integrative Anatomy and Evolutionary Biology, 292(9), pp.1370–1396.

88. Henderson, D.M., 1999. Estimating the masses and centers of mass of extinct animals by 3-D mathematical slicing. Paleobiology, 25(1), pp.88–106.

89. Hendrickx, C., Bell, P.R., Pittman, M., Milner, A.R., Cuesta, E., O’Connor, J., Loewen, M., Currie, P.J., Mateus, O., Kaye, T.G. and Delcourt, R., 2022. Morphology and distribution of scales, dermal ossifications, and other non-feather integumentary structures in non-avialan theropod dinosaurs. Biological Reviews, 97(3), pp.960–1004.

90. Henshaw, J. M. and Jones, A. G., 2020. Fisher’s lost model of runaway sexual selection. Evolution, 74(2), pp.487–494.

91. Heurich, M., Zeis, K., Küchenhoff, H., Müller, J., Belotti, E., Bufka, L. and Woelfing, B., 2016. Selective predation of a stalking predator on ungulate prey. PloS one, 11(8), p.e0158449.

92. Hill, R. V., 2006. Comparative anatomy and histology of xenarthran osteoderms. Journal of Morphology, 267(12), pp.1441–1460.

93. Höglundi, J., Kålås, J. A. and Fiske, P., 1992. The costs of secondary sexual characters in the lekking great snipe (*Gallinago media*). Behavioral Ecology and Sociobiology, 30, pp.309–315.

94. Holthaus, K. B., Eckhart, L., Dalla Valle, L. and Alibardi, L., 2018. Evolution and diversification of corneous beta-proteins, the characteristic epidermal proteins of reptiles and birds. Journal of Experimental Zoology Part B: Molecular and Developmental Evolution, 330(8), pp.438–453.

95. Hone, D. W. and Naish, D., 2013. The ‘species recognition hypothesis’ does not explain the presence and evolution of exaggerated structures in non-avialan dinosaurs. Journal of Zoology, 290(3), pp.172–180.

96. Hone, D. W., Naish, D. and Cuthill, I. C., 2012. Does mutual sexual selection explain the evolution of head crests in pterosaurs and dinosaurs?. Lethaia, 45(2), pp.139–156.

97. Hone, D. W., Wood, D. and Knell, R. J., 2016. Positive allometry for exaggerated structures in the ceratopsian dinosaur Protoceratops andrewsi supports socio-sexual signaling. Palaeontologia Electronica, 19(1), pp.1–13.

98. Humphreys, R. K. and Ruxton, G. D., 2020. The dicey dinner dilemma: Asymmetry in predator–prey risk-taking, a broadly applicable alternative to the life-dinner principle. Journal of evolutionary biology, 33(3), pp.377–383.

99. Isles, T. E., 2009. The socio-sexual behaviour of extant archosaurs: implications for understanding dinosaur behaviour. Historical Biology, 21(3-4), pp.139–214.

100. Kitchener, A. C., 1991. The evolution and mechanical design of horns and antlers. Pages 229–253 in J. M. V. Rayner & R. J. Wooton (ed.). Biomechanics in Evolution. Cambridge University Press, Cambridge, Massachusetts.

101. Knell, R. J. and Sampson, S. D., 2011. Bizarre structures in dinosaurs: species recognition or sexual selection? A response to Padian and Horner. Journal of Zoology, 283, pp.18–22.

102. Knell, R. J., Naish, D., Tomkins, J. L. and Hone, D. W., 2013. Is sexual selection defined by dimorphism alone? A reply to Padian and Horner. Trends in Ecology & Evolution, 28(5), pp.250–251.

103. Larson, P. L., 2008. Variation and sexual dimorphism in Tyrannosaurus rex. Tyrannosaurus rex, pp.103-130.

104. Lategano FR, Conti SI, Lozar FR. MIRAGAIA TAIL BIOMECHANICS AND DEFENCES. EVALUATION OF THE TAIL MOBILITY AND RESISTANCE TO LOADINGS AND COLLISIONS. Rivista Italiana di Paleontologia e Stratigrafia. 2024;130(2):475–86.

105. Leboeuf, B. J., 1972. Sexual behavior in the northern elephant seal *Mirounga angustirostris*. Behaviour, 41(1-2), pp.1–26.

106. Levin, M., 1999. Left-Right Asymmetry in Animal Embryogenesis. Advances in Biochirality.

107. Lockley, M.G., McCrea, R.T., Buckley, L.G., Deock Lim, J., Matthews, N.A., Breithaupt, B.H., Houck, K.J., Gierliński, G.D., Surmik, D., Soo Kim, K. and Xing, L., 2016. Theropod courtship: large scale physical evidence of display arenas and avian-like scrape ceremony behaviour by Cretaceous dinosaurs. Scientific Reports, 6(1), p.18952.

108. Lyon Jr, M. W., 1908. Remarks on the Horns and on the Systematic Position of the American Antelope. Proceedings of the United States National Museum.

109. Maidment, S.C., 2023. Diversity through time and space in the Upper Jurassic Morrison Formation, Western USA. Journal of Vertebrate Paleontology, 43(5), p.e2326027.

110. Maden, M., Polvadore, T., Polanco, A., Barbazuk, W. B. and Stanley, E., 2023. Osteoderms in a mammal the spiny mouse Acomys and the independent evolution of dermal armor. iScience, 26(6).

111. Maidment, S. C., Norman, D. B., Barrett, P. M. and Upchurch, P., 2008. Systematics and phylogeny of Stegosauria (Dinosauria: Ornithischia). Journal of Systematic Palaeontology, 6(4), pp.367–407.

112. Maidment, S. C. R., Linton, D. H., Upchurch P. and Barrett P. M., 2012. Limb-bone scaling indicates diverse stance and gait in quadrupedal ornithischian dinosaurs. PLoS ONE 7(5), e36904.

113. Maidment, S. C. R., Brassey, C. and Barrett, P. M., 2015. The postcranial skeleton of an exceptionally complete individual of the plated dinosaur *Stegosaurus stenops* (Dinosauria: Thyreophora) from the Upper Jurassic Morrison Formation of Wyoming, USA. PloS one, 10(10), p.e0138352.

114. Maidment, S. C., Woodruff, D. C. and Horner, J. R., 2018. A new specimen of the ornithischian dinosaur *Hesperosaurus mjosi* from the Upper Jurassic Morrison Formation of Montana, USA, and implications for growth and size in Morrison stegosaurs. Journal of Vertebrate Paleontology, 38(1), p.e1406366.

115. Main, R. P., De Ricqlès, A., Horner, J. R. and Padian, K., 2005. The evolution and function of thyreophoran dinosaur scutes: implications for plate function in stegosaurs. Paleobiology, 31(2), pp.291–314.

116. Mallison, H., 2010. CAD assessment of the posture and range of motion of Kentrosaurus aethiopicus Hennig 1915. Swiss Journal of Geosciences, 103(2), pp.211–233.

117. Mallison, H., 2011. Defense capabilities of Kentrosaurus aethiopicus Hennig, 1915. Palaeontologia Electronica, 14(2), pp.1–25.

118. Mallison, H., 2014. Osteoderm distribution has low impact on the centre of mass of stegosaurs. Fossil Record, 17(1), pp.33–39.

119. Mallon, J. C., 2017. Recognizing sexual dimorphism in the fossil record: lessons from nonavian dinosaurs. Paleobiology, 43(3), pp.495–507.

120. Maryańska, T., Chapman, R.E., Weishampel, D.B., 2004. Pachycephalosauria, in: Weishampel, D.B., Dodson, P., Osmólska, H. (Eds.), The Dinosauria. University of California Press, Berkeley, pp. 464–477.

121. Mateus, O., Maidment, S. C. and Christiansen, N. A., 2009. A new long-necked ‘sauropod-mimic’ stegosaur and the evolution of the plated dinosaurs. Proceedings of the royal society B: biological sciences, 276(1663), pp.1815–1821.

122. Marsh, O. C., 1877. A new order of extinct Reptilia (Stegosauria) from the Jurassic of the Rocky Mountains. American Journal of Science, 3(84), pp.513–514.

123. Maynard Smith, J. and Price, G. R., 1973. The logic of animal conflict. Nature, 246(5427), pp.15–18.

124. McWhinney, L. A., Rothschild, B. M. and Carpenter, K., 2001. Posttraumatic chronic osteomyelitis in Stegosaurus dermal spikes. The Armored Dinosaurs, pp.141-156.

125. Meissner, W., Remisiewicz, M. and Pilacka, L., 2021. Sexual size dimorphism and sex determination in Blacksmith Lapwing Vanellus armatus (Burchell, 1822)(Charadriiformes: Charadriidae). The European Zoological Journal, 88(1), pp.279–288.

126. Mendelson, T. C. and Shaw, K. L., 2013. Further misconceptions about species recognition: a reply to Padian and Horner. Trends in Ecology & Evolution, 28(5), pp.252–253.

127. Merchant, M., Hale, A., Brueggen, J., Harbsmeier, C. and Adams, C., 2018. Crocodiles alter skin color in response to environmental color conditions. Scientific reports, 8(1), p.6174.

128. Miles, M. C. and Fuxjager, M. J., 2018. Synergistic selection regimens drive the evolution of display complexity in birds of paradise. Journal of Animal Ecology, 87(4), pp.1149–1159.

129. Mora, J. M. and López Umaña, L. I., 2020. A strong case of dilution in the Yellow-throated Toucan (*Ramphastos ambiguus*). Huitzil, 21(2).

130. Motani, R. 2021. Sex estimation from morphology in living animals and dinosaurs. Zoological Journal of the Linnean Society. 10.1093/zoolinnean/zlaa181

131. Mukherjee, S. and Heithaus, M. R., 2013. Dangerous prey and daring predators: a review. Biological reviews, 88(3), pp.550–563.

132. Murphy, K., Ruth, T. K., Hornocker, M. and Negri, S., 2009. Diet and prey selection of a perfect predator. Cougar: ecology and conservation. *University of Chicago Press*, Chicago, Illinois, USA, pp.118–137.

133. Myhrvold, N.P. and Currie, P.J., 1997. Supersonic sauropods? Tail dynamics in the diplodocids. Paleobiology, 23(4), pp.393–409.

134. Myhrvold, N.P., Baumgart, S.L., Vidal, D., Fish, F.E., Henderson, D.M., Saitta, E.T. and Sereno, P.C., 2024. Diving dinosaurs? Caveats on the use of bone compactness and pFDA for inferring lifestyle. Plos one, 19(3), p.e0298957.

135. Nefdt, R. J. and Thirgood, S. J., 1997. Lekking, resource defense, and harassment in two subspecies of lechwe antelope. Behavioral Ecology, 8(1), pp.1–9.

136. Norman, D. B., Witmer, L. M. and Weishampel, D. B. (2004). “Basal Thyreophora”. In Weishampel, D. B., Dodson, P., & Osmolska, H. (eds.) The Dinosauria. Berkeley: Berkeley University Press, 343–362.

137. Nuechterlein, G.L. and Storer, R.W., 1985. Aggressive behavior and interspecific killing by Flying Steamer-Ducks in Argentina. The Condor, 87(1), pp.87–91.

138. Nweeia, M. T., Nutarak, C., Eichmiller, F. C., Eidelman, N., Giuseppetti, A. A., Quinn, J., Mead, J. G., K’issuk, K., Hauschka, P. V., Tyler, E. M. and Potter, C. W., 2009. Considerations of anatomy, morphology, evolution, and function for narwhal dentition. Smithsonian at the Poles: Contributions to International Polar Year Science.

139. Ord, T.J. and Stuart-Fox, D., 2006. Ornament evolution in dragon lizards: multiple gains and widespread losses reveal a complex history of evolutionary change. Journal of Evolutionary Biology, 19(3), pp.797–808.

140. Owen-Smith, N., 1993. Comparative mortality rates of male and female kudus: the costs of sexual size dimorphism. Journal of Animal Ecology, pp.428–440.

141. Packer, C., 1983. Sexual dimorphism: the horns of African antelopes. Science, 221(4616), pp.1191–1193.

142. Padian, K. and Horner, J. R., 2011. The evolution of ‘bizarre structures’ in dinosaurs: biomechanics, sexual selection, social selection or species recognition?. Journal of Zoology, 283(1), pp.3–17.

143. Padian, K. and Horner, J. R., 2013. Misconceptions of sexual selection and species recognition: a response to Knell et al. and to Mendelson and Shaw. Trends in Ecology & Evolution, 28(5), pp.249–250.

144. Padian, K. and Horner, J. R., 2014. The species recognition hypothesis explains exaggerated structures in non-avialan dinosaurs better than sexual selection does. Comptes Rendus Palevol, 13(2), pp.97–107.

145. Paladino, F. V., O’Connor, M. P. and Spotila, J. R., 1990. Metabolism of leatherback turtles, gigantothermy, and thermoregulation of dinosaurs. Nature, 344(6269), pp.858–860.

146. Palmer, A. R., 2009. Animal asymmetry. Current Biology, 19(12), pp.R473–R477.

147. Panagiotopoulou, O., Spyridis, P., Abraha, H.M., Carrier, D.R. and Pataky, T.C., 2016. Architecture of the sperm whale forehead facilitates ramming combat. PeerJ, 4, p.e1895.

148. Pérez-Barbería, F. J., Gordon, I. J. and Pagel, M., 2002. The origins of sexual dimorphism in body size in ungulates. Evolution, 56(6), pp.1276–1285.

149. Persons IV, W. S., Funston, G. F., Currie, P. J. and Norell, M. A., 2015. A possible instance of sexual dimorphism in the tails of two oviraptorosaur dinosaurs. Scientific reports, 5, p.9472.

150. Peterson, J.E., Dischler, C. and Longrich, N.R., 2013. Distributions of cranial pathologies provide evidence for head-butting in dome-headed dinosaurs (Pachycephalosauridae). PloS one, 8(7), p.e68620.

151. Pianka, E.R. and Vitt, L.J., 2003. Lizards: windows to the evolution of diversity (Vol. 5). Univ of California Press.

152. Picard, K., Thomas, D. W., Festa-Bianchet, M. and Lanthier, C., 1994. Bovid horns: an important site for heat loss during winter?. Journal of Mammalogy, 75(3), pp.710–713.

153. Picard, K., Festa-Bianchet, M. and Thomas, D., 1996. The cost of horniness: heat loss may counter sexual selection for large horns in temperate bovids. Ecoscience, 3(3), pp.280–284.

154. Picard, K., Thomas, D. W., Festa-Bianchet, M., Belleville, F. and Laneville, A., 1999. Differences in the thermal conductance of tropical and temperate bovid horns. Ecoscience, 6(2), pp.148–158.

155. Pontzer, H., Allen, V. and Hutchinson, J. R., 2009. Biomechanics of running indicates endothermy in bipedal dinosaurs. PLoS One, 4(11), p.e7783.

156. Prado-Irwin, S.R., Revell, L.J. and Winchell, K.M., 2019. Variation in tail morphology across urban and forest populations of the crested anole (Anolis cristatellus). Biological Journal of the Linnean Society, 128(3), pp.632–644.

157. Prins, H.H.T., 1989. Condition changes and choice of social environment in African buffalo bulls. Behaviour, 108(3-4), pp.297–323.

158. Pruitt, W. O., 1966. The function of the brow-tine in caribou antlers. Arctic, 19(2), pp.111–113.

159. Prum, R. O. and Torres, R., 2003. Structural colouration of avian skin: convergent evolution of coherently scattering dermal collagen arrays. Journal of Experimental Biology, 206(14), pp.2409–2429.

160. Rand, A.L., 1954. On the spurs on birds’ wings. The Wilson Bulletin, pp.127-134.

161. Raven, T. J. and Maidment, S. C., 2017. A new phylogeny of Stegosauria (Dinosauria, Ornithischia). Palaeontology, 60(3), pp.401–408.

162. Reaney, L. T., Milner, R. N., Detto, T. and Backwell, P. R., 2008. The effects of claw regeneration on territory ownership and mating success in the fiddler crab Uca mjoebergi. Animal Behaviour, 75(4), pp.1473–1478.

163. Redelstorff, R. and Sander, P. M., 2009. Long and girdle bone histology of *Stegosaurus*: implications for growth and life history. Journal of Vertebrate Paleontology, 29(4), pp.1087–1099.

164. Richmond, D., 2023. Stratigraphy, sedimentology, and paleoclimatic proxies of the Upper Jurassic Morrison Formation of central Montana: Geology of the Intermountain West. Geology of the Intermountain West, 10, pp.223–276.

165. Rico-Guevara, A. and Hurme, K.J., 2019. Intrasexually selected weapons. Biological Reviews, 94(1), pp.60–101.

166. Rinehart, L. F., Lucas, S. G., Heckert, A. B., Spielmann, J. A. and Celeskey, M. D., 2009. The Paleobiology of Coelophysis bauri (Cope) from the Upper Triassic (Apachean) Whitaker quarry, New Mexico, with detailed analysis of a single quarry block: Bulletin 45 (Vol. 45). New Mexico Museum of Natural History and Science.

167. Richter, M. 2011. Fighting pair of dino skeletons goes for $2.75 million. NBC News. June 12, 2011. https://www.nbcnews.com/id/wbna43374471

168. Roberts, S. C., 1996. The evolution of hornedness in female ruminants. Behaviour, 133(5-6), pp.399–442.

169. Rothschild, B.M. and Berman, D.S., 1991. Fusion of caudal vertebrae in Late Jurassic sauropods. Journal of Vertebrate Paleontology, 11(1), pp.29–36.

170. Roy, A., Pittman, M., Saitta, E. T., Kaye, T. G. and Xu, X., 2020. Recent advances in amniote palaeocolour reconstruction and a framework for future research. Biological Reviews, 95(1), pp.22–50.

171. Rutland, C. S., Cigler, P. and Kubale, V., 2019. Reptilian skin and its special histological structures. Veterinary anatomy and physiology, pp.135-156.

172. Saitta, E. T., 2014. Paleobiology of North American Stegosaurs: Evidence for Sexual Dimorphism. Princeton University Senior Thesis.

173. Saitta, E. T. 2015. Evidence for Sexual Dimorphism in the Plated Dinosaur *Stegosaurus mjosi* (Ornithischia, Stegosauria) from the Morrison Formation (Upper Jurassic) of Western USA. PLoS ONE, 10:e0123503.

174. Saitta, E. T., Stockdale, M. T., Longrich, N. R., Bonhomme, V., Benton, M. J., Cuthill, I. C. and Makovicky, P. J., 2020. An effect size statistical framework for investigating sexual dimorphism in non-avian dinosaurs and other extinct taxa. Biological Journal of the Linnean Society, 131(2), pp.231–273.

175. Saleiro, A. and Mateus, O., 2017. Upper Jurassic bonebeds around Ten Sleep, Wyoming, USA: overview and stratigraphy. Abstract Book of the XV Encuentro de Jóvenes Investigadores en Paleontología/XV Encontro de Jovens Investigadores em Paleontologia, Lisboa, pp.357–361.

176. Schall, J.J., Bromwich, C.R., Werner, Y.L. and Midlege, J., 1989. Clubbed regenerated tails in Agama agama and their possible use in social interactions. Journal of Herpetology, 23(3), pp.303–305.

177. Schilling, M.F., Watkins, A.E. and Watkins, W., 2002. Is human height bimodal?. The American Statistician, 56(3), pp.223–229.

178. Schmude, D. E. and Weege, C. J., 1996. Stratigraphic relationship, sedimentology, and taphonomy of Meilyn, a dinosaur quarry in the basal Morrison Formation of Wyoming. The Continental Jurassic. Museum of Northern Arizona Bulletin, 60, pp.547–554.

179. Sereno, P.C., Myhrvold, N., Henderson, D.M., Fish, F.E., Vidal, D., Baumgart, S.L., Keillor, T.M., Formoso, K.K. and Conroy, L.L., 2022. Spinosaurus is not an aquatic dinosaur. Elife, 11, p.e80092.

180. Siber, H. J., & Möckli, U. (2009). The Stegosaurs of the Sauriermuseum Aathal. Aathal,! Switzerland: Sauriermuseum Aathal.

181. Simmons, R.E. and Scheepers, L., 1996. Winning by a neck: sexual selection in the evolution of giraffe. The American Naturalist, 148(5), pp.771–786.

182. Smith, D. K., 1998. A morphometric analysis of *Allosaurus*. Journal of Vertebrate Paleontology, 18(1), pp.126–142.

183. Snively, E. and Theodor, J.M., 2011. Common functional correlates of head-strike behavior in the pachycephalosaur *Stegoceras validum* (Ornithischia, Dinosauria) and combative artiodactyls. PLoS One, 6(6), p.e21422.

184. Sonoda, T. and Noda, Y., 2016. Transfer of museum collection from the hayashibara museum of natural sciences to the fukui prefectural dinosaur museum. Memoir of the Fukui Prefectural Dinosaur Museum, (15), pp.93–98.

185. Soto-Acuña, S., Vargas, A.O., Kaluza, J., Leppe, M.A., Botelho, J.F., Palma-Liberona, J., Simon-Gutstein, C., Fernández, R.A., Ortiz, H., Milla, V. and Aravena, B., 2021. Bizarre tail weaponry in a transitional ankylosaur from subantarctic Chile. Nature, 600(7888), pp.259–263.

186. Stankowich, T. and Caro, T., 2009. Evolution of weaponry in female bovids. Proceedings of the Royal Society B: Biological Sciences, 276(1677), pp.4329–4334.

187. Sternberg, C.M., 1950. Pachyrhinosaurus canadensis, representing a new family of the Ceratopsia, from southern Alberta. National Museum of Canada Bulletin, 118, pp.109–120.

188. Summers, T. C. and Ord, T. J., 2022. Female preference for super-sized male ornaments and its implications for the evolution of ornament allometry. Evolutionary Ecology, 36(4), pp.701–716.

189. Swaddle, J. P., 2003. Fluctuating asymmetry, animal behavior, and evolution. Advances in the Study of Behavior, 32, pp.169–206.

190. Tattersall, G. J., 2016. Infrared thermography: A non-invasive window into thermal physiology. Comparative Biochemistry and Physiology Part A: Molecular & Integrative Physiology, 202, pp.78–98.

191. Tattersall, G. J., Andrade, D. V. and Abe, A. S., 2009. Heat exchange from the toucan bill reveals a controllable vascular thermal radiator. science, 325(5939), pp.468–470.

192. Taylor, G.W., Santos, J.C., Perrault, B.J., Morando, M., Vásquez Almazán, C.R. and Sites Jr, J.W., 2017. Sexual dimorphism, phenotypic integration, and the evolution of head structure in casque-headed lizards. Ecology and Evolution, 7(21), pp.8989–8998.

193. Terai, S., Endo, H., Rerkamnuaychoke, W., Hondo, E., Agungpriyono, S., Kitamura, N., Kurohmaru, M., Kimura, J., Hayashi, Y., Nishida, T. and Yamada, J., 1998. An osteometrical study of the cranium and mandible of the lesser mouse deer (Chevrotain), Tragulus javanicus. Journal of veterinary medical science, 60(10), pp.1097–1105.

194. Tomkins, J. L., LeBas, N. R., Witton, M. P., Martill, D. M. and Humphries, S., 2010. Positive allometry and the prehistory of sexual selection. The American Naturalist, 176(2), pp.141–148.

195. Vickaryous, M.K. and Hall, B.K., 2006. Osteoderm morphology and development in the nine-banded armadillo, Dasypus novemcinctus (Mammalia, Xenarthra, Cingulata). Journal of Morphology, 267(11), pp.1273–1283.

196. Vickaryous, M.K. and Hall, B.K., 2008. Development of the dermal skeleton in *Alligator mississippiensis* (Archosauria, Crocodylia) with comments on the homology of osteoderms. Journal of morphology, 269(4), pp.398–422.

197. Vickaryous, M. K., Maryańska, T. and Weishampel, D. B. (2004). “Ankylosauria”. In Weishampel, D. B., Dodson, P., & Osmolska, H. (eds.) The Dinosauria. Berkeley: Berkeley University Press, 343–362.

198. Vidal, D., Ortega, F., Gascó, F., Serrano-Martínez, A. and Sanz, J.L., 2017. The internal anatomy of titanosaur osteoderms from the Upper Cretaceous of Spain is compatible with a role in oogenesis. Scientific reports, 7(1), p.42035.

199. Wiemann, J., Menéndez, I., Crawford, J.M., Fabbri, M., Gauthier, J.A., Hull, P.M., Norell, M.A. and Briggs, D.E., 2022. Fossil biomolecules reveal an avian metabolism in the ancestral dinosaur. Nature, 606(7914), pp.522–526.

200. Weishampel, D. B. and Chapman, R. E., 1990. Morphometric study of Plateosaurus from Trossingen (Baden-Württemberg, Federal Republic of Germany). Dinosaur Systematics: Approaches and Perspectives, pp.43–51.

201. Weissenböck, N.M., Weiss, C.M., Schwammer, H.M. and Kratochvil, H., 2010. Thermal windows on the body surface of African elephants (*Loxodonta africana*) studied by infrared thermography. Journal of Thermal Biology, 35(4), pp.182–188.

202. Whitlock, J. A., Trujillo, K. C. and Hanik, G. M., 2018. Assemblage-level structure in Morrison Formation dinosaurs, western interior, USA. Geology of the Intermountain West, 5, pp.9–22.

203. Woodruff, D. C., Trexler, D. and Maidment, S. C., 2019. Two new stegosaur specimens from the Upper Jurassic Morrison Formation of Montana, USA. Acta Palaeontologica Polonica, 64(3).

204. Wright, E., Grawunder, S., Ndayishimiye, E., Galbany, J., McFarlin, S. C., Stoinski, T. S. and Robbins, M. M., 2021. Chest beats as an honest signal of body size in male mountain gorillas (*Gorilla beringei beringei*). Scientific Reports, 11(1), p.6879.

