## Supplementary material for "The function and Evolution of Stegosaur Osteoderms and Hypothesized Sexual Dimorphism in *Hesperosaurus*": PCA Script.R

```
#Read data
Data<-read.csv("Data for PCA.csv", header = TRUE)

#####################################################################################################
#With juvenile plate
Data_pca<-prcomp(Data[,2:7], scale = TRUE)
Data_pca
summary(Data_pca)
plot(Data_pca$x[0:40,1],Data_pca$x[0:40,3],cex=1.5, cex.lab=1.25,xlab=expression("PC1"),ylab=expression("PC3"))
points(Data_pca$x[41,1], Data_pca$x[41,3], col="black", pch=2, cex=1.5)
abline(h=0)

Combo <- cbind(Data_pca$x[,1:3],Data)
wide <- Combo[which(Combo$S..mjosi.morph=="Wide"),]
tall <- Combo[which(Combo$S..mjosi.morph=="Tall"),]
juv <- Combo[which(Combo$S..mjosi.morph=="Stegosaurus"),]
points(wide$PC1, wide$PC3, col="black", pch=16, cex=1.5)
points(tall$PC1, tall$PC3, col="grey", pch=16, cex=1.5)
points(Data_pca$x[41,1], Data_pca$x[41,3], col="black", pch=2, cex=1.5)
points(juv$PC1, juv$PC3, col="white", pch=17, cex=1.5)

#####################################################################################################
#Without juvenile plate
NoJuv<-Data[0:40,]
NoJuv_pca<-prcomp(NoJuv[,2:7], scale = TRUE)
plot(NoJuv_pca$x[,1],NoJuv_pca$x[,3],cex=1.5, cex.lab=1.25,xlab=expression("PC1"),ylab=expression("PC3"))
summary(NoJuv_pca)
NoJuv_pca

Combo_NoJuv <- cbind(NoJuv_pca$x[,1:3],NoJuv)
wide_NoJuv <- Combo_NoJuv[which(Combo_NoJuv$S..mjosi.morph=="Wide"),]
tall_NoJuv <- Combo_NoJuv[which(Combo_NoJuv$S..mjosi.morph=="Tall"),]
points(wide_NoJuv$PC1, wide_NoJuv$PC3, col="black", pch=16, cex=1.5)
points(tall_NoJuv$PC1, tall_NoJuv$PC3, col="orange", pch=16, cex=1.5)
abline(h=0)

# Fit linear regression to data
REG<-cbind(NoJuv_pca$x[,1],NoJuv_pca$x[,3])
colnames(REG)<-c("PC1", "PC3")
row.names(REG)<-paste0("Specimen_", c(1:length(REG[,1])))
model.s1<-lm(REG[,2]~REG[,1])
model.s1
summary(model.s1)

x <- REG[,1]
y <- REG[,2]
plot(x, y, cex.lab=1.25, xlab="PC1", ylab="PC3")
predict(lm(y ~ x))
min<-min(REG[,1])
max<-max(REG[,1])
new <- data.frame(x = seq(min, max, 0.01))
predict(lm(y ~ x), new, se.fit = TRUE)
pred.w.plim <- predict(lm(y ~ x), new, interval = "prediction")
pred.w.clim <- predict(lm(y ~ x), new, interval = "confidence")
lines(new$x, pred.w.clim[ , 1], lty=1, col="red", lwd=3)
lines(new$x, pred.w.clim[ , 2], lty=2, col='blue', lwd=3)
lines(new$x, pred.w.clim[ , 3], lty=2, col='blue', lwd=3)
lines(new$x, pred.w.plim[ , 2], lty=3, col='green', lwd=3)
lines(new$x, pred.w.plim[ , 3], lty=3, col='green', lwd=3)

# Residual plot
residuals.1<-model.s1$residuals
plot(REG[,1], as.vector(residuals.1), pch=19,cex=1.5,xlab=expression("PC1"),ylab=expression("Residuals"), cex.lab=1.5)
abline(h=0)

# Estimate sex based on Saitta (2015)
steg.male <- subset(Combo_NoJuv, Combo_NoJuv$S..mjosi.morph=="Wide")
steg.female <- subset(Combo_NoJuv, Combo_NoJuv$S..mjosi.morph=="Tall")
plot(x=as.vector(steg.male[,1]), y=as.vector(steg.male[,3]), xlab="PC1", ylab="PC3", xlim=c(-6,3), ylim=c(-1.2,1.2),cex.lab=1.25, col="black")
points(x=as.vector(steg.female[,1]), y=as.vector(steg.female[,3]), col="orange")

# Wide morph Intervals
x <- as.numeric(as.vector(steg.male[,1]))
y <- as.numeric(as.vector(steg.male[,3]))
plot(x, y, xlab="PC1", ylab="PC3", xlim=c(-6,3), ylim=c(-1.2,1.2), col="black", cex=1.5, pch=16, cex.lab=1.5)
predict(lm(y ~ x))
min<-min(as.numeric(as.vector(steg.male[,1])))
max<-max(as.numeric(as.vector(steg.male[,1])))
new <- data.frame(x = seq(min, max, 0.01))
predict(lm(y ~ x), new, se.fit = TRUE)
pred.w.plim <- predict(lm(y ~ x), new, interval = "prediction")
pred.w.clim <- predict(lm(y ~ x), new, interval = "confidence")
lines(new$x, pred.w.clim[ , 1], lty=1, col="black", lwd=3)
lines(new$x, pred.w.clim[ , 2], lty=2, col='black', lwd=3)
lines(new$x, pred.w.clim[ , 3], lty=2, col='black', lwd=3)
lines(new$x, pred.w.plim[ , 2], lty=3, col='black', lwd=3)
lines(new$x, pred.w.plim[ , 3], lty=3, col='black', lwd=3)

# Tall morph Intervals
x <- as.numeric(as.vector(steg.female[,1]))
y <- as.numeric(as.vector(steg.female[,3]))
points(x=as.vector(steg.female[,1]), y=as.vector(steg.female[,3]), col="grey", cex=1.5, pch=16)
predict(lm(y ~ x))
min<-min(as.numeric(as.vector(steg.female[,1])))
max<-max(as.numeric(as.vector(steg.female[,1])))
new <- data.frame(x = seq(min, max, 0.01))
predict(lm(y ~ x), new, se.fit = TRUE)
pred.w.plim <- predict(lm(y ~ x), new, interval = "prediction")
pred.w.clim <- predict(lm(y ~ x), new, interval = "confidence")
lines(new$x, pred.w.clim[ , 1], lty=1, col="grey", lwd=3)
lines(new$x, pred.w.clim[ , 2], lty=2, col='grey', lwd=3)
lines(new$x, pred.w.clim[ , 3], lty=2, col='grey', lwd=3)
lines(new$x, pred.w.plim[ , 2], lty=3, col='grey', lwd=3)
lines(new$x, pred.w.plim[ , 3], lty=3, col='grey', lwd=3)

#####################################################################################################
#Without juvenile plate and transforming surface area and angle according to Motani (2021)
rm(list = ls())
Data<-read.csv("Data for PCA.csv", header = TRUE)

#Without juvenile plate
NoJuv<-Data[0:40,]
NoJuv_pca<-prcomp(NoJuv[,c(3:5,7,9:10)], scale = TRUE)
plot(NoJuv_pca$x[,1],NoJuv_pca$x[,2],cex=1.5, cex.lab=1.25,xlab=expression("PC1"),ylab=expression("PC2"))
summary(NoJuv_pca)
NoJuv_pca

Combo_NoJuv <- cbind(NoJuv_pca$x[,1:2],NoJuv)
wide_NoJuv <- Combo_NoJuv[which(Combo_NoJuv$S..mjosi.morph=="Wide"),]
tall_NoJuv <- Combo_NoJuv[which(Combo_NoJuv$S..mjosi.morph=="Tall"),]
points(wide_NoJuv$PC1, wide_NoJuv$PC2, col="black", pch=16, cex=1.5)
points(tall_NoJuv$PC1, tall_NoJuv$PC2, col="orange", pch=16, cex=1.5)
abline(h=0)

# Fit linear regression to data
REG<-cbind(NoJuv_pca$x[,1],NoJuv_pca$x[,2])
colnames(REG)<-c("PC1", "PC2")
row.names(REG)<-paste0("Specimen_", c(1:length(REG[,1])))
model.s1<-lm(REG[,2]~REG[,1])
model.s1
summary(model.s1)

x <- REG[,1]
y <- REG[,2]
plot(x, y, cex.lab=1.25, xlab="PC1", ylab="PC2")
predict(lm(y ~ x))
min<-min(REG[,1])
max<-max(REG[,1])
new <- data.frame(x = seq(min, max, 0.01))
predict(lm(y ~ x), new, se.fit = TRUE)
pred.w.plim <- predict(lm(y ~ x), new, interval = "prediction")
pred.w.clim <- predict(lm(y ~ x), new, interval = "confidence")
lines(new$x, pred.w.clim[ , 1], lty=1, col="red", lwd=3)
lines(new$x, pred.w.clim[ , 2], lty=2, col='blue', lwd=3)
lines(new$x, pred.w.clim[ , 3], lty=2, col='blue', lwd=3)
lines(new$x, pred.w.plim[ , 2], lty=3, col='green', lwd=3)
lines(new$x, pred.w.plim[ , 3], lty=3, col='green', lwd=3)

# Residual plot
residuals.1<-model.s1$residuals
plot(REG[,1], as.vector(residuals.1), pch=19,cex=1.5,xlab=expression("PC1"),ylab=expression("Residuals"), cex.lab=1.5)
abline(h=0)

# Estimate sex based on Saitta (2015)
steg.male <- subset(Combo_NoJuv, Combo_NoJuv$S..mjosi.morph=="Wide")
steg.female <- subset(Combo_NoJuv, Combo_NoJuv$S..mjosi.morph=="Tall")
plot(x=as.vector(steg.male[,1]), y=as.vector(steg.male[,2]), xlab="PC1", ylab="PC2", xlim=c(-6,3), ylim=c(-2,2),cex.lab=1.25, col="black")
points(x=as.vector(steg.female[,1]), y=as.vector(steg.female[,2]), col="orange")

# Wide morph Intervals
x <- as.numeric(as.vector(steg.male[,1]))
y <- as.numeric(as.vector(steg.male[,2]))
plot(x, y, xlab="PC1", ylab="PC2", xlim=c(-6,3), ylim=c(-2,2), col="black", cex=1.5, pch=16, cex.lab=1.5)
predict(lm(y ~ x))
min<-min(as.numeric(as.vector(steg.male[,1])))
max<-max(as.numeric(as.vector(steg.male[,1])))
new <- data.frame(x = seq(min, max, 0.01))
predict(lm(y ~ x), new, se.fit = TRUE)
pred.w.plim <- predict(lm(y ~ x), new, interval = "prediction")
pred.w.clim <- predict(lm(y ~ x), new, interval = "confidence")
lines(new$x, pred.w.clim[ , 1], lty=1, col="black", lwd=3)
lines(new$x, pred.w.clim[ , 2], lty=2, col='black', lwd=3)
lines(new$x, pred.w.clim[ , 3], lty=2, col='black', lwd=3)
lines(new$x, pred.w.plim[ , 2], lty=3, col='black', lwd=3)
lines(new$x, pred.w.plim[ , 3], lty=3, col='black', lwd=3)

# Tall morph Intervals
x <- as.numeric(as.vector(steg.female[,1]))
y <- as.numeric(as.vector(steg.female[,2]))
points(x=as.vector(steg.female[,1]), y=as.vector(steg.female[,2]), col="grey", cex=1.5, pch=16)
predict(lm(y ~ x))
min<-min(as.numeric(as.vector(steg.female[,1])))
max<-max(as.numeric(as.vector(steg.female[,1])))
new <- data.frame(x = seq(min, max, 0.01))
predict(lm(y ~ x), new, se.fit = TRUE)
pred.w.plim <- predict(lm(y ~ x), new, interval = "prediction")
pred.w.clim <- predict(lm(y ~ x), new, interval = "confidence")
lines(new$x, pred.w.clim[ , 1], lty=1, col="grey", lwd=3)
lines(new$x, pred.w.clim[ , 2], lty=2, col='grey', lwd=3)
lines(new$x, pred.w.clim[ , 3], lty=2, col='grey', lwd=3)
lines(new$x, pred.w.plim[ , 2], lty=3, col='grey', lwd=3)
lines(new$x, pred.w.plim[ , 3], lty=3, col='grey', lwd=3)
```
