## Supplementary material for "The function and Evolution of Stegosaur Osteoderms and Hypothesized Sexual Dimorphism in *Hesperosaurus*": Figure Code.R

```
############################################################################################ Set the Working Directory ##
#########################################################################################################################

setwd("C:/Users/ms9112/OneDrive - University of Bristol/Desktop/Stegosaur stuff")

#########################################################################################################################
##################################################################################################### Import libraries ##
#########################################################################################################################

library(Momocs)
library(MASS)

##########################################################################################################################
############################################################################################## import the total dataset ##
##########################################################################################################################

total_dataset = read.csv("coordinates2.csv", header=TRUE)

firstrows_specimen = which(duplicated(total_dataset$name)==FALSE)

total_dataset_fac = total_dataset[firstrows_specimen, c(2:4)]

td_ldk = total_dataset[firstrows_specimen, c(8:9)]

total_dataset_ldk = list()

for(g in 1:dim(td_ldk)[1]){
  
  tdldk_rg = as.vector(td_ldk[g,])
  names(tdldk_rg) = NULL
  total_dataset_ldk[[g]] = as.numeric(tdldk_rg)
  
}

total_dataset_coo = list()

for(h in 1:length(firstrows_specimen)){
  
  if(h < length(firstrows_specimen)) specimen_subset_rows = (firstrows_specimen[h] : (firstrows_specimen[(h+1)])-1)
  if(h == length(firstrows_specimen)) specimen_subset_rows = firstrows_specimen[h]:dim(total_dataset)[1]
  specimen_rows = total_dataset[specimen_subset_rows,c(6,7)]
  specimen_rows = as.matrix(specimen_rows)
  row.names(specimen_rows) = NULL
  colnames(specimen_rows) = NULL
  total_dataset_coo[[h]] = specimen_rows
  
}

names(total_dataset_coo) = total_dataset$name[firstrows_specimen]


###########################################################################################################################
######################################################################################### Create the OUT format variable ##
###########################################################################################################################

#steg <- Out(total@coo, fac=total@fac, ldk=total@ldk)


steg <- Out(total_dataset_coo, fac=total_dataset_fac, ldk=total_dataset_ldk)

###########################################################################################################################
################################################################################### smooth moving average of coordinates ##
###########################################################################################################################

steg <- coo_smooth(steg, 500)

############################################################################################################################
############################################################################### resample outlines to 360 coordinates each ##
############################################################################################################################

steg <- coo_sample(steg, 360)

############################################################################################################################
################################################################################################## preview plate outlines ##
############################################################################################################################


for(h in 1:length(steg[[1]])){
  
  name = strsplit(strsplit(names(steg[[1]])[h], split="/Users/evansaitta/Desktop/Thesis/Momocs/ALL PLATES BLACK/")[[1]][2], split=";")[[1]][1]
  jpeg(filename=paste0("plate_no_", h, ".jpg"), height=20, width=20, res=300, units="cm")
  plot(steg[[1]][[h]], type="l",lty=1, lwd=2, col="RED", asp=1, ylab="", xlab="", yaxt="n", xaxt="n", main=names(steg[[1]])[h])
  polygon(steg[[1]][[h]], col="BLACK")
  dev.off()
  
  print(paste0("successfully printed plate preview no.", h))
  
}


############################################################################################################################
####################################################################################### procrustese alignment of outlines ##
############################################################################################################################

# VB: if you pass fgProcrustes with coo=T it means that coordinates themselves (not the landmarks)
# will be used for the alignment. And also coo_slide(ldk=1) is mandatory so that the first 
# landmarks becomes a ± homologous point from where to start EFTs
# I would go for a coo_bookstein instead which possibly leads to a more sensible 
# alignment. Pretty sure we had the discussion before but cannot remember well.

# Also (later), if you do not use efourier with norm=FALSE then the numerical aligment
# will be used (the so-called 1st ellipsis alignment) and in any case it would care of 
# fgProcrustes, coo_bookstein or any other alignment on the outlines.

# The two problems combined led to symmetry on the morphospaces. 
# Where symmetry happens is shape-dependant but there you have on PC2
# (on top of PC2 is more or less the bottom of PC2 but flipped)
# this is bad, yet dozens of papers are published this way every year...

# Finally, I think the morphospace is just produced seamlessly by calling plot_PCA 
# and not plot(your_PCA object) which calls plot.default, not plot_PCA.
steg_al <- fgProcrustes(steg, coo=T)
steg_al <- coo_slide(steg_al, ldk=1)  

# I would go for:
steg_eft_vb <- steg %>% coo_bookstein() %>% coo_slide(ldk=1) %>% efourier(10, norm=FALSE)

# rather than:
#steg_eft <- steg_al %>% efourier(10)

#steg_eft %>% PCA %>% plot_PCA(~position)    # symmetry on PC2: bad
steg_eft_vb %>% PCA %>% plot_PCA(~position) # no symmetry on PC2: good
steg_eft_vb %>% PCA %>% plot_PCA(~morph) 

steg_eft_vb %>% PCA() %>% MANOVA_PW(~position) # and all differences are captured
steg_eft_vb %>% PCA() %>% MANOVA_PW(~morph) 
# this is not crystal clear on the PCA (which is not meant to capture inter group differences anyway)
# but see this lovely LDA plane ("yours" was nice too ;-)
steg_eft_vb %>% LDA(~position) %>% plot_LDA(morphospace=T)
steg_eft_vb %>% LDA(~morph) %>% plot_LDA(morphospace=T)

# In short, I'd go for coo_bookstein %>% coo_slide %>% efourier(norm=F)
# as it is the right way to do it I think and still, maintain your results. Which is fine, isn't ? :-)

#### end VB
```
