## Supplementary material for "The function and Evolution of Stegosaur Osteoderms and Hypothesized Sexual Dimorphism in *Hesperosaurus*": Angle Plot.R

```
#Read data
data<-read.csv("Angle Data.csv")
#0 is Stegosaurus
#1 is H. mjosi wide morph
#2 is H. mjosi tall morph

#Articulated Stegosaurus sp. data
sumstego<-data[data$H..mjosi.morph=="0", ]
hist(sumstego$Angle.between.the.base.of.the.plate.and.the.apex..degrees., col=NULL, cex.axis=1.5, cex.lab=0.5)
sortsumstego<-sort(sumstego$Angle.between.the.base.of.the.plate.and.the.apex..degrees., decreasing=FALSE)
plot(sortsumstego, cex.axis=1.5, pch=16, cex.lab=0.5)

#Hesperosaurus data both morphs
sumhesp<-data[data$H..mjosi.morph!="0", ]
hist(sumhesp$Angle.between.the.base.of.the.plate.and.the.apex..degrees., col=NULL, cex.axis=1.5, cex.lab=0.5)
sortsumhesp<-sort(sumhesp$Angle.between.the.base.of.the.plate.and.the.apex..degrees.,decreasing=FALSE)
plot(sortsumhesp, cex.axis=1.5, pch=16, cex.lab=0.5)

#Hesperosaurus tall morph
sumtall<-data[data$H..mjosi.morph=="2", ]
hist(sumtall$Angle.between.the.base.of.the.plate.and.the.apex..degrees., col=NULL, cex.axis=1.5, cex.lab=0.5)
sortsumtall<-sort(sumtall$Angle.between.the.base.of.the.plate.and.the.apex..degrees.,decreasing=FALSE)
plot(sortsumtall, cex.axis=1.5, pch=16, cex.lab=0.5)
  
#Hesperosaurus wide morph
sumwide<-data[data$H..mjosi.morph=="1", ]
hist(sumwide$Angle.between.the.base.of.the.plate.and.the.apex..degrees., col=NULL, cex.axis=1.5, cex.lab=0.5)
sortsumwide<-sort(sumwide$Angle.between.the.base.of.the.plate.and.the.apex..degrees.,decreasing=FALSE)
plot(sortsumwide, cex.axis=1.5, pch=16, cex.lab=0.5)
```
