## Supplementary material for "The function and Evolution of Stegosaur Osteoderms and Hypothesized Sexual Dimorphism in *Hesperosaurus*": Spike Code.R

```
data<-read.csv("Spikes Final.csv", header=TRUE)

###############################
#ALL SPIKES

#anterior spike length density
ant<-data[!is.na(as.numeric(data$Length.from.tip.to.center.of.base..cm.)), ]
ant<-ant[ant$Anterior.or.Posterior=="anterior",]
ant<-ant$Length.from.tip.to.center.of.base..cm.
ant<-as.numeric(ant)
dant<-density(ant)
plot(dant, lty=2)
dant

#density function spike length
data$Length.from.tip.to.center.of.base..cm.
L<-data[!is.na(as.numeric(data$Length.from.tip.to.center.of.base..cm.)), ]
L<-L$Length.from.tip.to.center.of.base..cm.
L<-as.numeric(L)
dL<-density(L)
lines(dL)
dL

#posterior spike length density
post<-data[!is.na(as.numeric(data$Length.from.tip.to.center.of.base..cm.)), ]
post<-post[post$Anterior.or.Posterior=="posterior",]
post<-post$Length.from.tip.to.center.of.base..cm.
post<-as.numeric(post)
dpost<-density(post)
lines(dpost, lty=3)
dpost

#histogram spike length
hist(L)

#length vs. width at midshaft
LvW<-data[!is.na(as.numeric(data$Length.from.tip.to.center.of.base..cm.)), ]
LvW<-LvW[!is.na(as.numeric(LvW$Width.at.midshaft..anteroposterior..cm.)), ]
plot(LvW$Length.from.tip.to.center.of.base..cm., LvW$Width.at.midshaft..anteroposterior..cm.)
LvWhesp<-LvW[LvW$Genus=="Hesperosaurus",]
points(LvWhesp$Length.from.tip.to.center.of.base..cm., LvWhesp$Width.at.midshaft..anteroposterior..cm., pch=16)

#length vs. base length
LvL<-data[!is.na(as.numeric(data$Length.from.tip.to.center.of.base..cm.)), ]
LvL<-LvL[!is.na(as.numeric(LvL$Length.of.base..anteroposterior..cm.)), ]
plot(LvL$Length.from.tip.to.center.of.base..cm., LvL$Length.of.base..anteroposterior..cm.)
LvLhesp<-LvL[LvL$Genus=="Hesperosaurus",]
points(LvLhesp$Length.from.tip.to.center.of.base..cm., LvLhesp$Length.of.base..anteroposterior..cm., pch=16)

###############################
#Stegosaurus SPIKES

#anterior spike length density
antS<-data[!is.na(as.numeric(data$Length.from.tip.to.center.of.base..cm.)), ]
antS<-antS[antS$Anterior.or.Posterior=="anterior",]
antS<-antS[antS$Genus=="Stegosaurus",]
antS<-antS$Length.from.tip.to.center.of.base..cm.
antS<-as.numeric(antS)
dantS<-density(antS)
plot(dantS, lty=2, cex=1.5, cex.lab=1.25)
dantS

#add median line
abline(v=median(antS), lty=2, col="grey", lwd=2)

#density function spike length
data$Length.from.tip.to.center.of.base..cm.
Ls<-data[!is.na(as.numeric(data$Length.from.tip.to.center.of.base..cm.)), ]
Ls<-Ls[Ls$Genus=="Stegosaurus",]
Ls<-Ls$Length.from.tip.to.center.of.base..cm.
Ls<-as.numeric(Ls)
dLs<-density(Ls)
lines(dLs)
dLs

#add median line
abline(v=median(Ls), col="grey", lwd=2)

#posterior spike length density
postS<-data[!is.na(as.numeric(data$Length.from.tip.to.center.of.base..cm.)), ]
postS<-postS[postS$Anterior.or.Posterior=="posterior",]
postS<-postS[postS$Genus=="Stegosaurus",]
postS<-postS$Length.from.tip.to.center.of.base..cm.
postS<-as.numeric(postS)
dpostS<-density(postS)
lines(dpostS, lty=3)
dpostS

#add median line
abline(v=median(postS), lty=3, col="grey", lwd=2)

###############################
#Hesperosaurus SPIKES

#posterior spike length density
postH<-data[!is.na(as.numeric(data$Length.from.tip.to.center.of.base..cm.)), ]
postH<-postH[postH$Anterior.or.Posterior=="posterior",]
postH<-postH[postH$Genus=="Hesperosaurus",]
postH<-postH$Length.from.tip.to.center.of.base..cm.
postH<-as.numeric(postH)
dpostH<-density(postH)
plot(dpostH, lty=3, xlim=c(14,72))
dpostH

#anterior spike length density
antH<-data[!is.na(as.numeric(data$Length.from.tip.to.center.of.base..cm.)), ]
antH<-antH[antH$Anterior.or.Posterior=="anterior",]
antH<-antH[antH$Genus=="Hesperosaurus",]
antH<-antH$Length.from.tip.to.center.of.base..cm.
antH<-as.numeric(antH)
dantH<-density(antH)
lines(dantH, lty=2)
dantH

#density function spike length
data$Length.from.tip.to.center.of.base..cm.
Lh<-data[!is.na(as.numeric(data$Length.from.tip.to.center.of.base..cm.)), ]
Lh<-Lh[Lh$Genus=="Hesperosaurus",]
Lh<-Lh$Length.from.tip.to.center.of.base..cm.
Lh<-as.numeric(Lh)
dLh<-density(Lh)
lines(dLh)
dLh
```
