## Supplementary figures and images for "The function and Evolution of Stegosaur Osteoderms and Hypothesized Sexual Dimorphism in *Hesperosaurus*"

### LDA morph.png

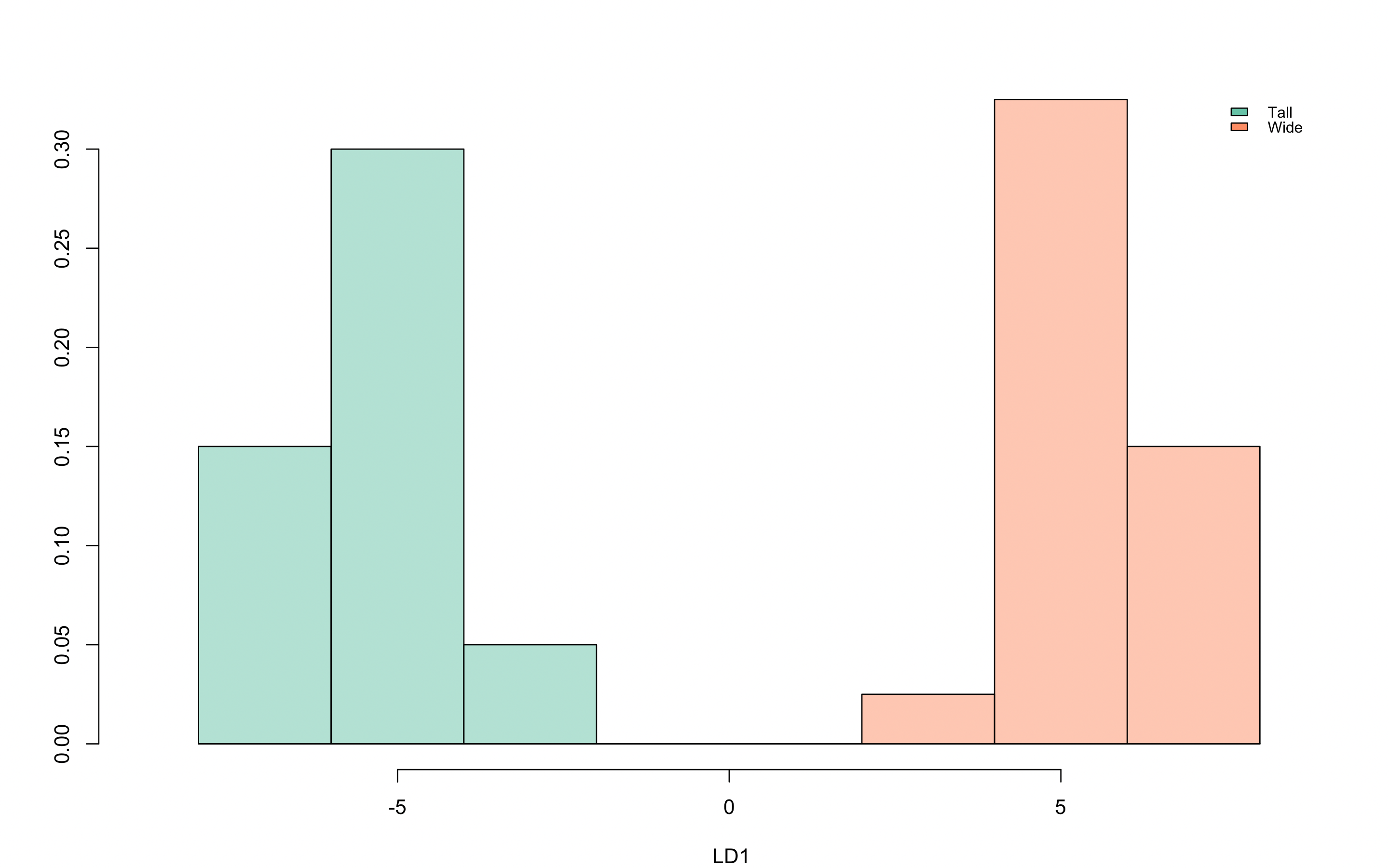

### LDA position.png

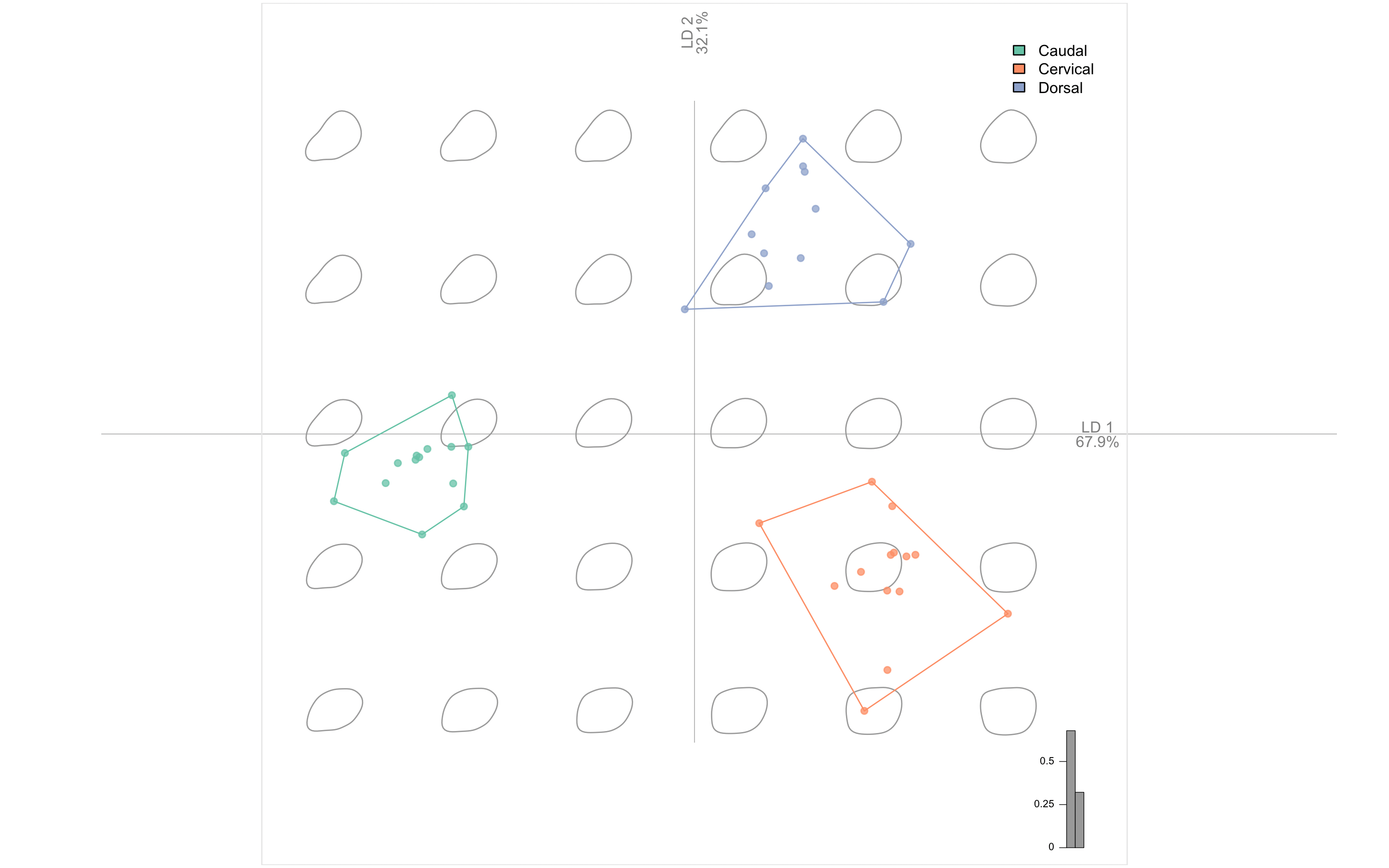

### MANOVA morph.png

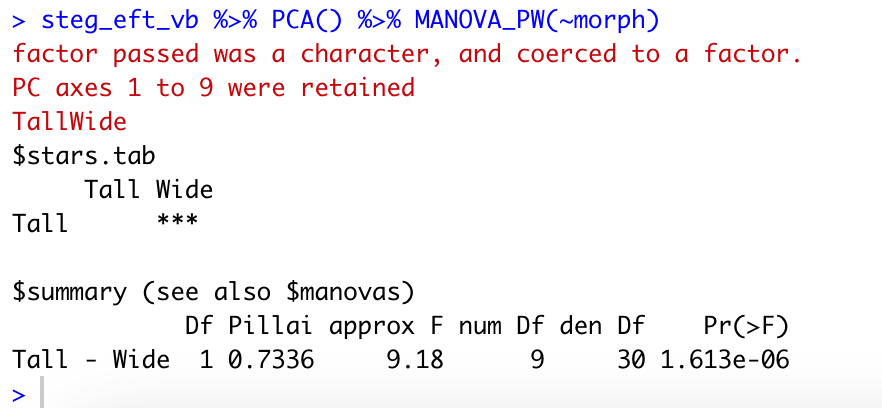

### MANOVA position.png

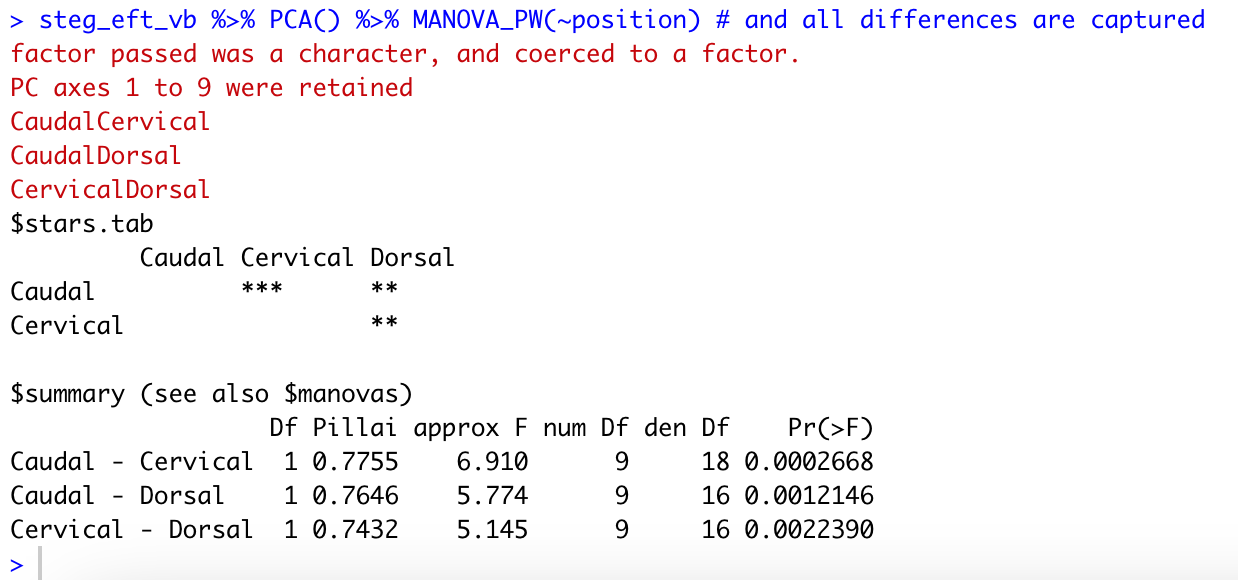

### PCA morph.png

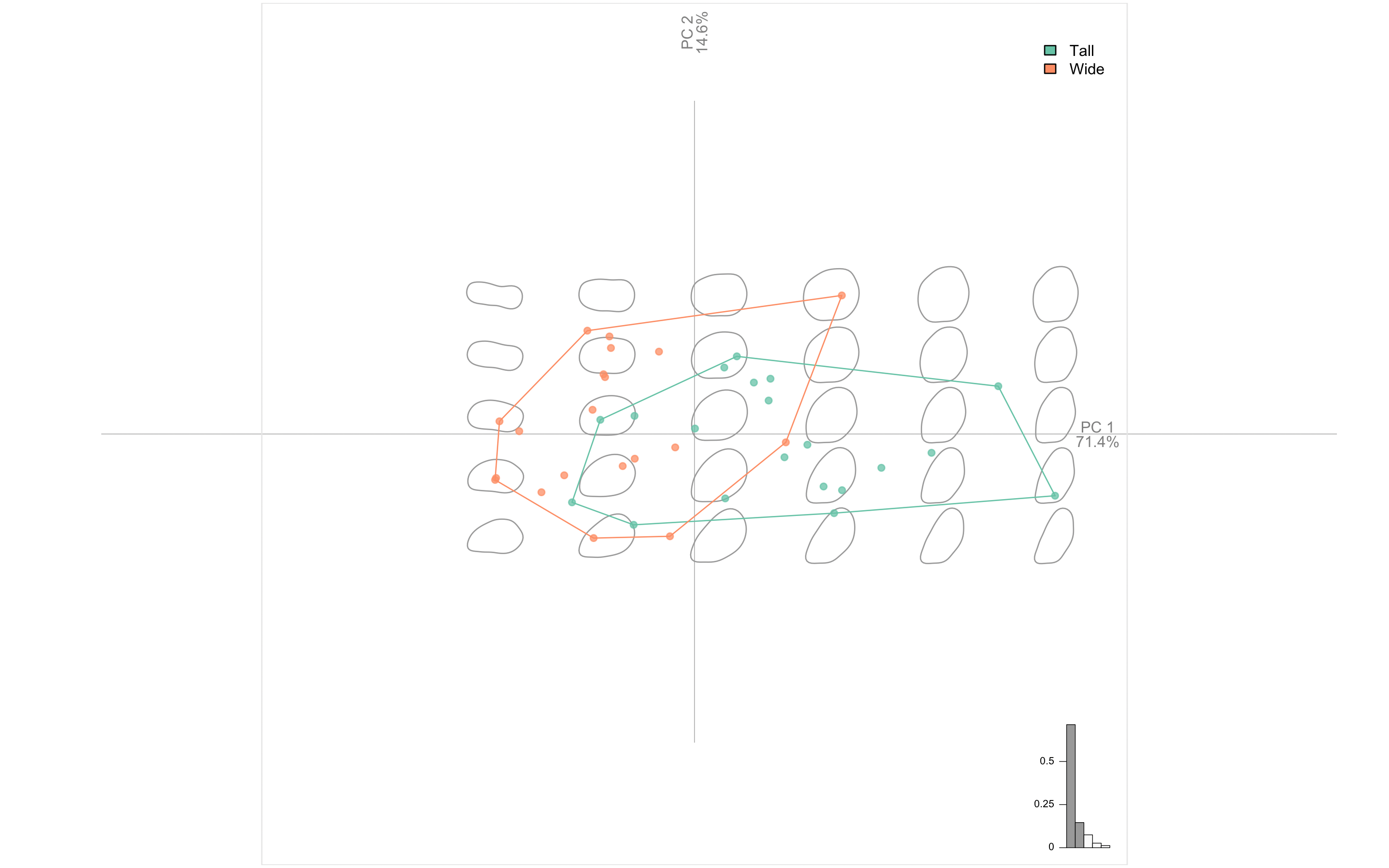

### PCA position.png

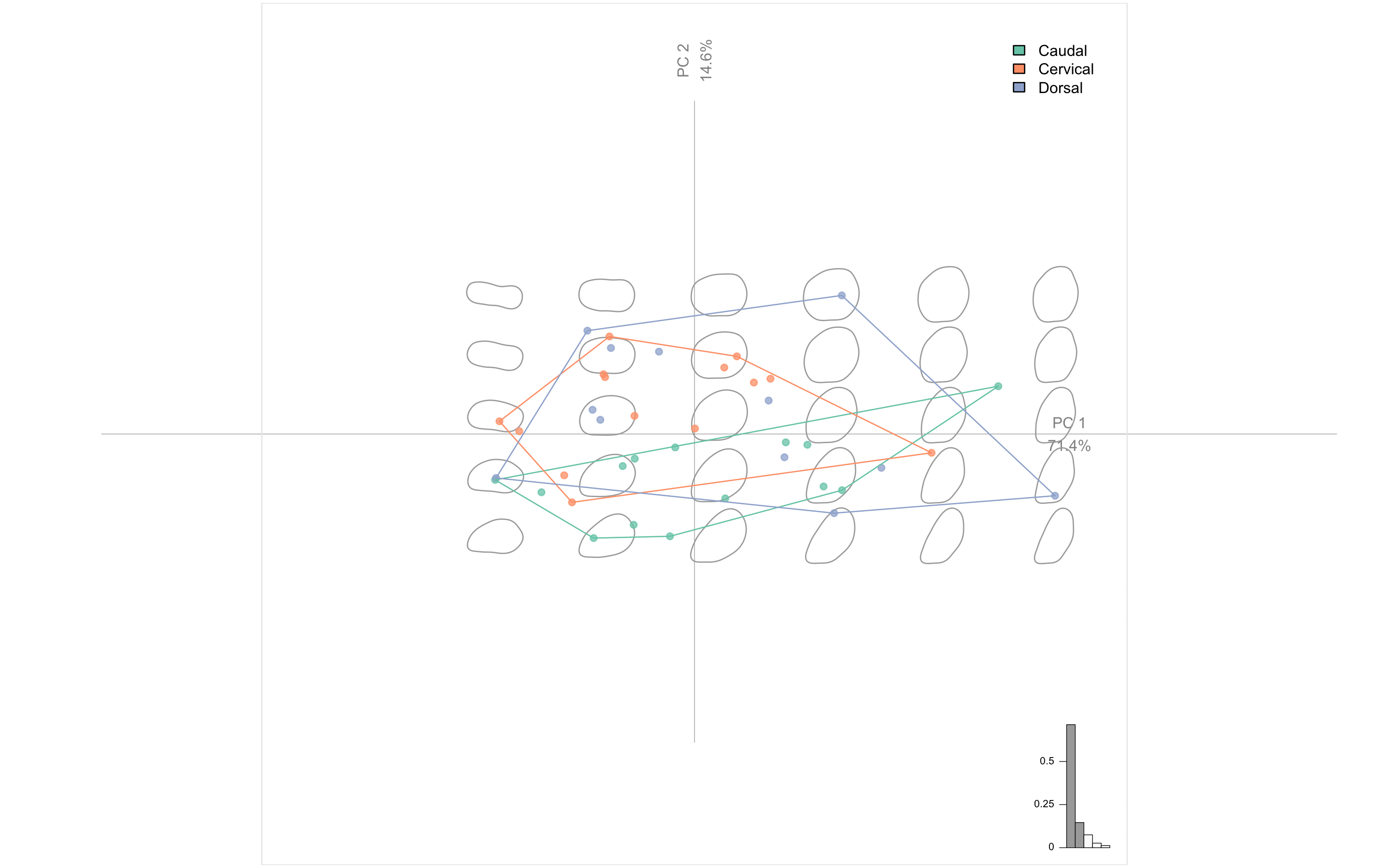

### PCA_12_morph.jpg

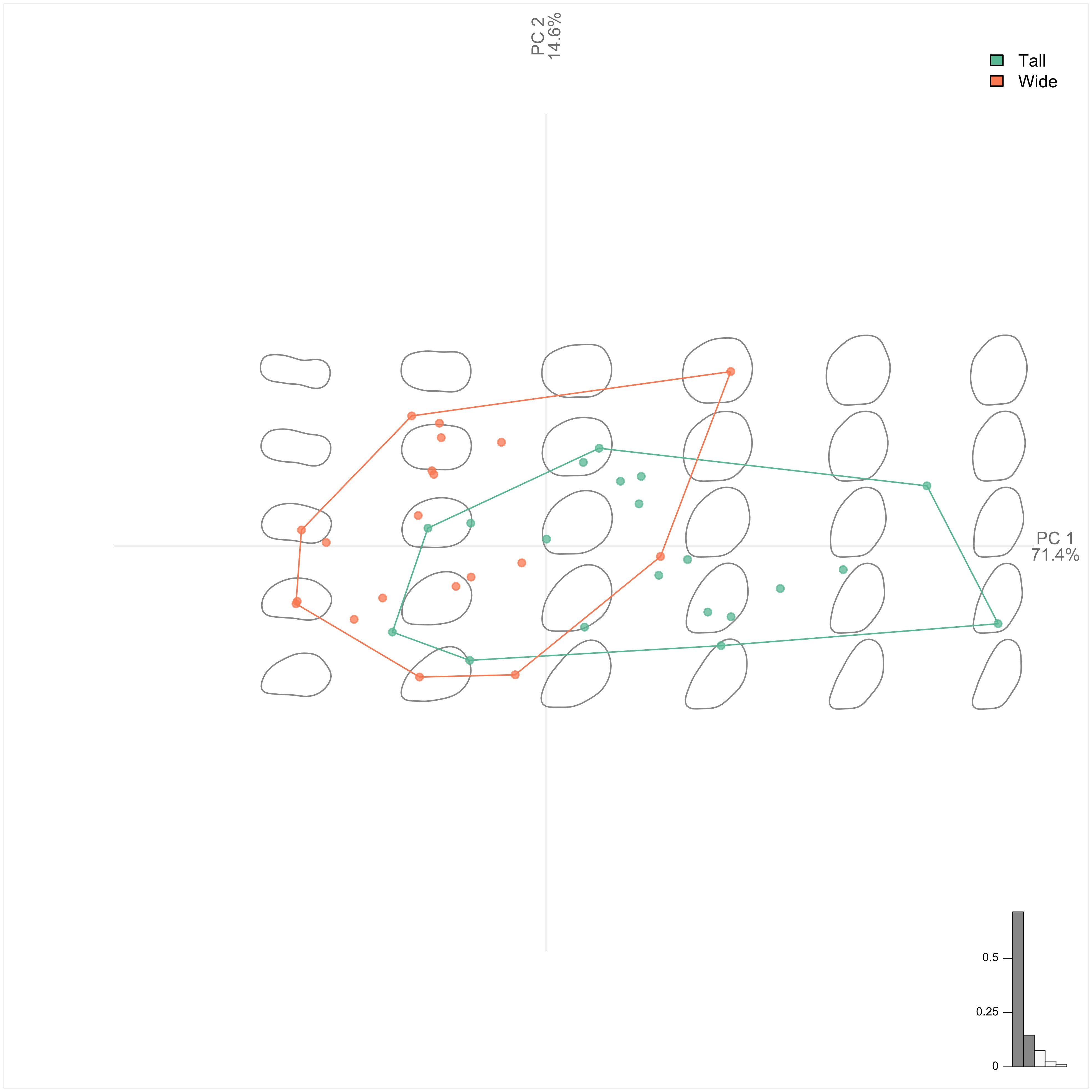

### PCA_12_position.jpg

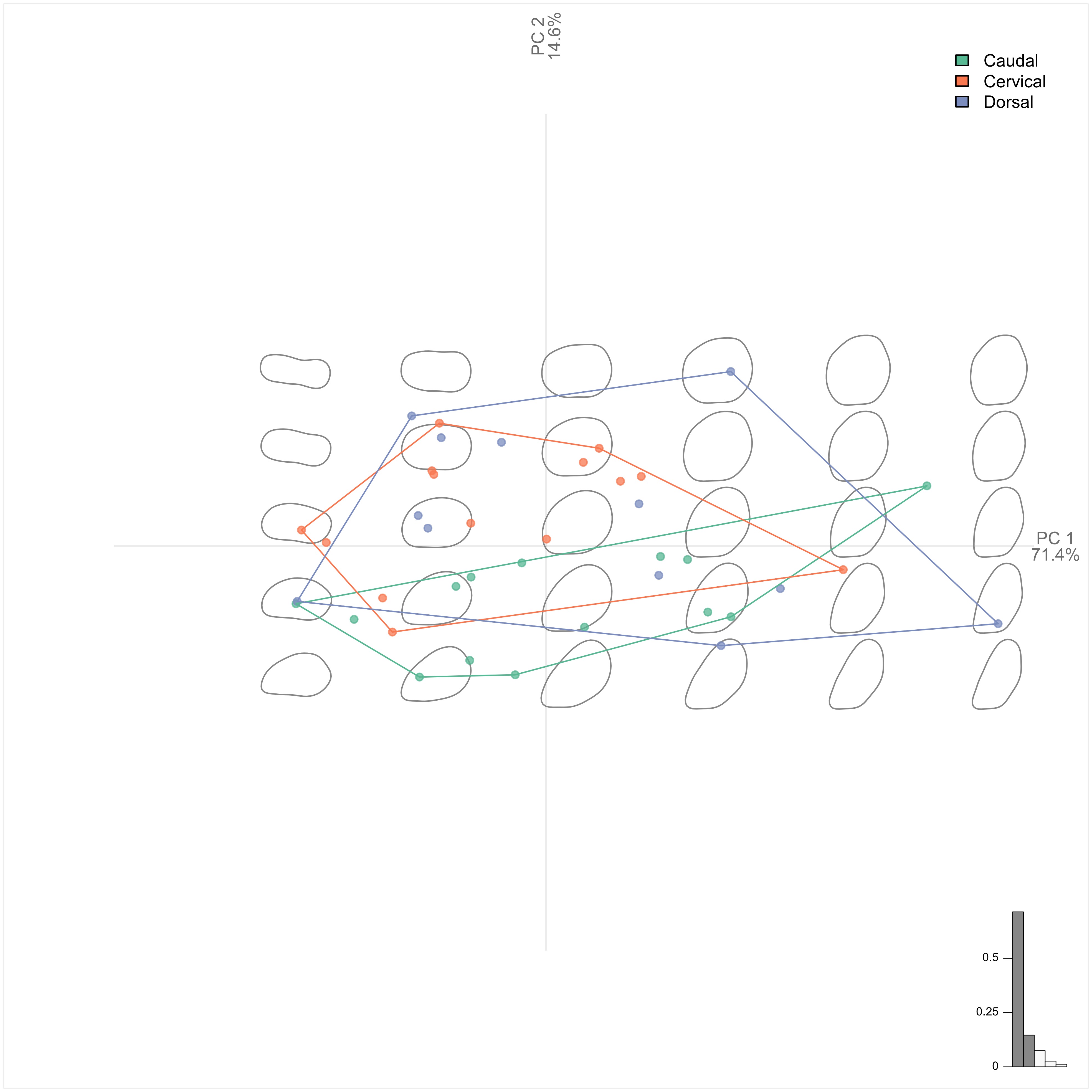

### PCA_13_morph.jpg

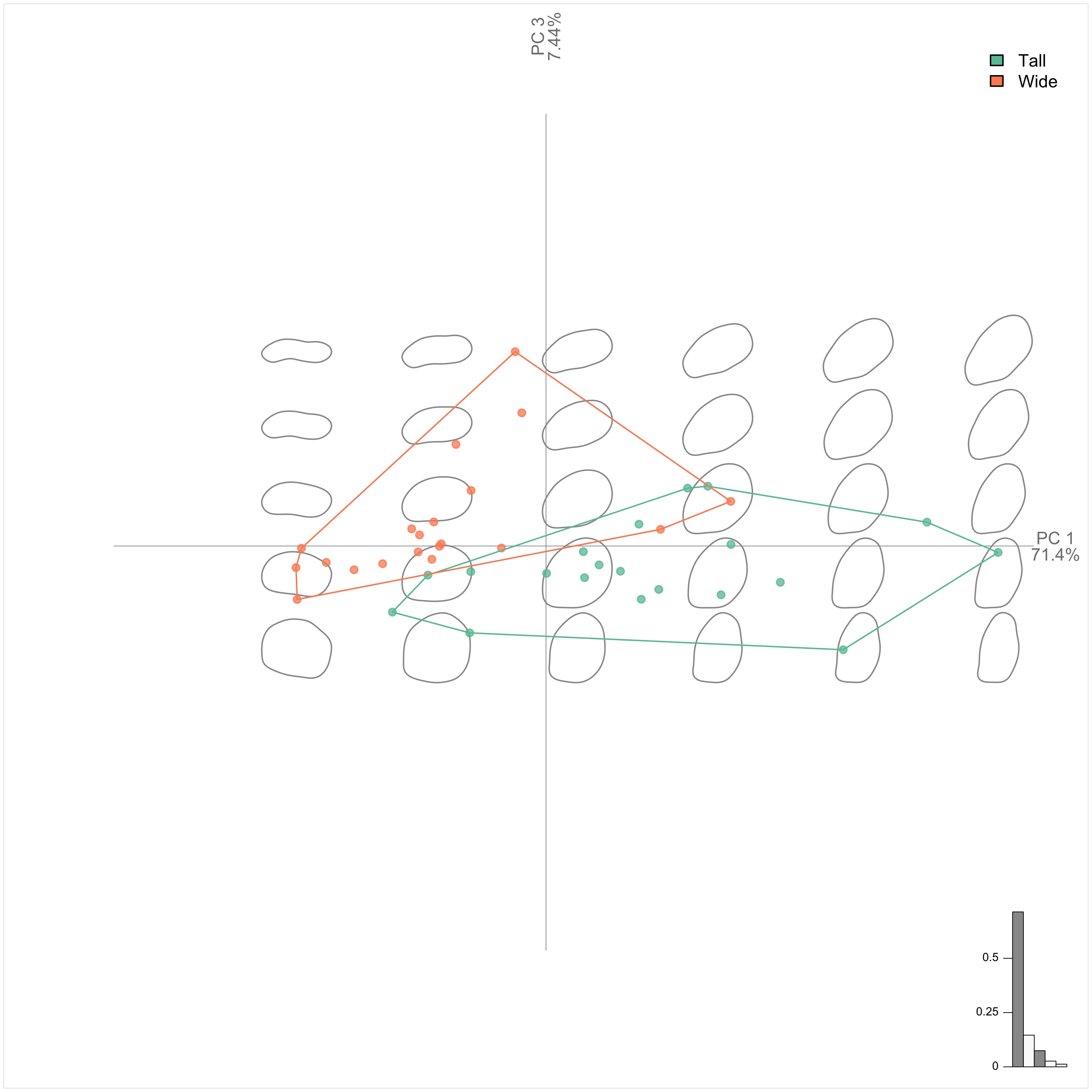

### PCA_13_position.jpg

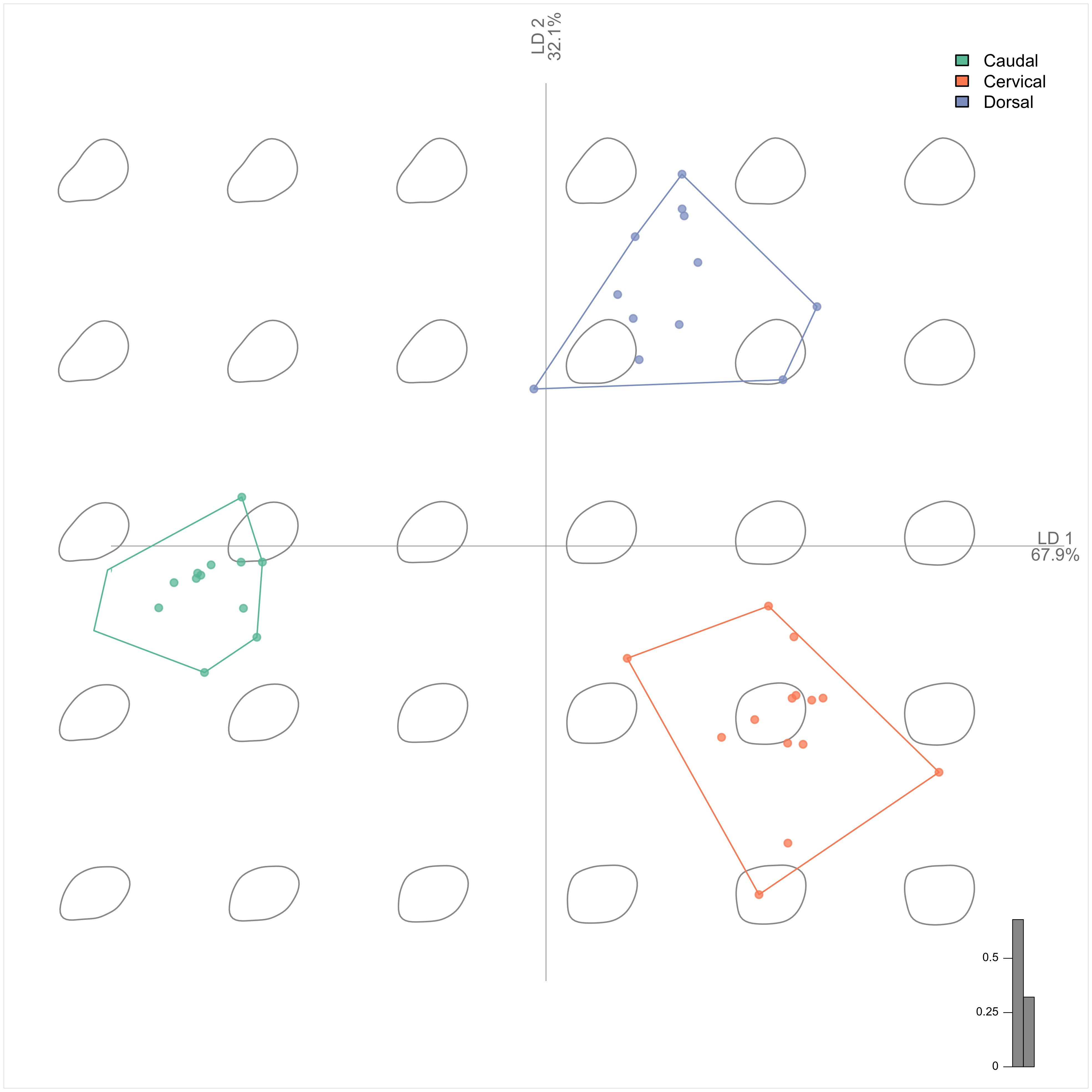

### plate_no_1.jpg

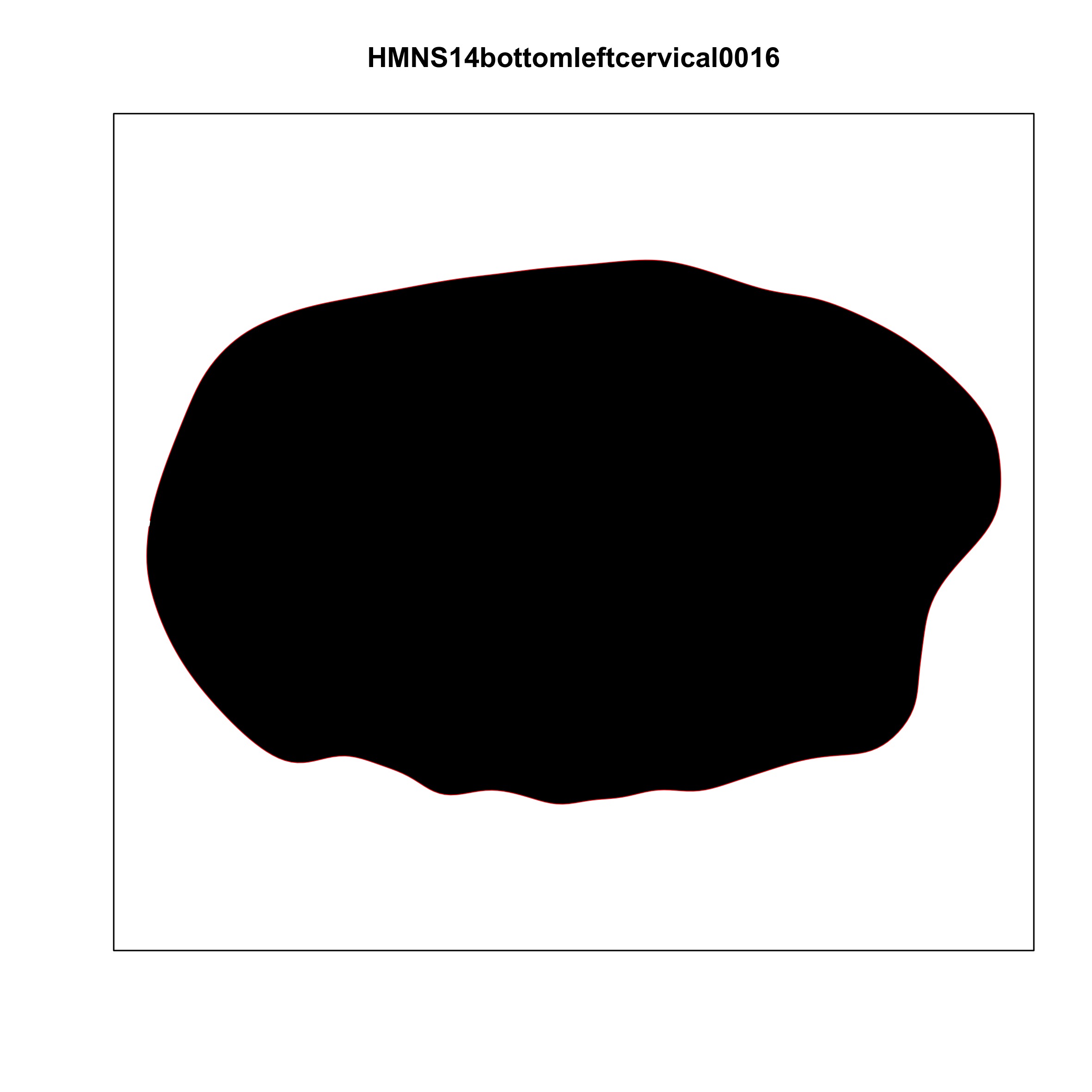

### plate_no_2.jpg

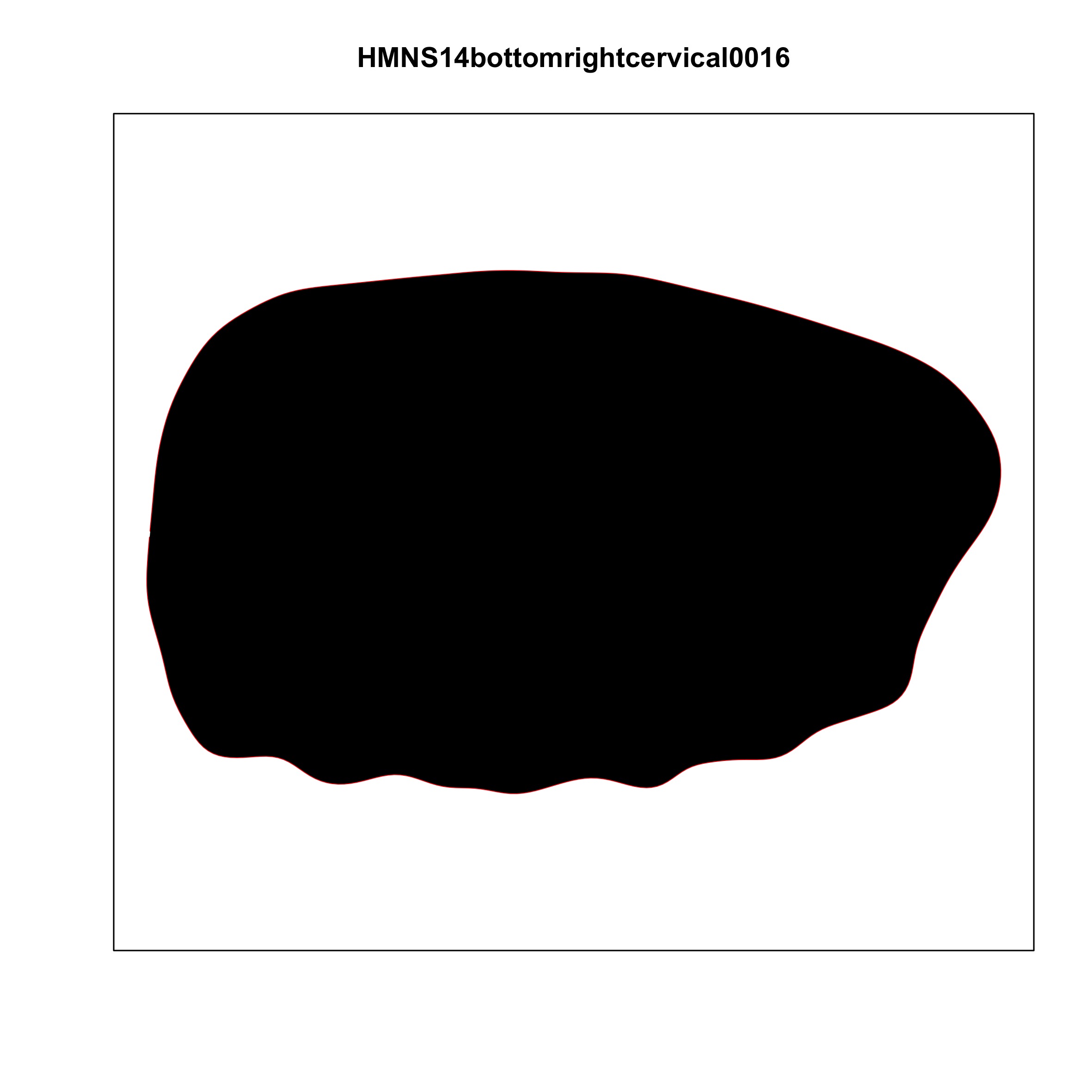

### plate_no_3.jpg

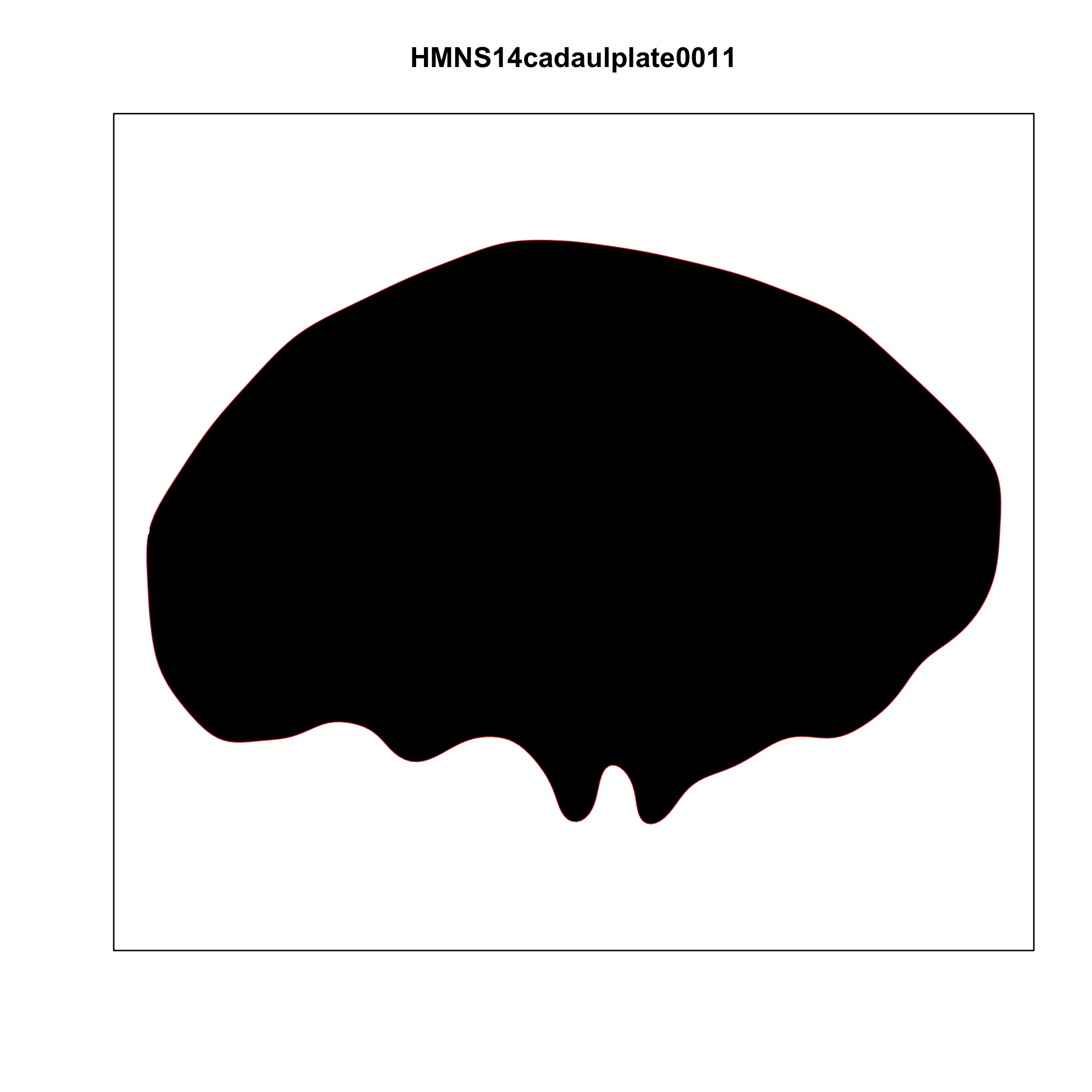

### plate_no_4.jpg

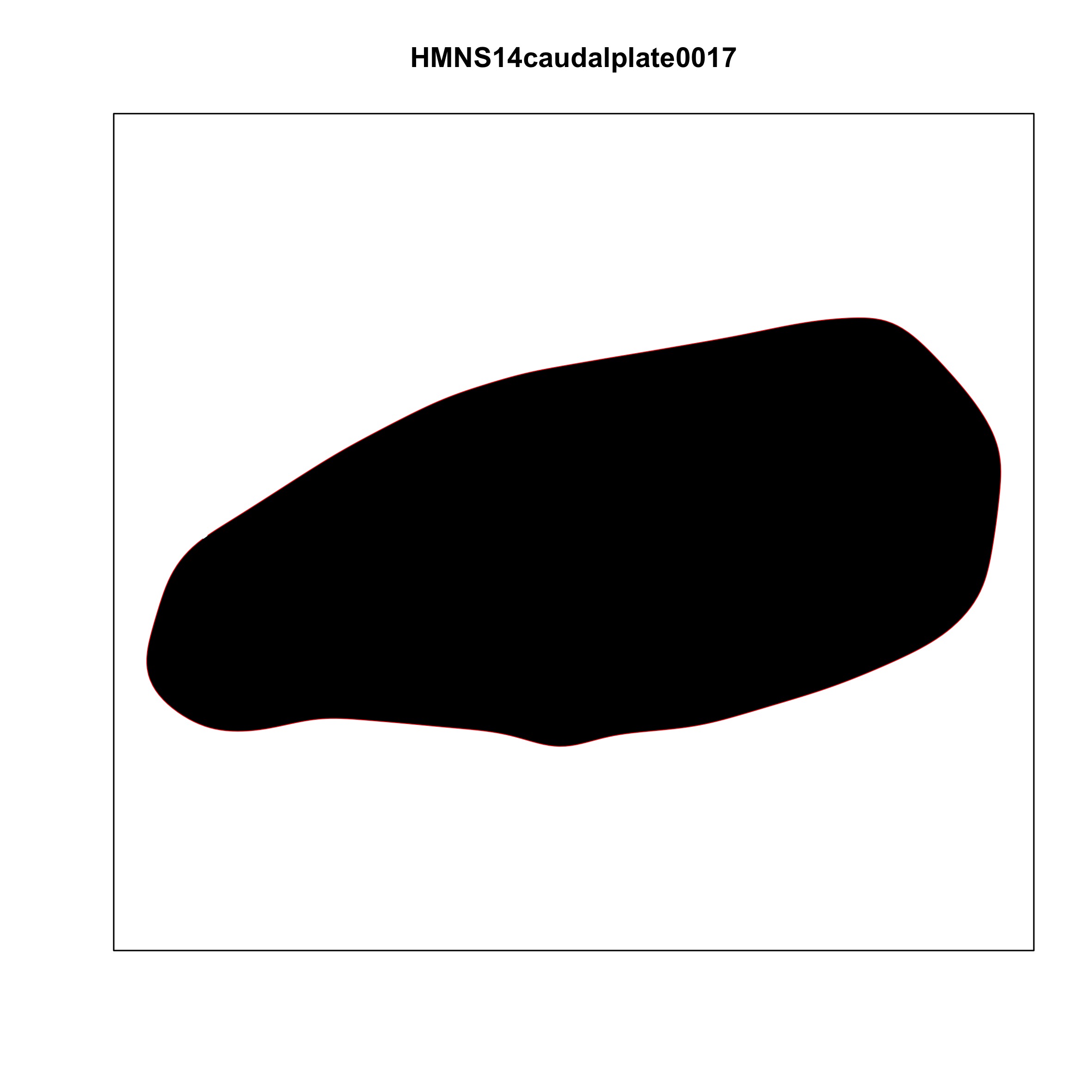

### plate_no_5.jpg

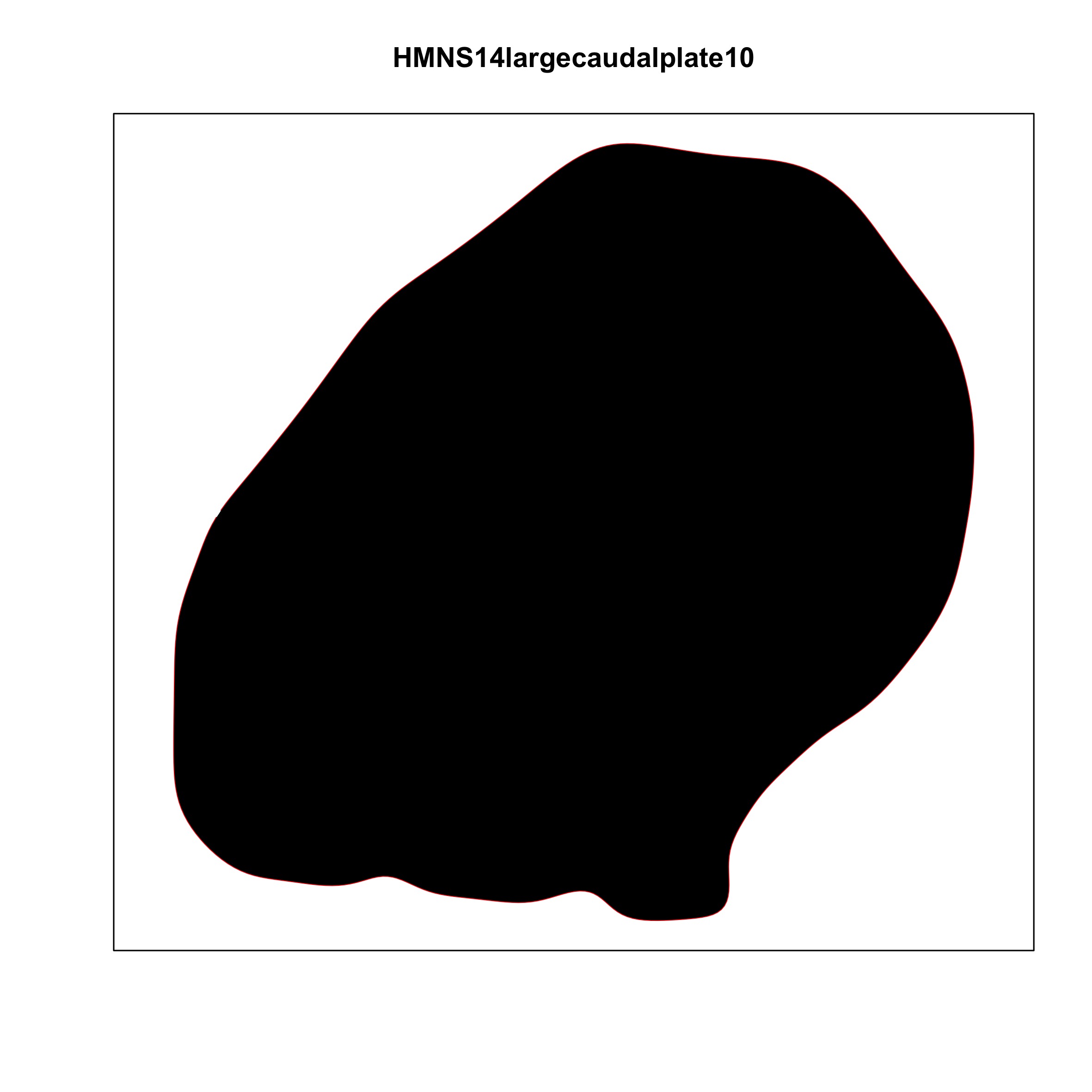

### plate_no_6.jpg

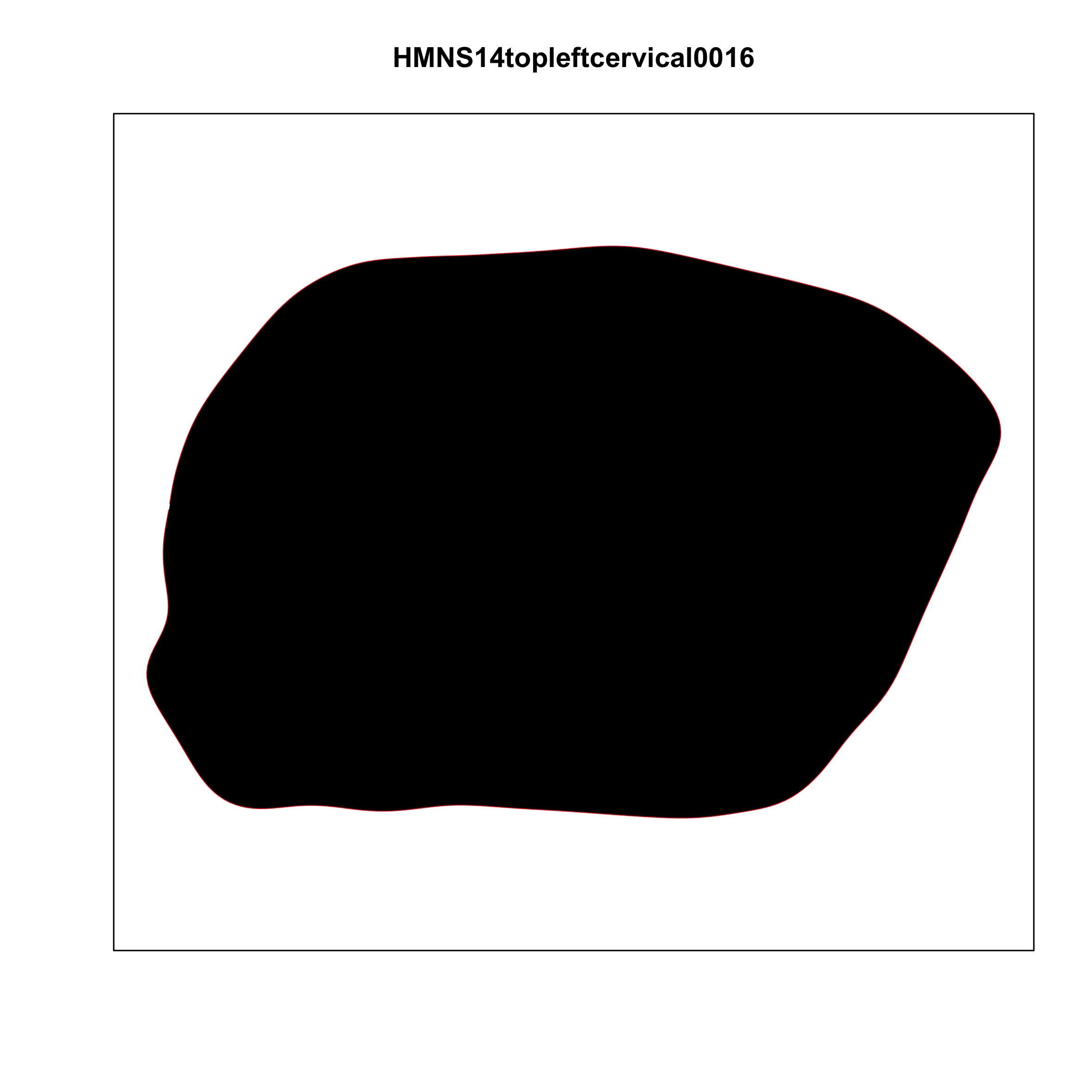

### plate_no_7.jpg

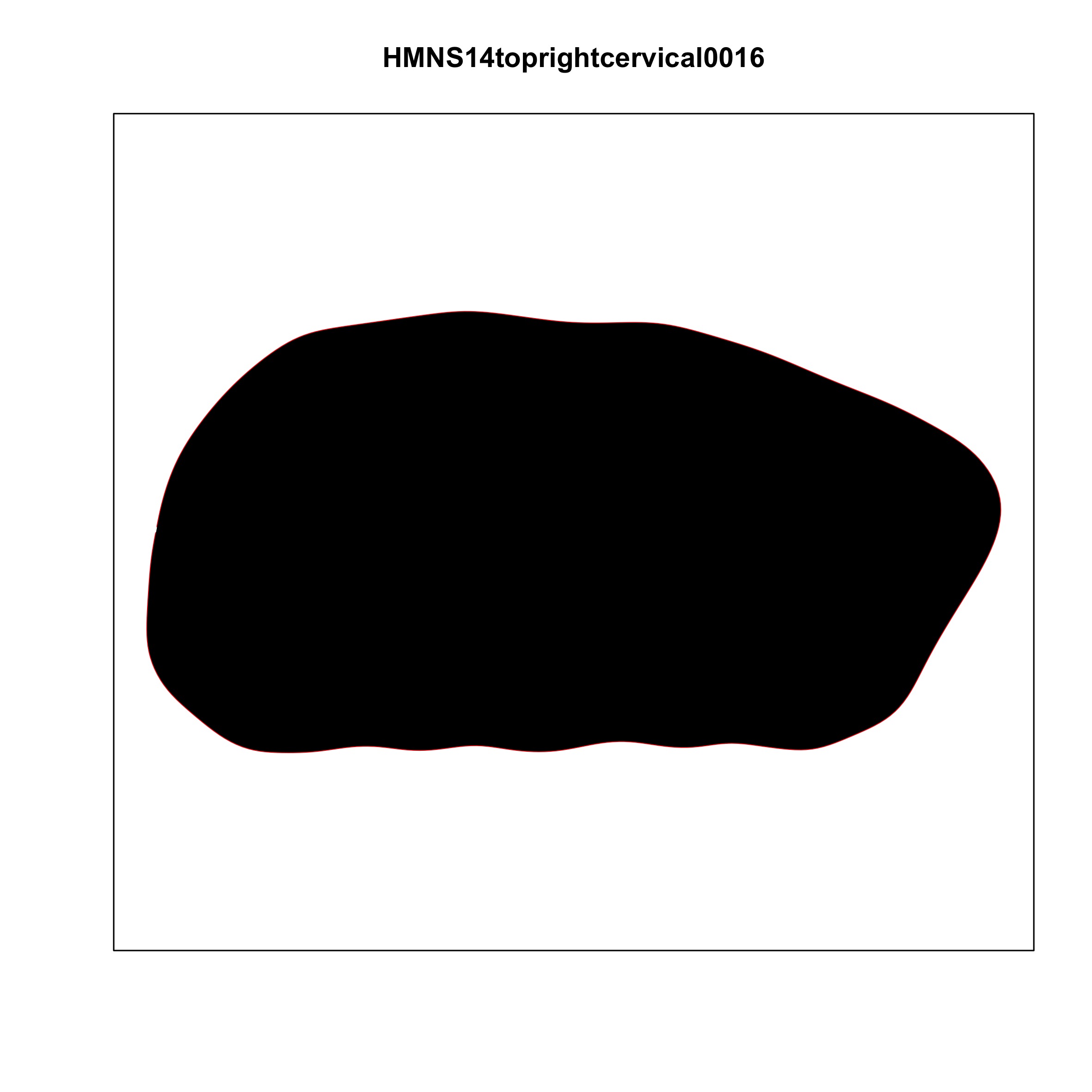

### plate_no_8.jpg

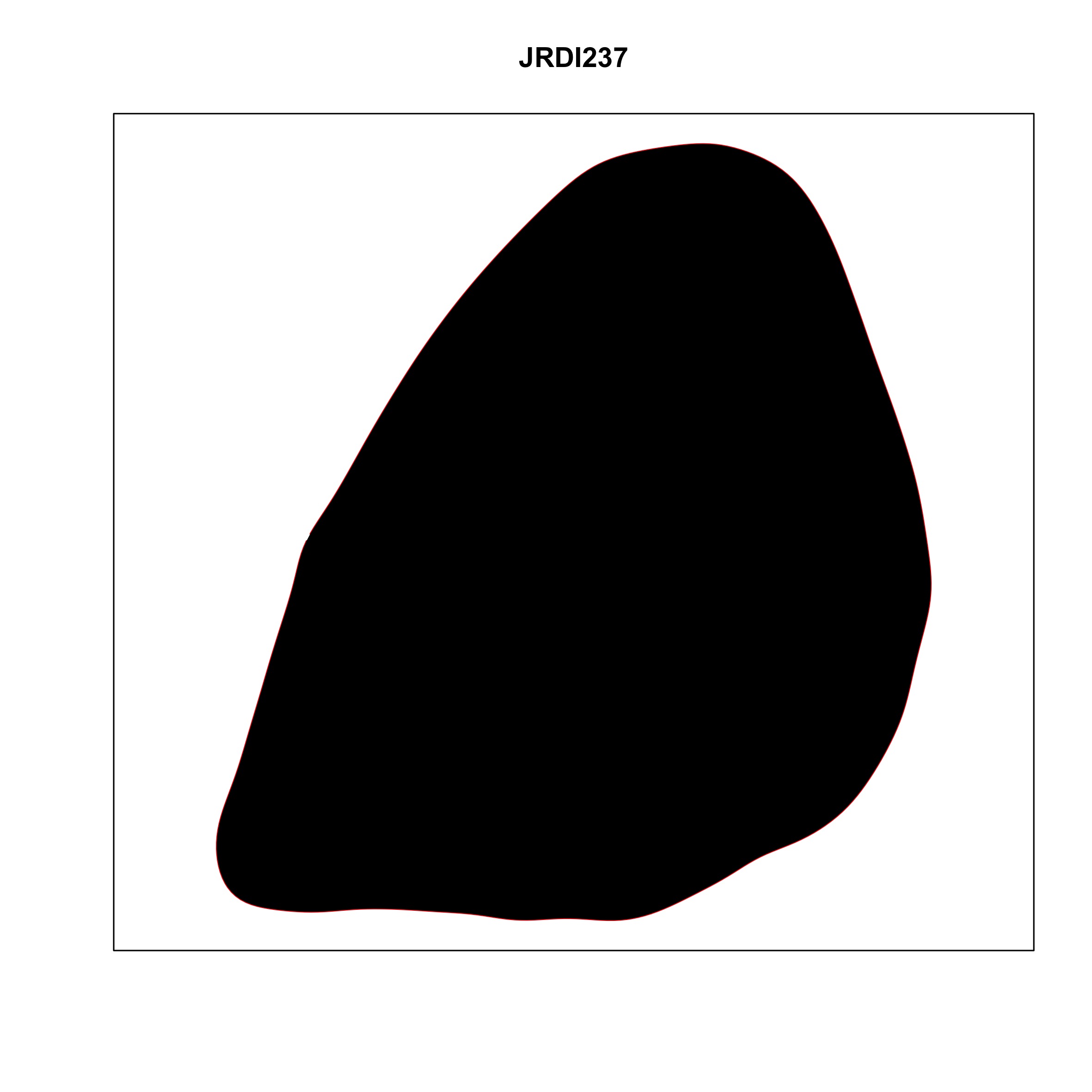

### plate_no_9.jpg

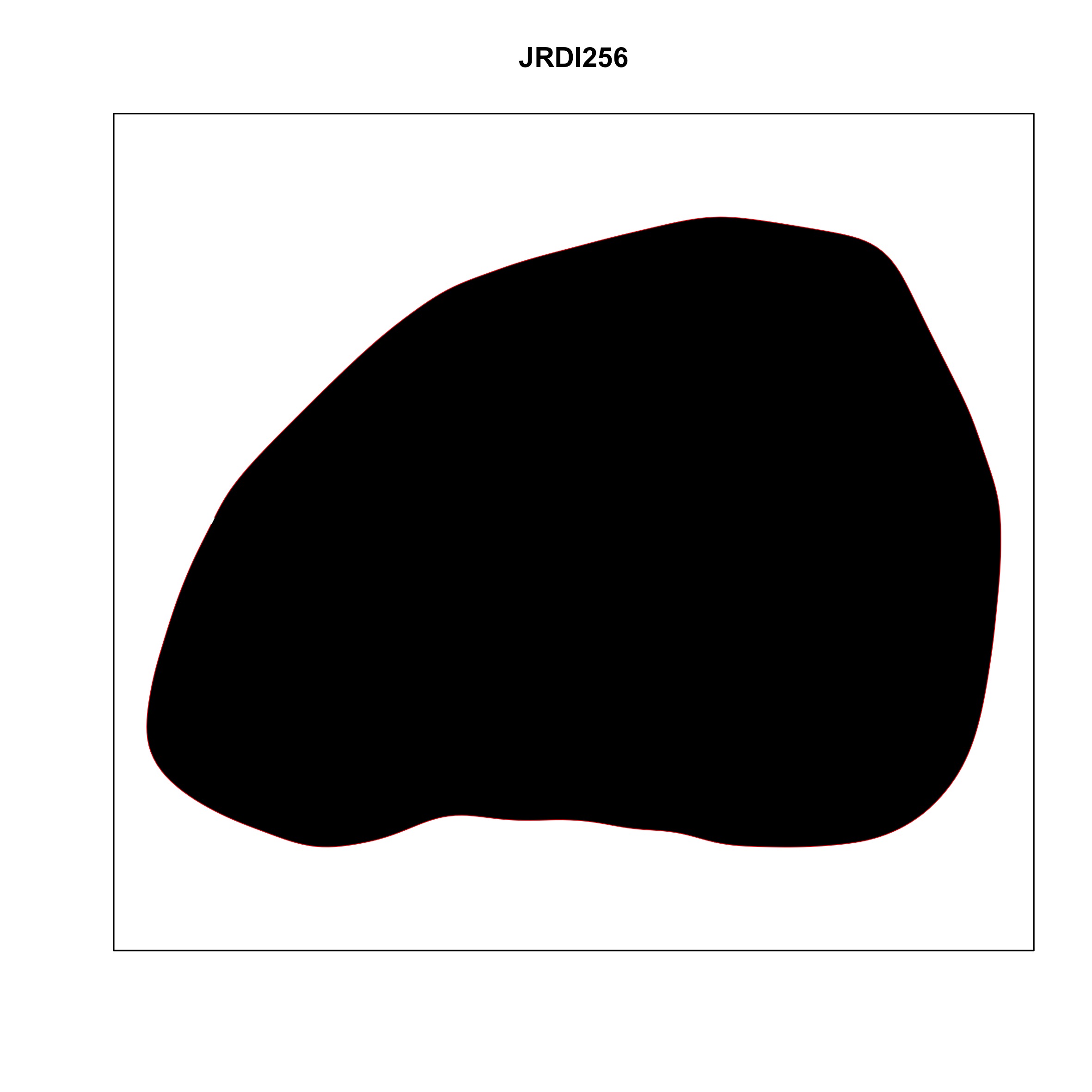

### plate_no_10.jpg

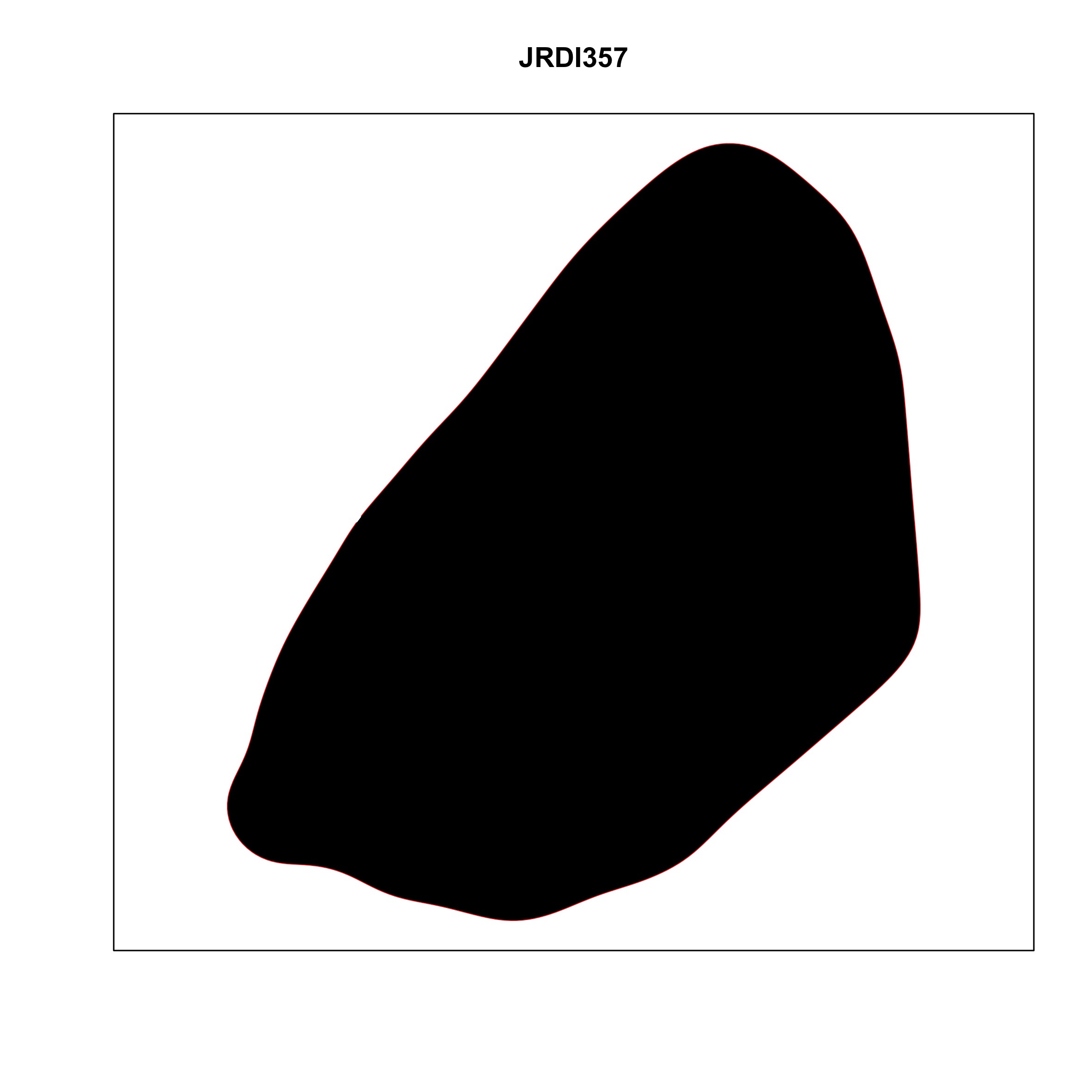

### plate_no_11.jpg

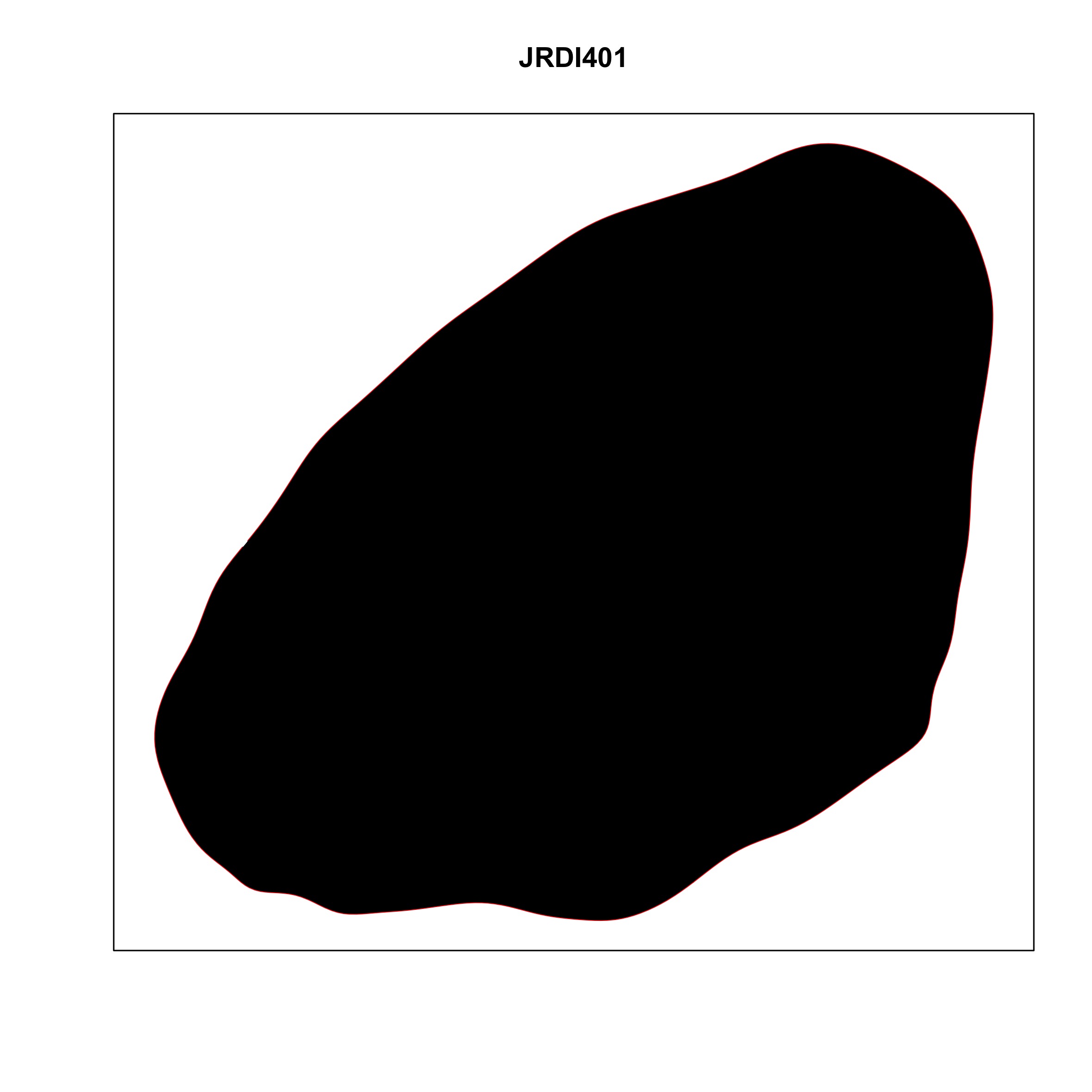

### plate_no_12.jpg

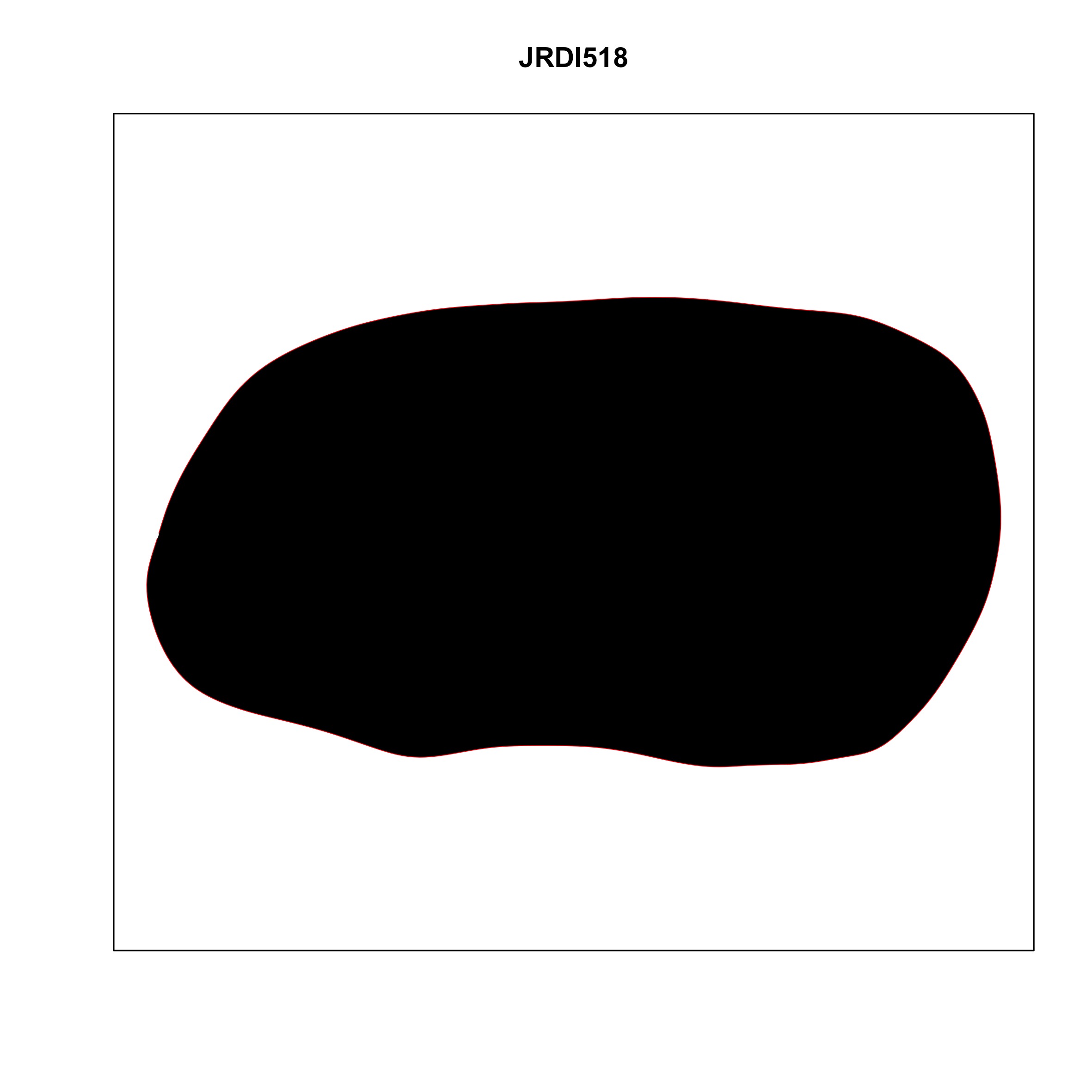

### plate_no_13.jpg

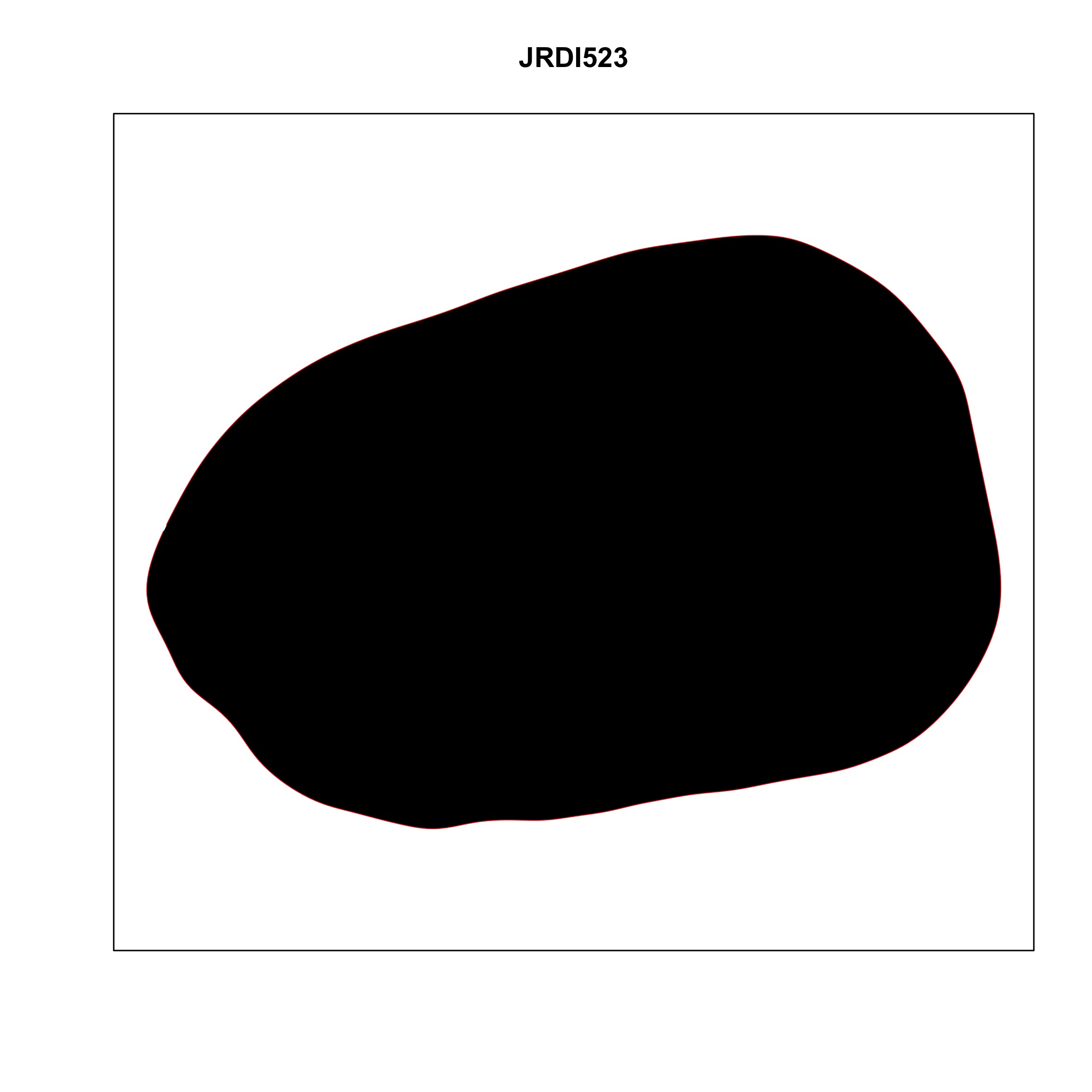

### plate_no_14.jpg

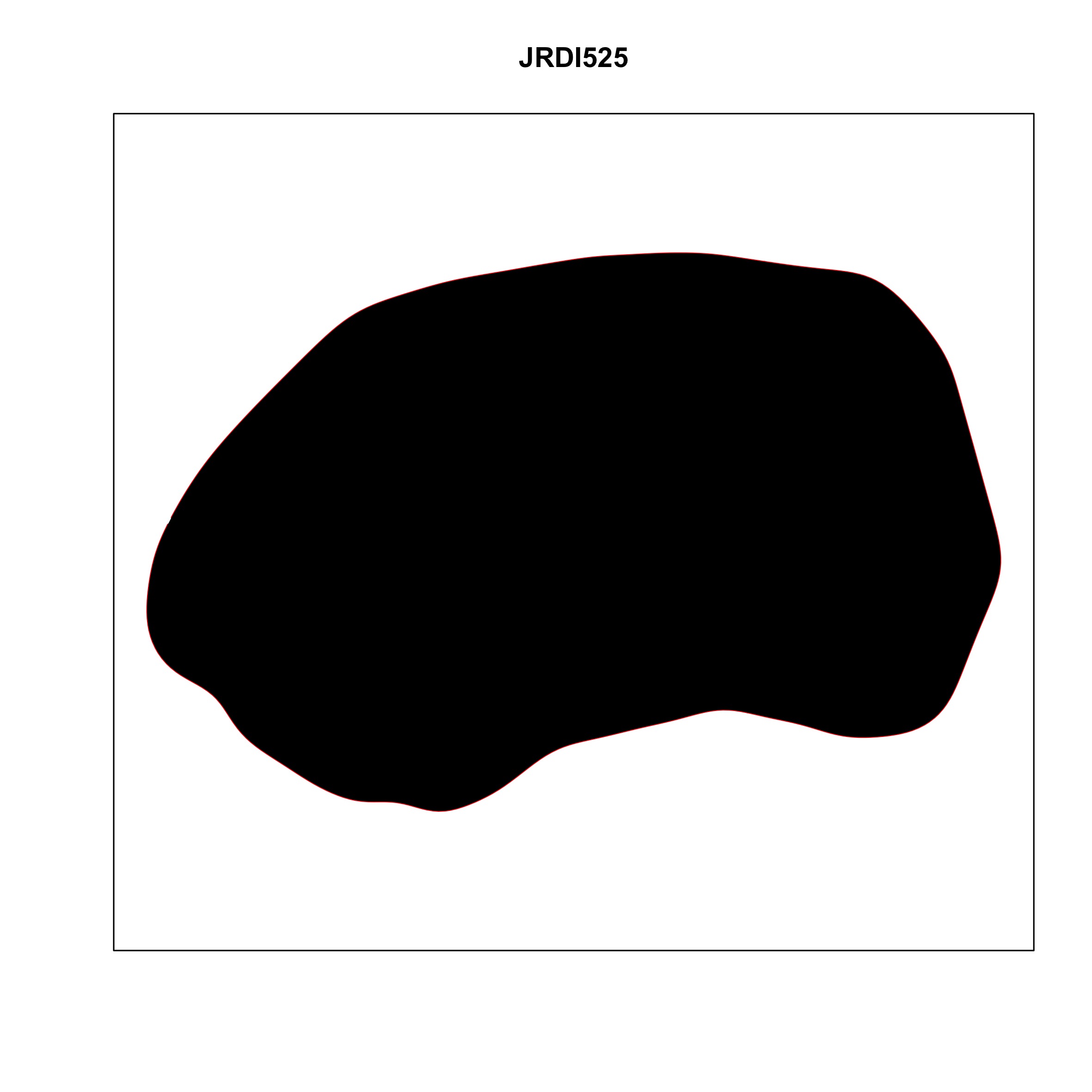

### plate_no_15.jpg

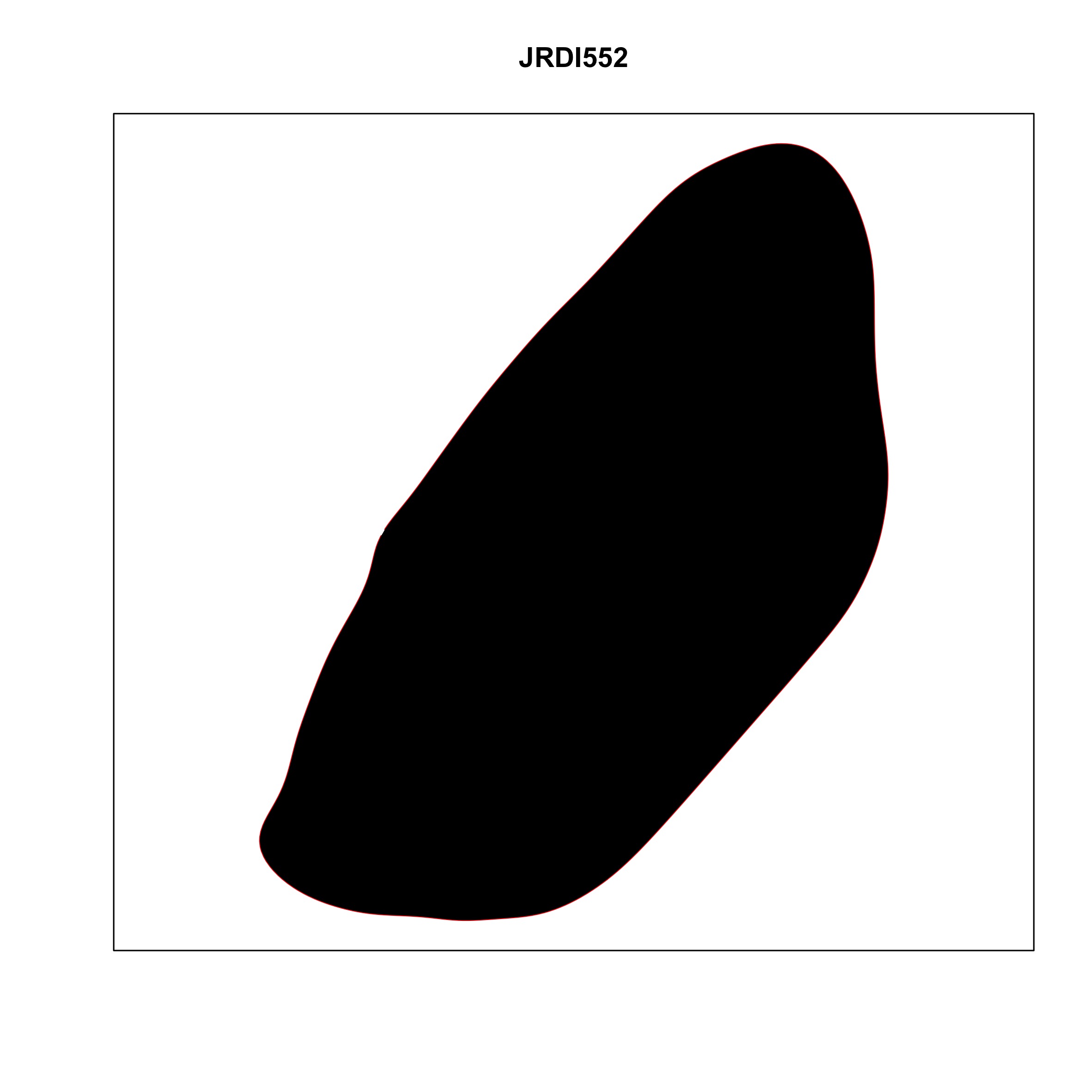

### plate_no_16.jpg

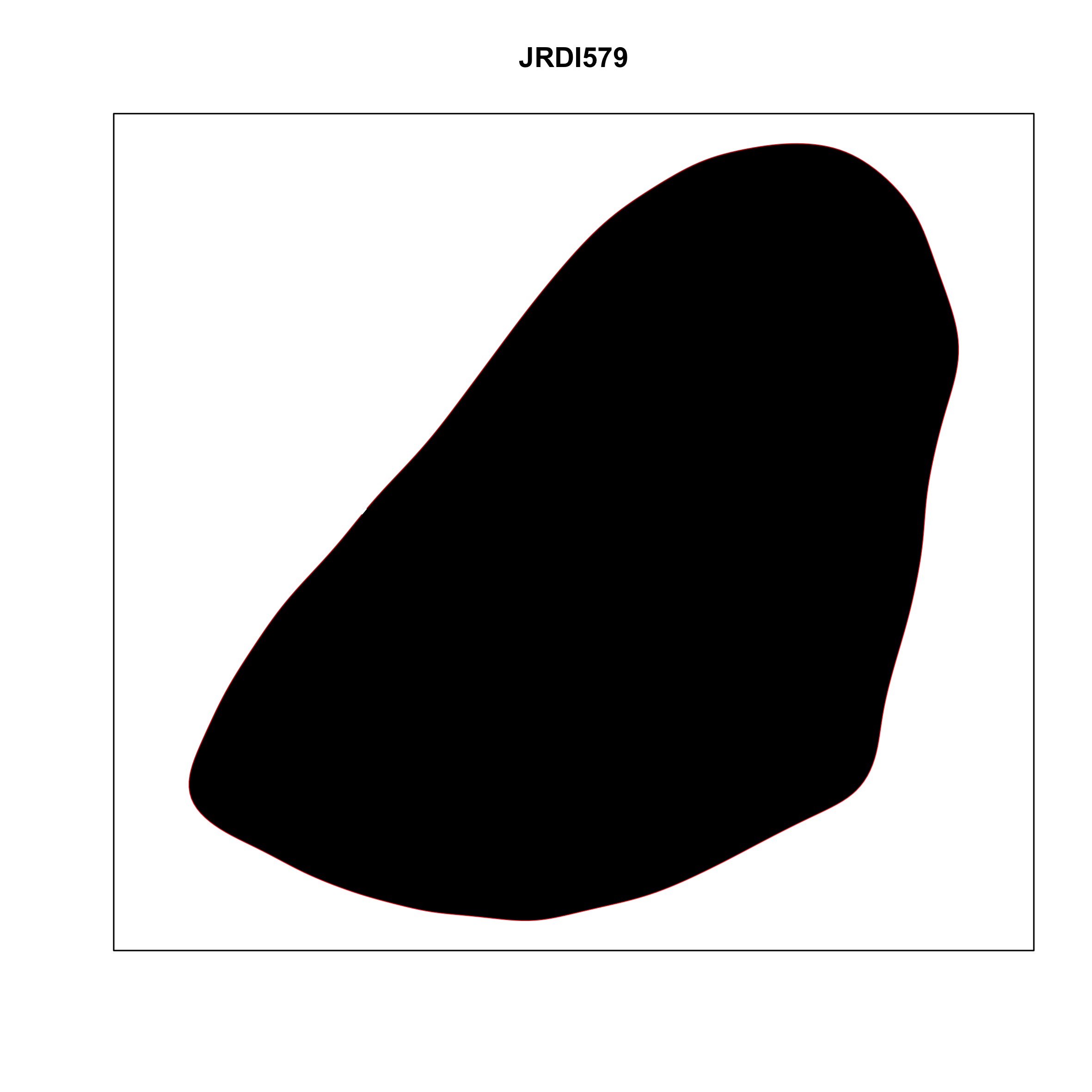

### plate_no_17.jpg

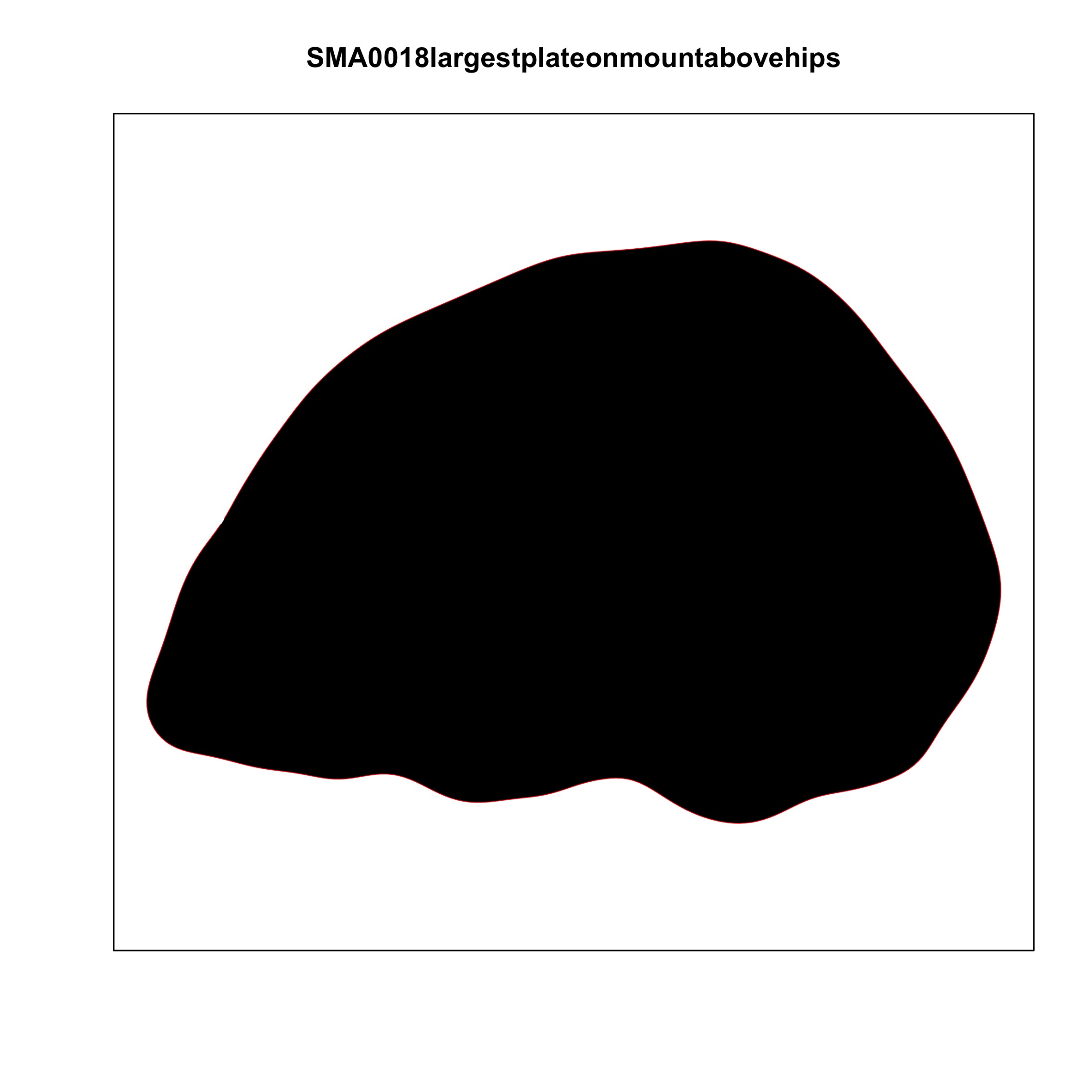

### plate_no_18.jpg

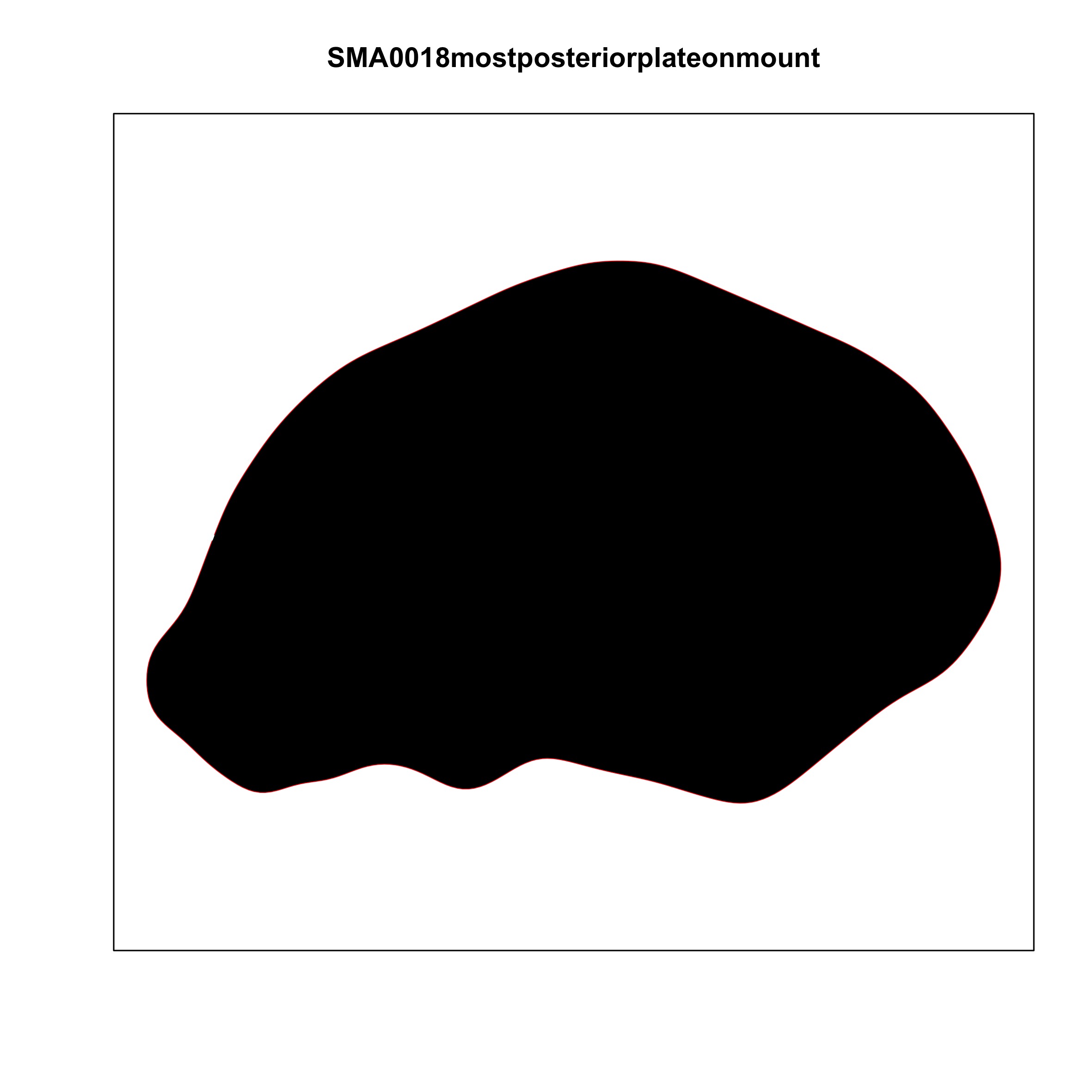

### plate_no_19.jpg

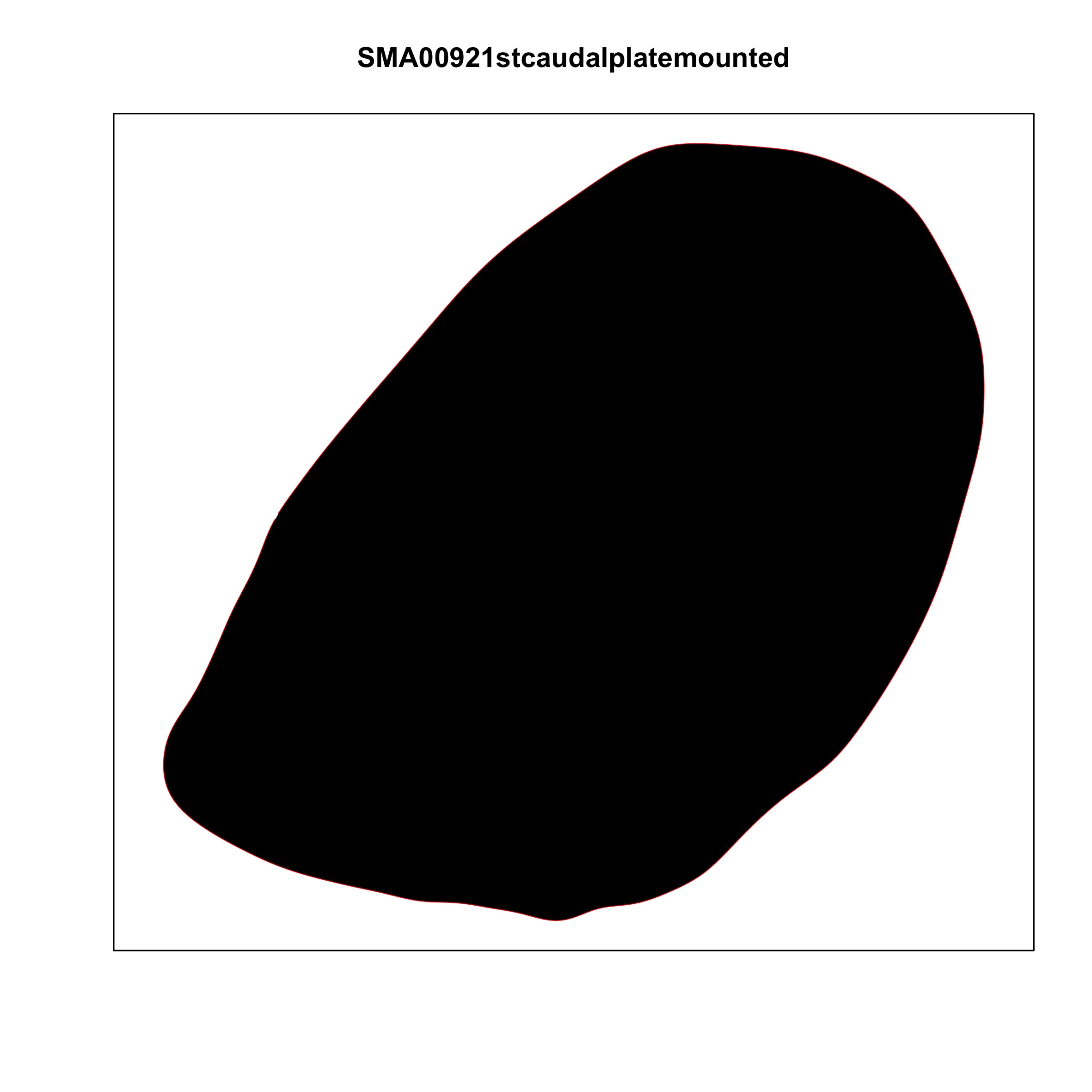

### plate_no_20.jpg

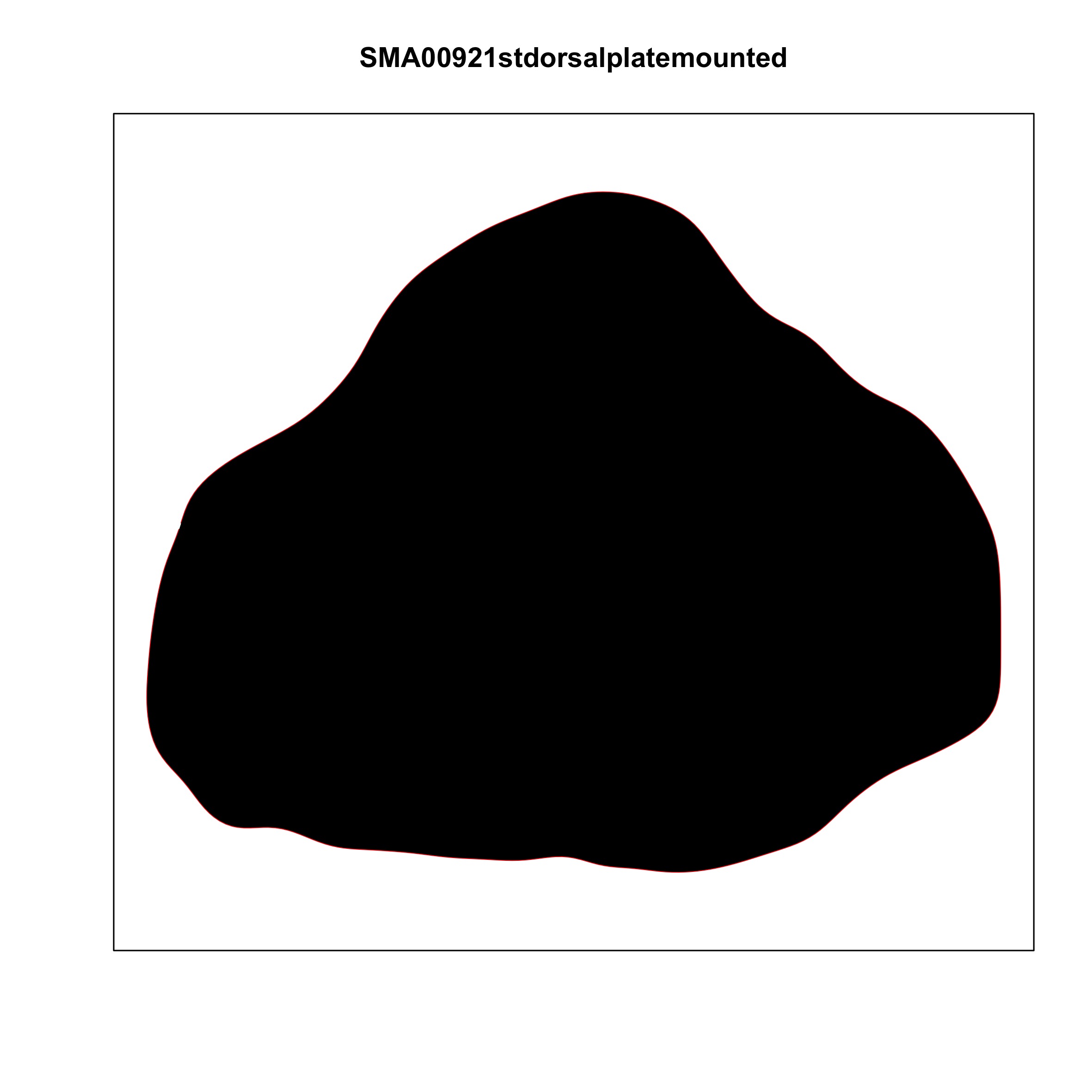

### plate_no_21.jpg

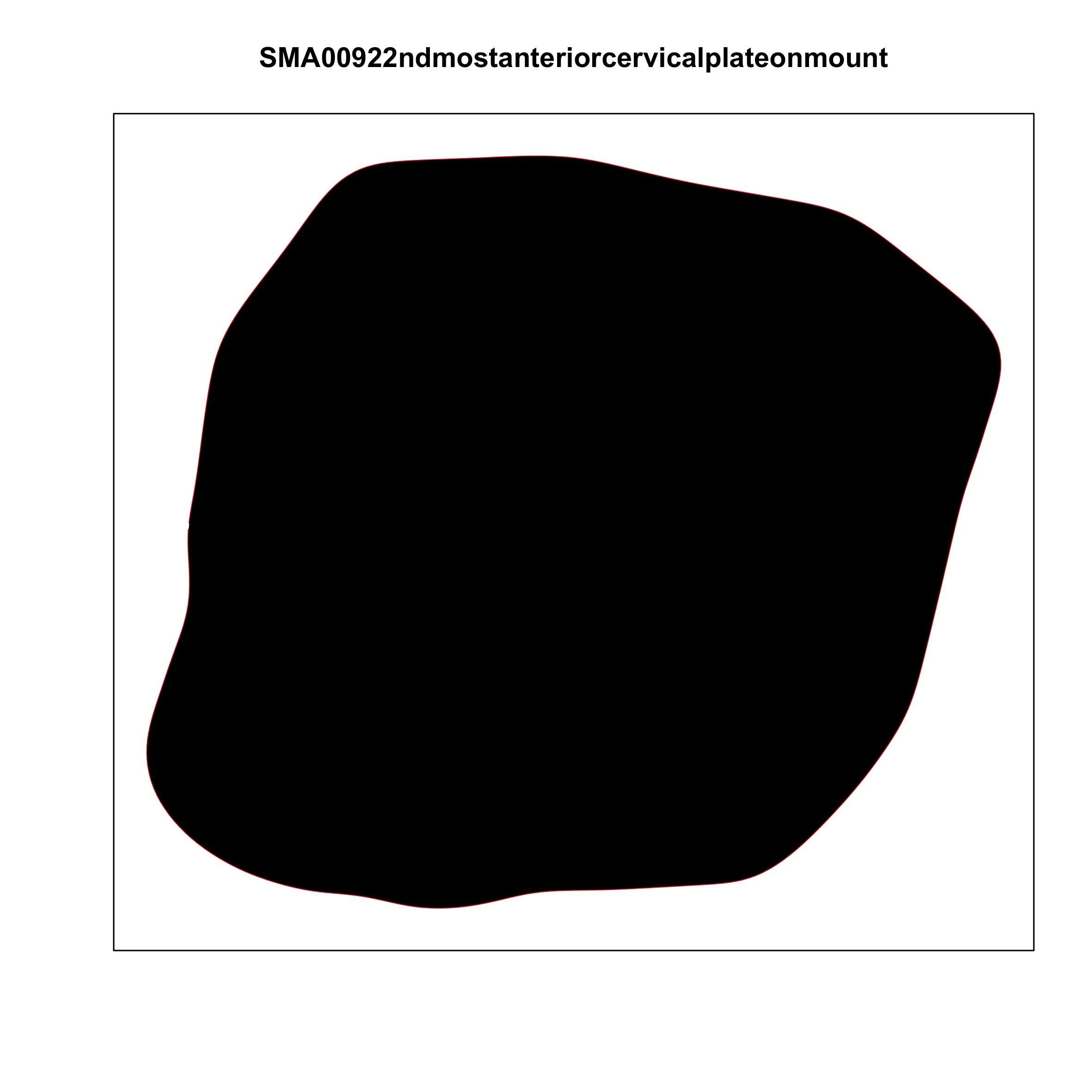

### plate_no_22.jpg

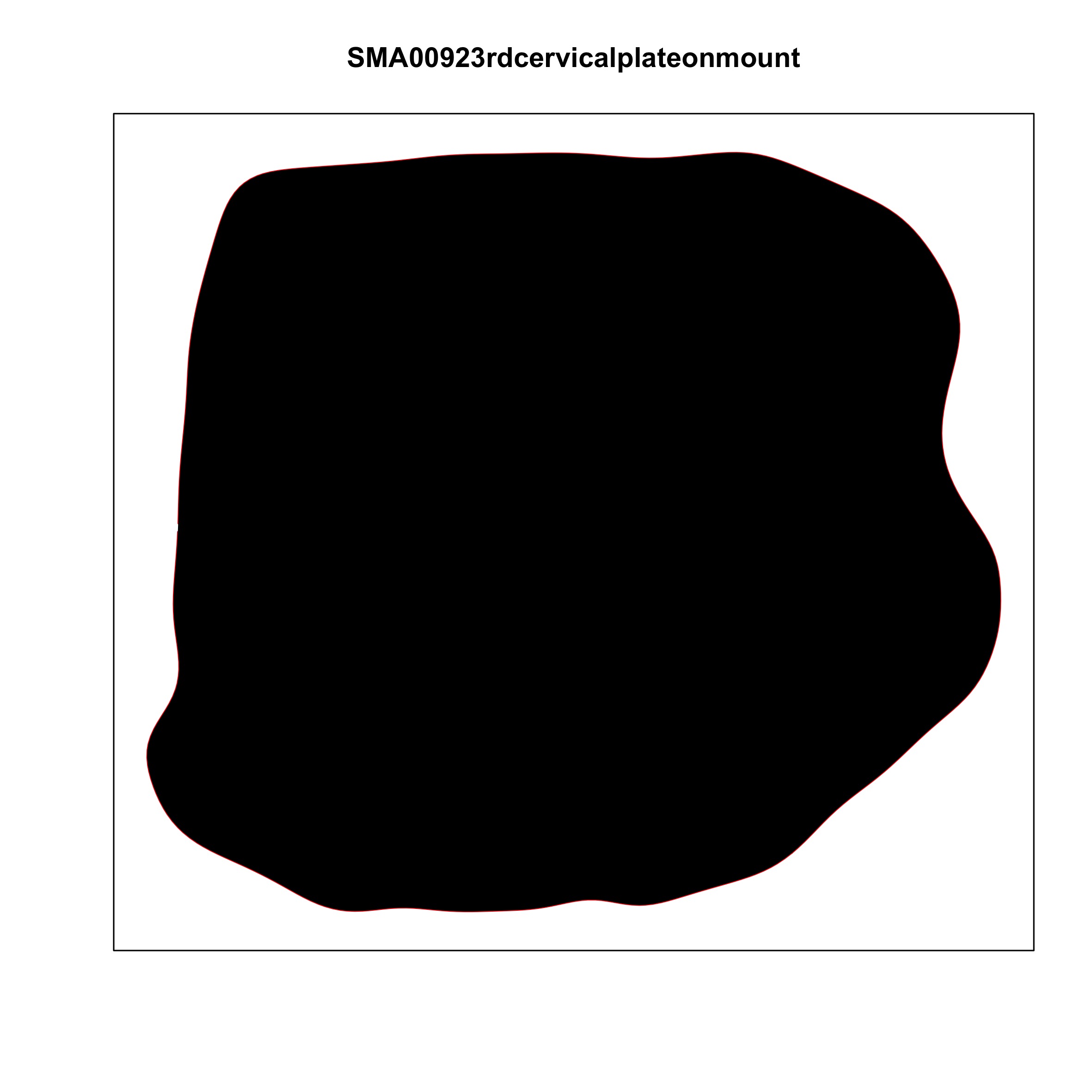

### plate_no_23.jpg

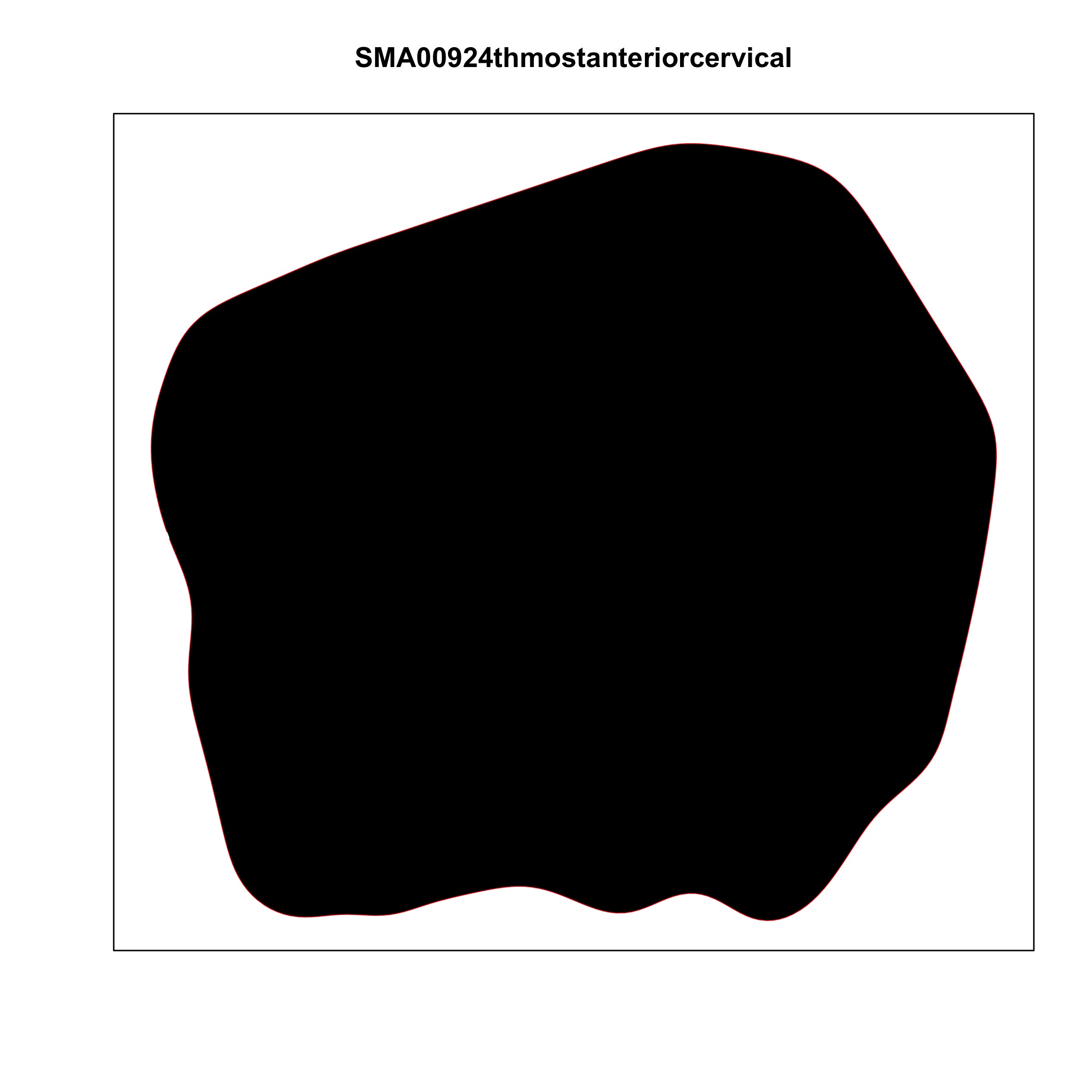

### plate_no_24.jpg

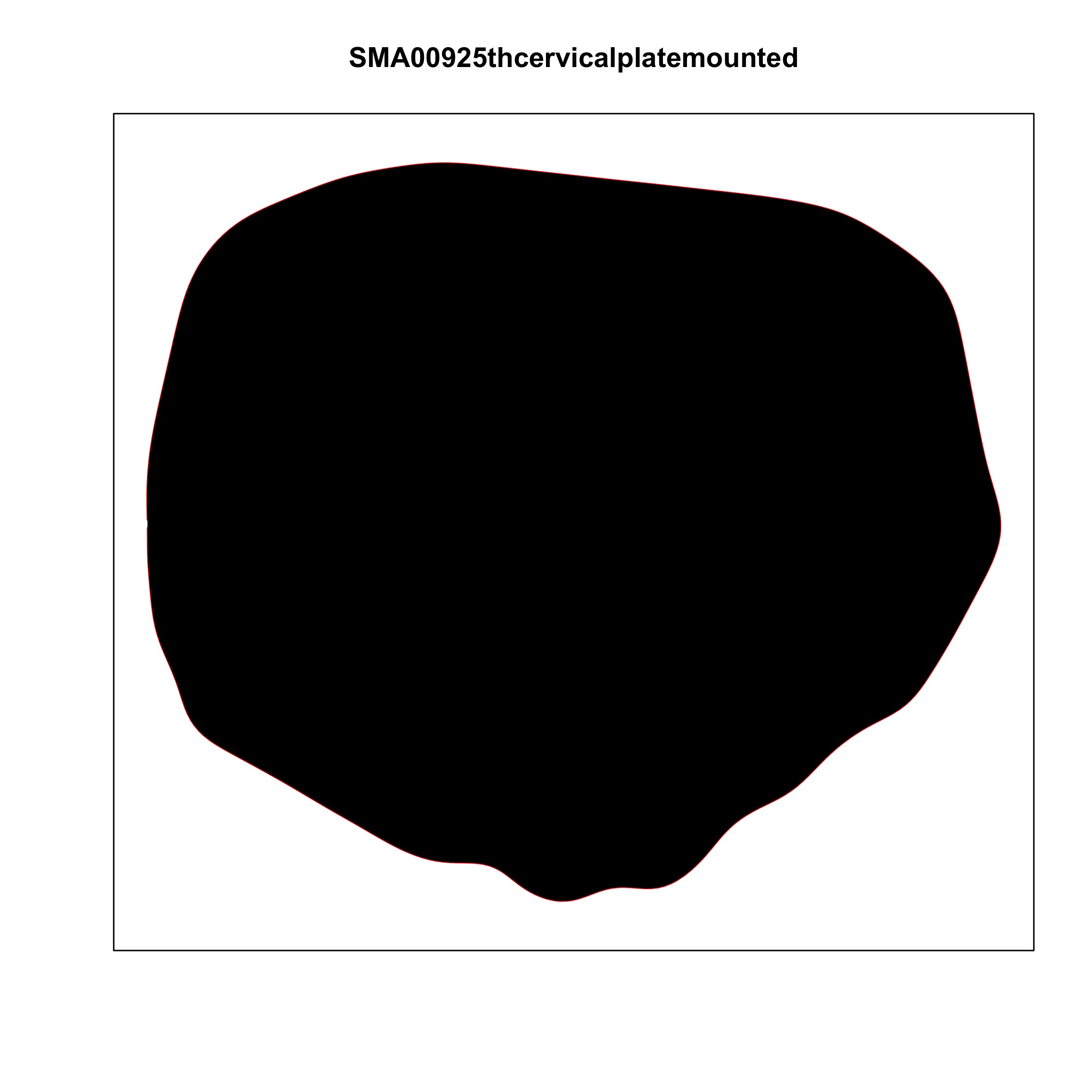

### plate_no_25.jpg

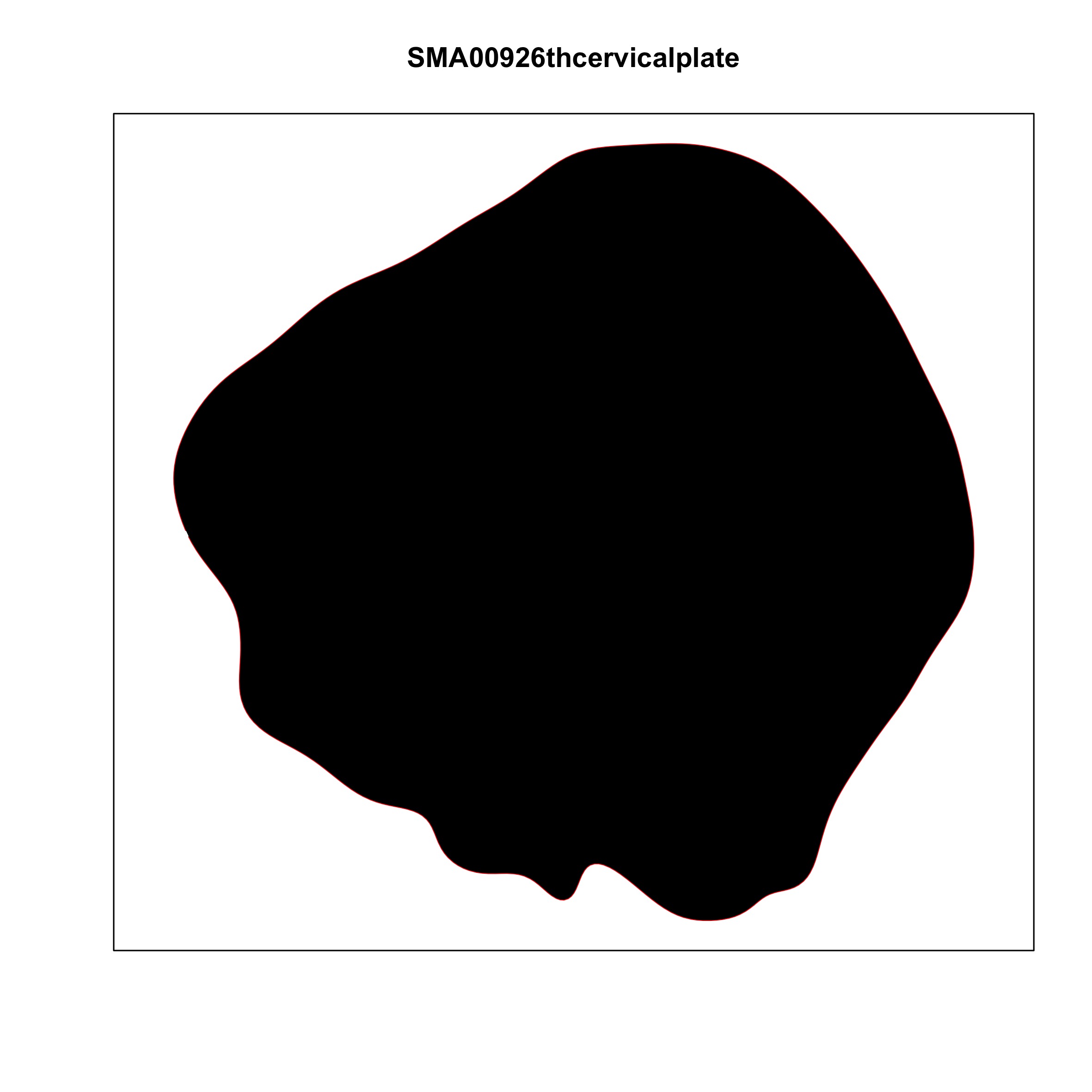

### plate_no_26.jpg

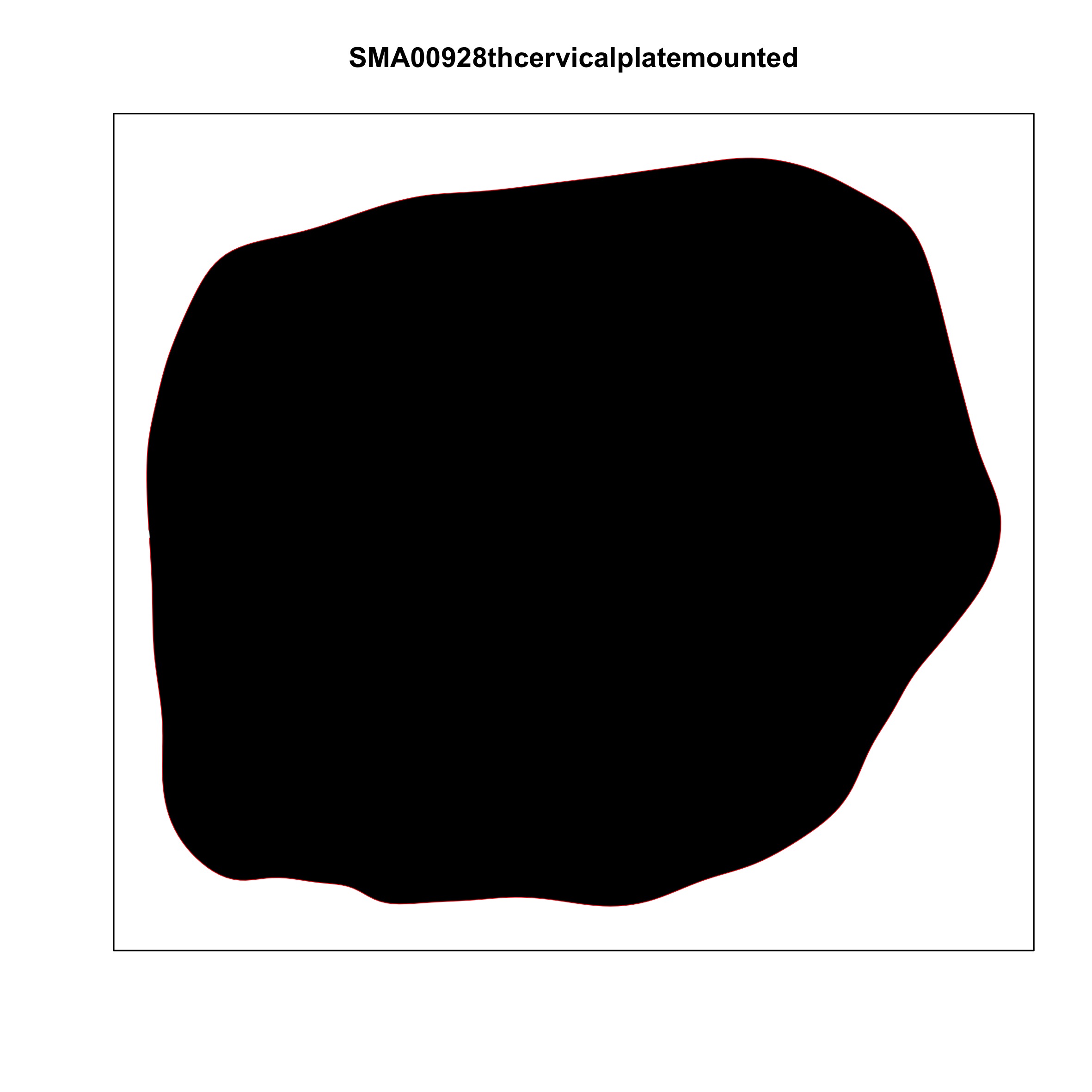

### plate_no_27.jpg

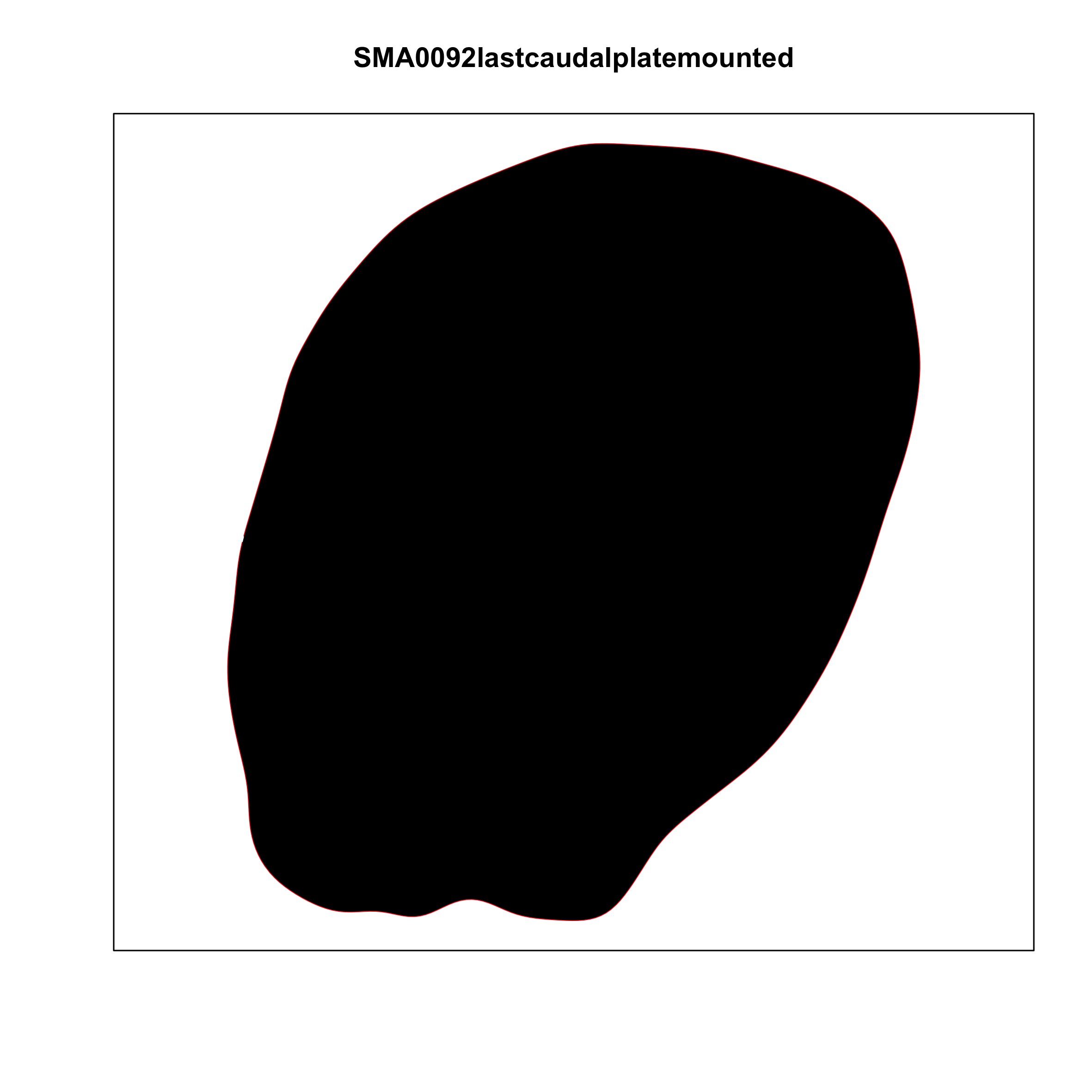

### plate_no_28.jpg

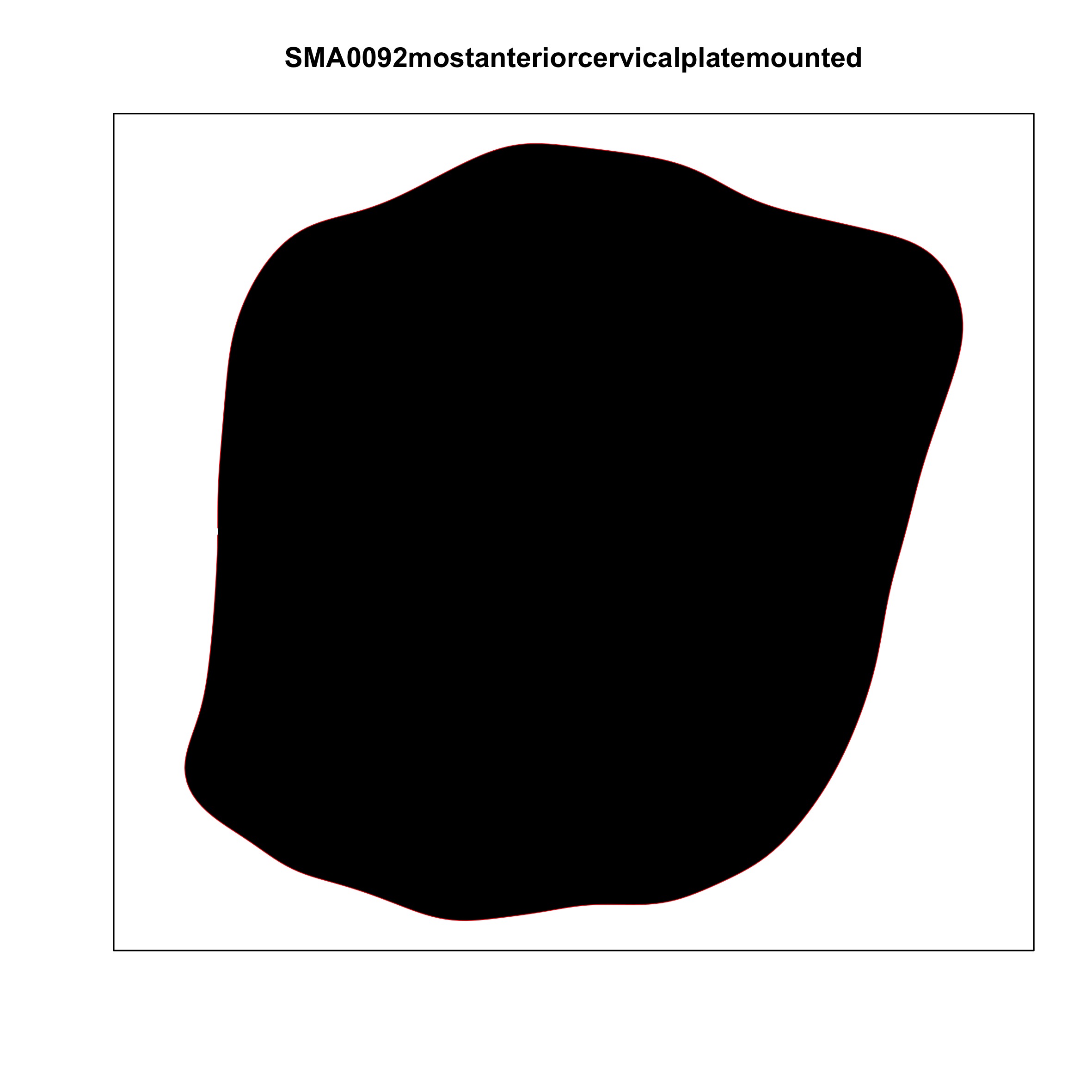

### plate_no_29.jpg

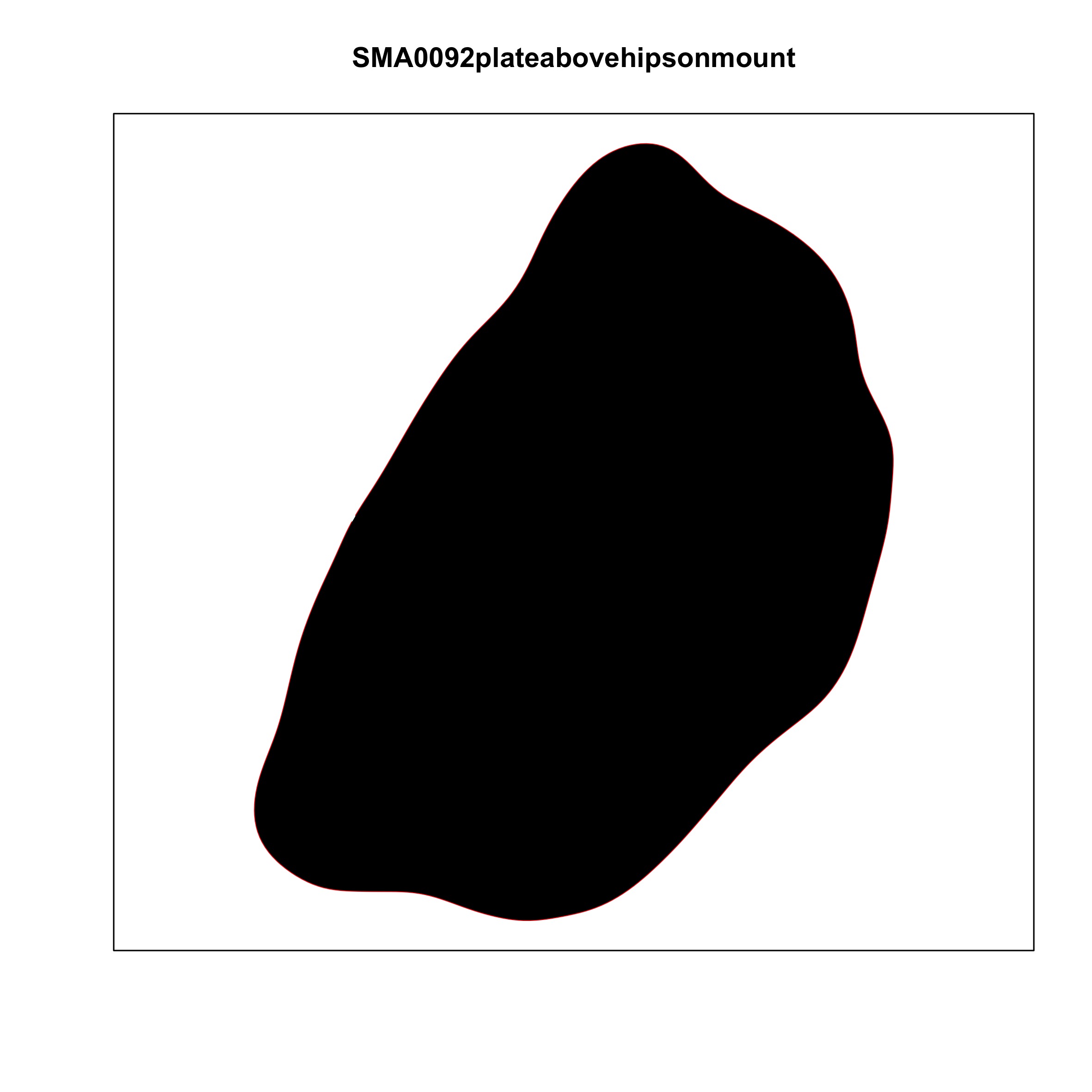

### plate_no_30.jpg

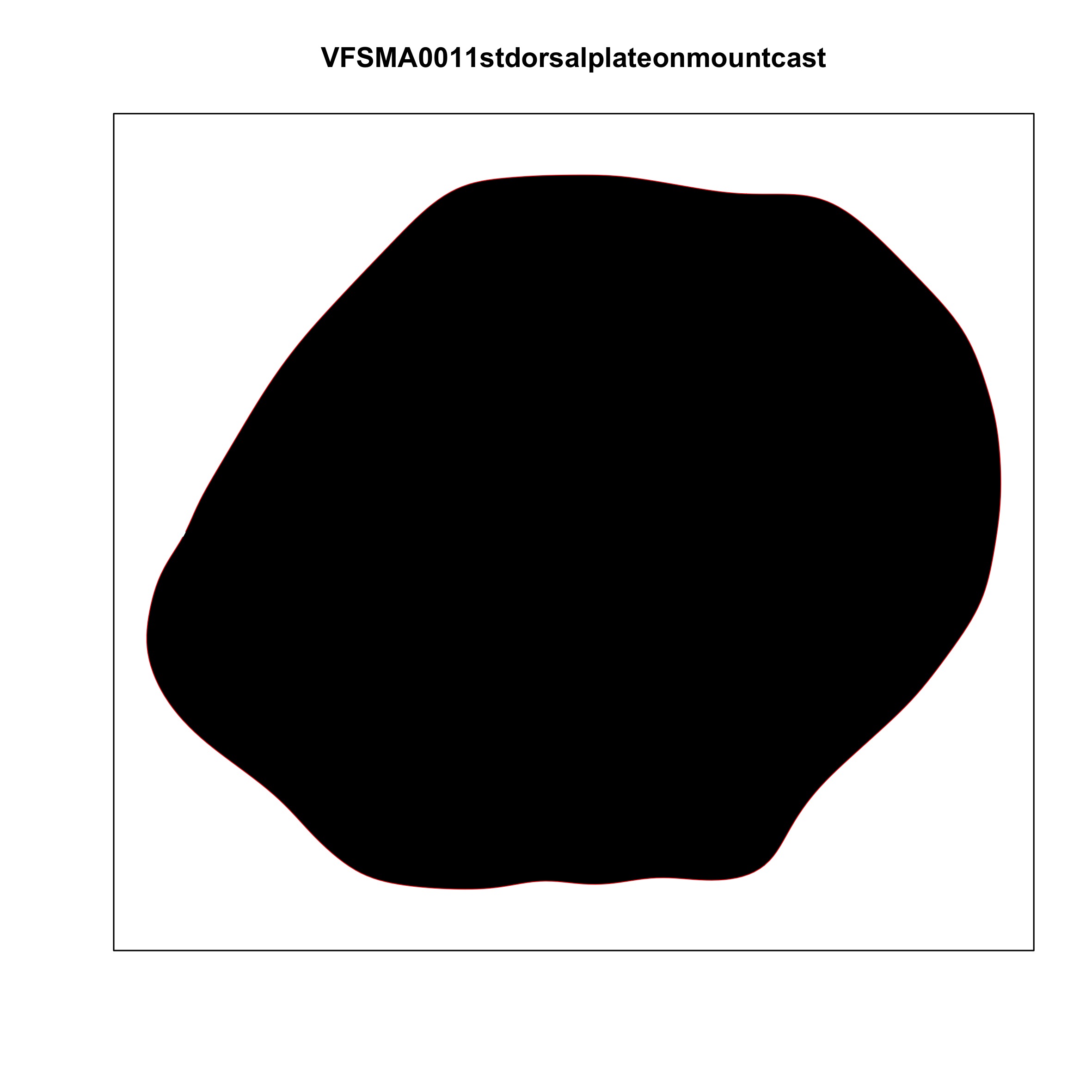

### plate_no_31.jpg

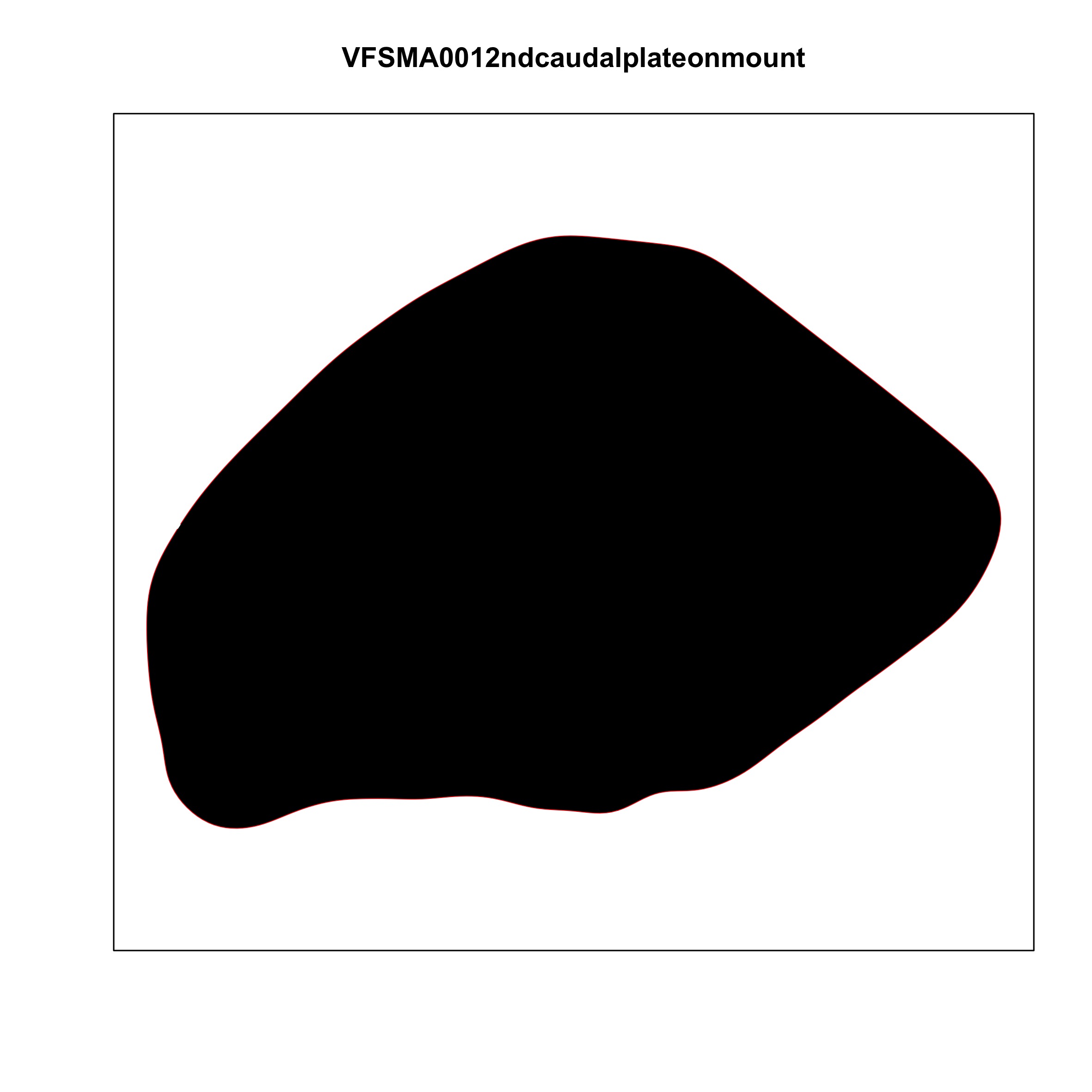

### plate_no_32.jpg

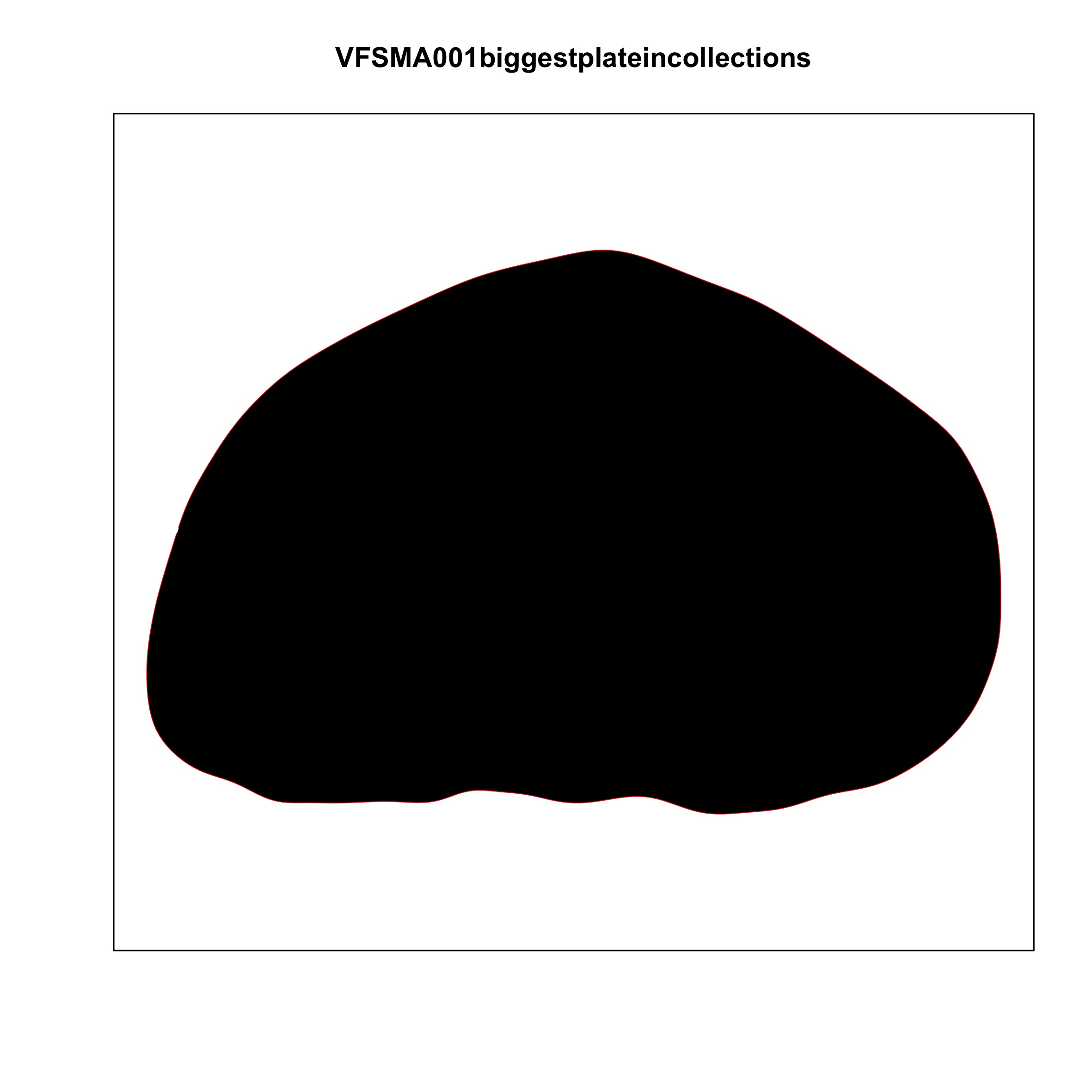

### plate_no_33.jpg

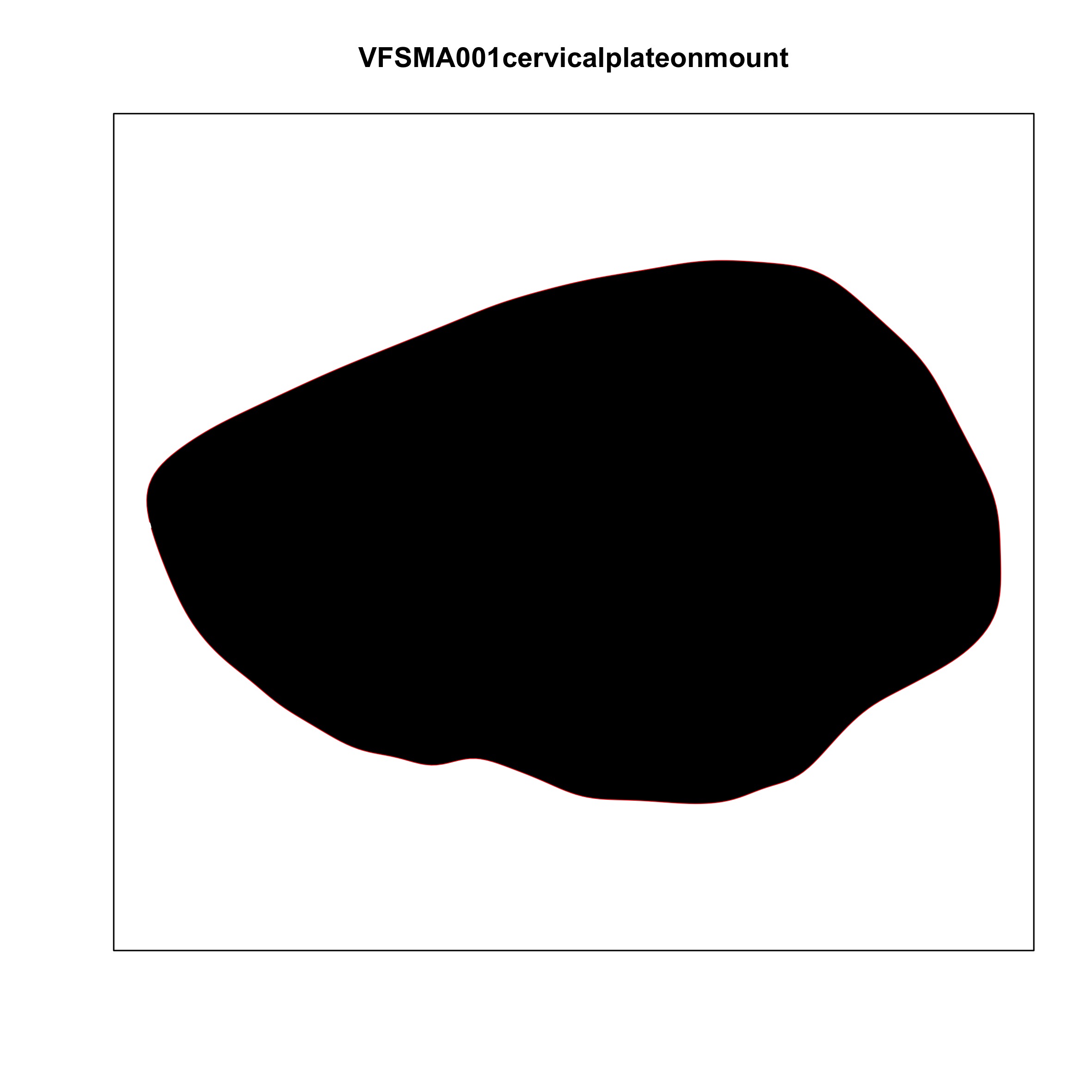

### plate_no_34.jpg

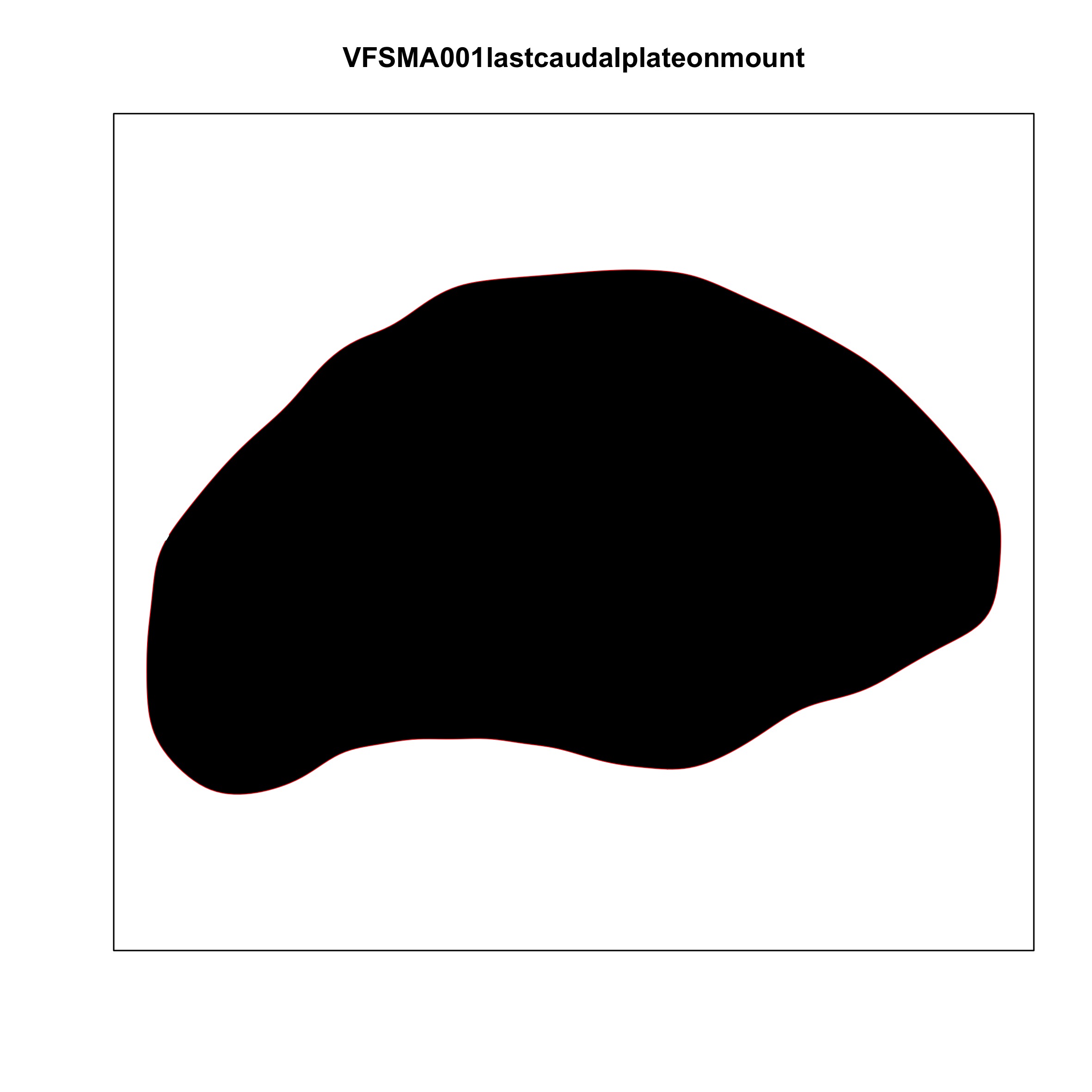

### plate_no_35.jpg

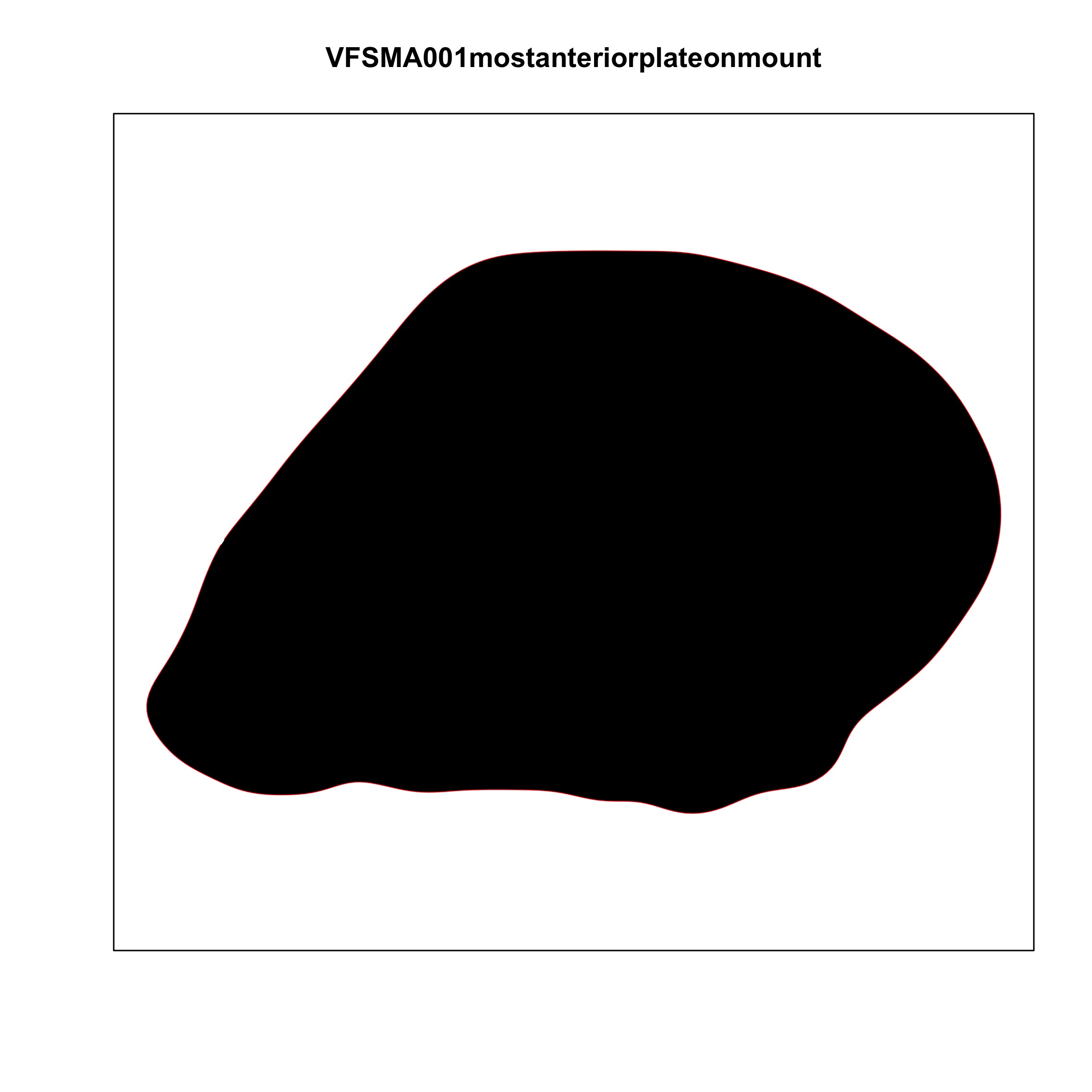

### plate_no_36.jpg

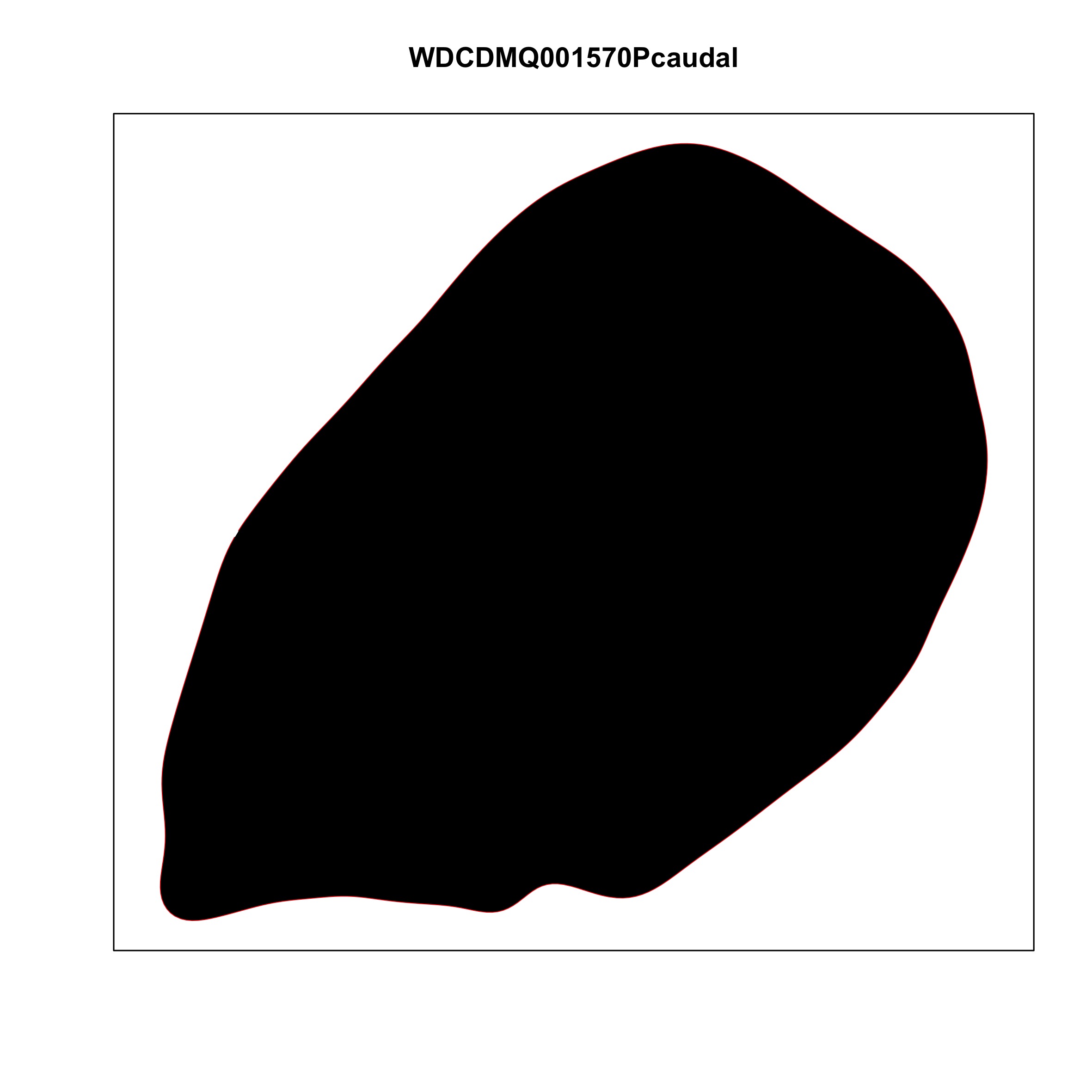

### plate_no_37.jpg

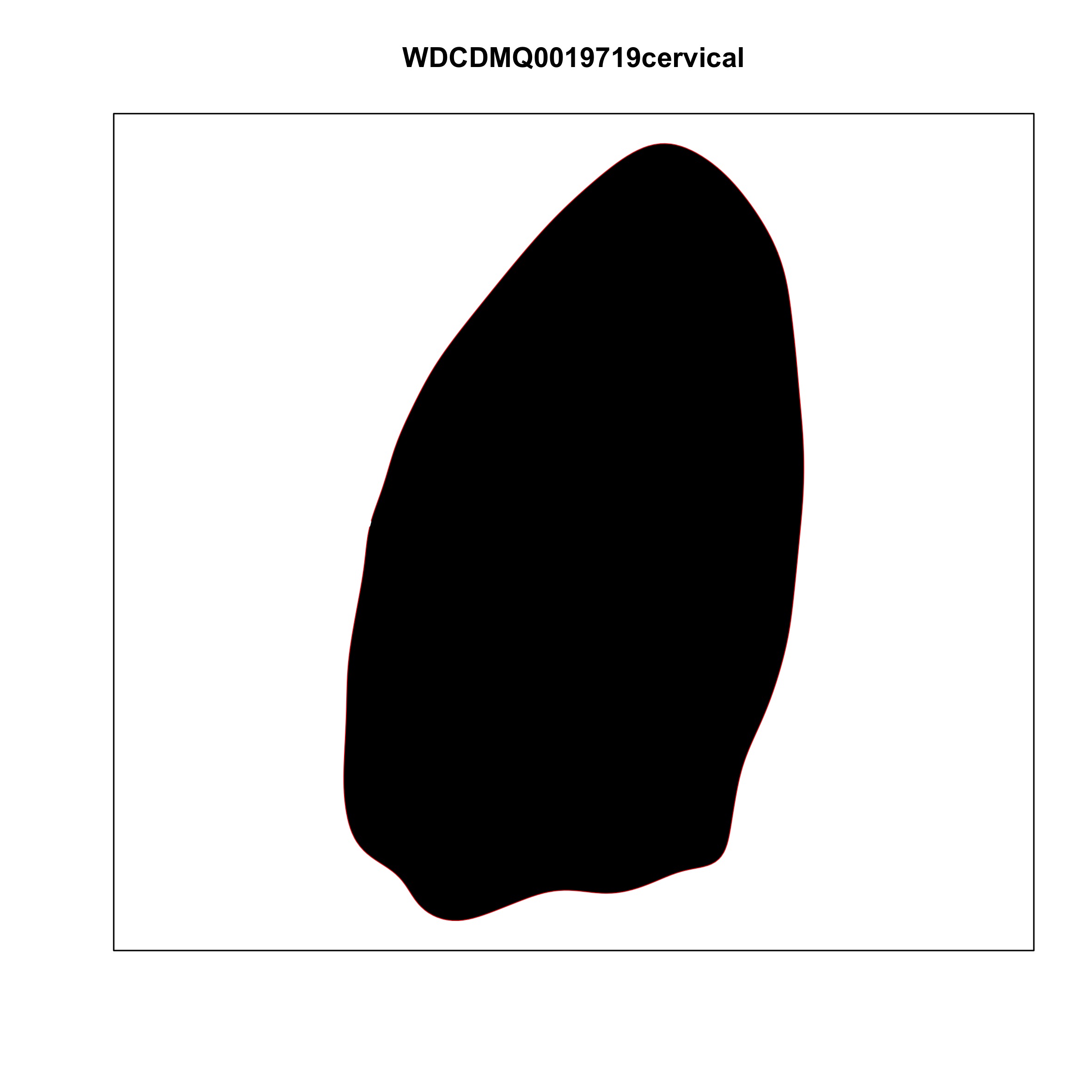

### plate_no_38.jpg

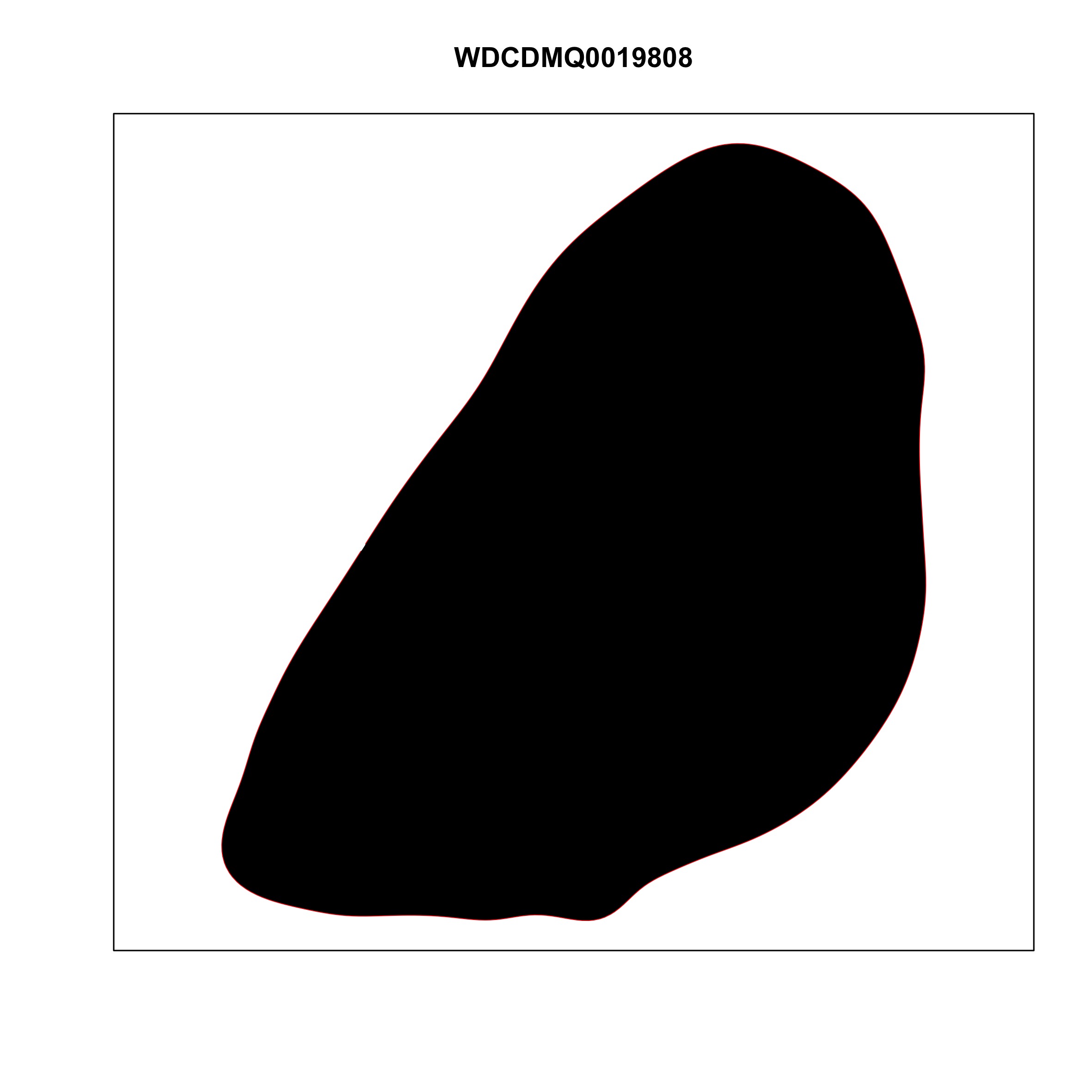

### plate_no_39.jpg

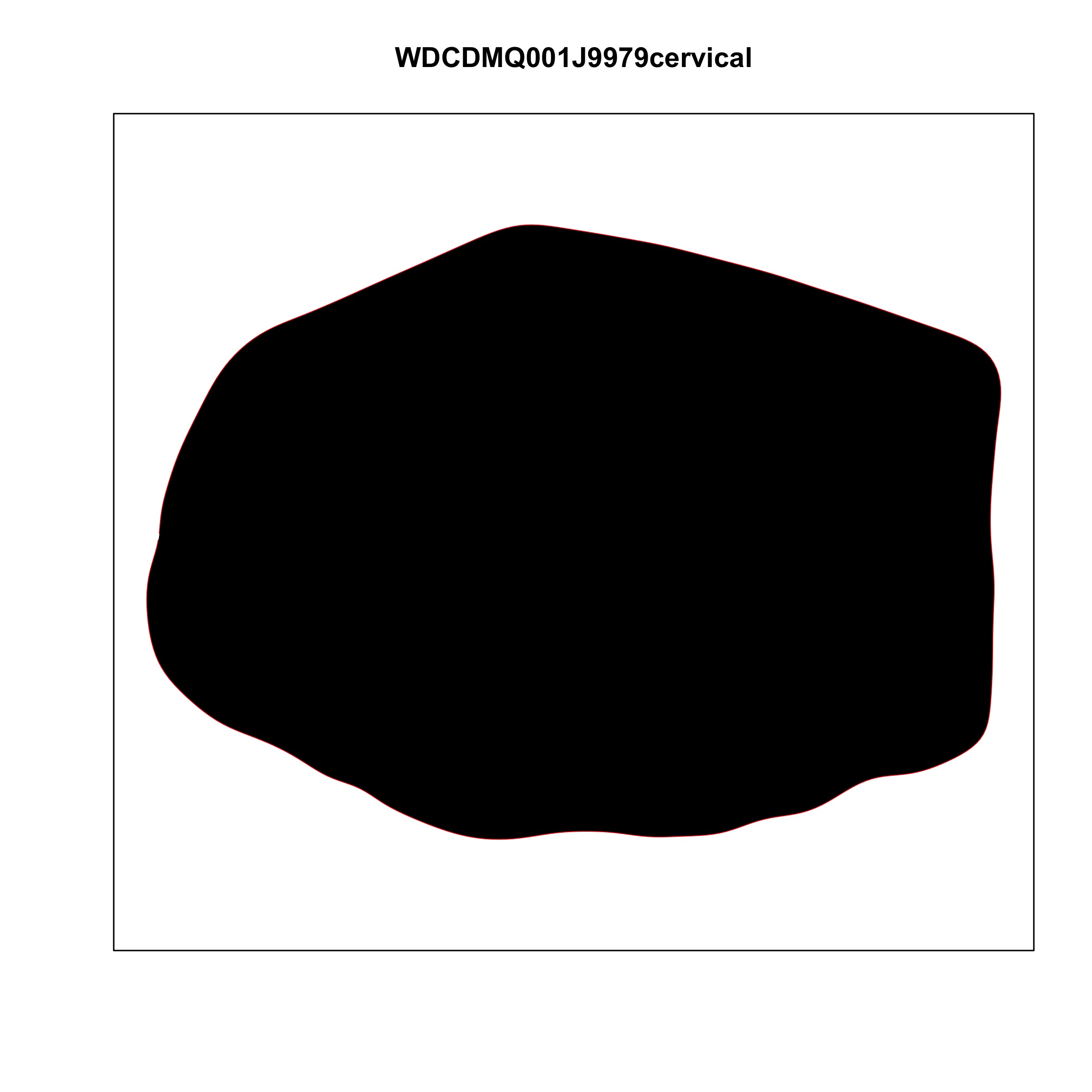

### plate_no_40.jpg

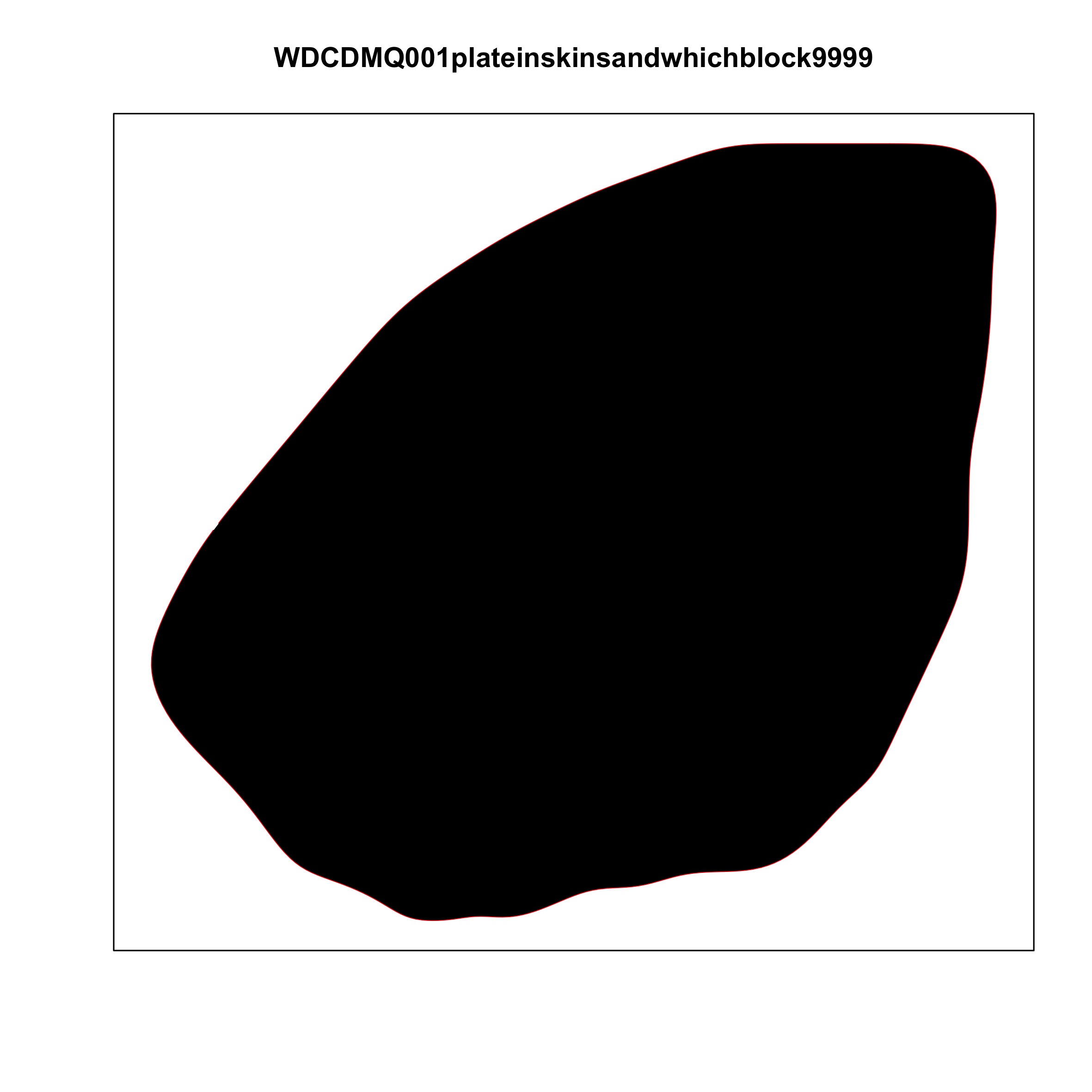
